# Mouse lemur transcriptomic atlas informs primate genes, mutations, physiology, and disease

**DOI:** 10.1101/2022.08.06.503035

**Authors:** The Tabula Microcebus Consortium, Camille Ezran, Shixuan Liu, Stephen Chang, Jingsi Ming, Lisbeth A. Guethlein, Michael F.Z. Wang, Roozbeh Dehghannasiri, Julia Olivieri, Hannah K. Frank, Alexander Tarashansky, Winston Koh, Qiuyu Jing, Olga Botvinnik, Jane Antony, Angela Oliveira Pisco, Jim Karkanias, Can Yang, James E. Ferrell, Scott D. Boyd, Peter Parham, Jonathan Z. Long, Bo Wang, Julia Salzman, Iwijn De Vlaminck, Angela Ruohao Wu, Stephen R. Quake, Mark A. Krasnow

## Abstract

Mouse lemurs (*Microcebus* spp.) are an emerging primate model organism. However, little is known about their genetics or cellular and molecular biology. In the accompanying paper, we used large-scale single cell RNA-sequencing of 27 organs and tissues to identify over 750 molecular cell types, characterize their full transcriptomic profiles, and study evolution of primate cell types. Here we use the atlas to characterize mouse lemur genes, mutations, physiology, and disease. We uncover thousands of previously unidentified lemur genes and hundreds of thousands of new splice junctions that globally define lemur gene structures and reveal over 85,000 primate splice junctions missing in mice. We systematically explore the lemur immune system, comparing the global expression profiles of key immune genes in health and disease, and molecular mapping of immune cell development, trafficking, and their local and global activation to infection. We characterize primate/lemur-specific physiology and disease including molecular features of the immune program, of lemur adipocytes that exhibit dramatic seasonal rhythms, and of metastatic endometrial cancer that resembles the human malignancy. We identify and describe the expression patterns of over 400 primate genes missing in mice, many with similar expression patterns in human and lemur and some implicated in human disease. Finally, we provide an experimental framework for reverse genetic analysis by identifying naturally-occurring nonsense (null) mutations in three primate genes missing in mice and analyzing their transcriptional phenotypes. This work establishes mouse lemur as a tractable primate model organism for genetic and molecular analysis, and it prioritizes primate genes, splice junctions, physiology, and disease for future study.

## INTRODUCTION

Although many of the genes, pathways, and principles of modern biology and the molecular foundations of medicine were uncovered in studies of the canonical genetic model organisms such as yeast, *Drosophila*, and mouse, new model organisms are being developed to study aspects of biology and medicine not observed or poorly modeled in the canonical model organisms^1–5^. The recent explosion of these emerging model organisms in diverse areas of biology has been fueled by advances in DNA sequencing and genomics, which has made construction of a high quality reference genome for a newly studied species readily attainable, and by gene editing tools like CRISPR-Cas9 systems that have made introduction of genes and targeted mutations in the genome practical in many species. It has remained challenging, however, to establish a rich cellular and molecular understanding of a new model organism and its biology, which has generally required decades to accrue through intensive effort of many researchers developing precision technologies and applying them to various aspects of the organism’s development, physiology, behavior, and diseases. We reasoned that single cell transcriptomics performed at organism-wide scale could greatly facilitate such an understanding, and in the accompanying manuscript (Tabula Microcebus)^6^ we created an extensive molecular cell atlas of the gray mouse lemur, *Microcebus murinus*.

Mouse lemurs provide an appealing model for the study of primate genes, biology and health. Practical advantages include their small size, easy lab husbandry, short generation time, and great abundance in nature among primates. Mouse lemurs are also favorably placed phylogenetically for a model organism, roughly half the evolutionary distance between mouse and human^7–9^. Their physiology has been studied for decades in laboratory colonies, especially their circadian and dramatic seasonal rhythms, metabolism, cognition and ageing^10–16^. Likewise, their ecology, behavior, and phylogeny have been investigated through field studies in their native Madagascar^17–21^. However, little has been known about their genetics, genomics, or cellular and molecular biology. That began to change with the publication of a reference genome sequence for *M. murinus*^*22*^. And, in the accompanying paper (Tabula Microcebus)^6^, we used large-scale droplet (10x) and plate-based (Smart-seq2, SS2) single cell RNA-sequencing (scRNA-seq) of ∼226,000 cells from 27 mouse lemur organs and tissues opportunistically procured from four aged donors that were clinically and histologically characterized^23^ (see clinical summary in Supplementary Results). This identified over 750 molecular cell types and their full transcriptomic profiles, and defined the molecular relationships of the cell types to each other and to the homologous cell types in human, mouse, and macaque. The reference genome and molecular cell atlas substantiate the close genetic proximity of mouse lemur to human, showing that human genes (Fig. S1), expression patterns, and cell types^6^ are each better modeled in mouse lemur than in mouse.

Here, we use this extensive molecular cell atlas to analyze primate genes, mutations, physiology, and disease. We find thousands of previously unidentified genes (Section 1) and hundreds of thousands of new splice junctions (Section 2) that globally define lemur gene structures and reveal over 85,000 primate splice junctions missing in mice, and we show how their cross-species expression profiles aid in gene annotation and characterizing gene evolution (Section 3). We use the atlas to probe the genes, cells, and local and global functions of the lemur immune system in development, health and disease (Section 4) and to characterize primate- and lemur-specific aspects of physiology and disease (Section 5). Finally, we describe the expression profiles of over 400 primate genes missing in mice (Section 6) and present an experimental framework for reverse genetic analysis by identifying naturally-occurring nonsense mutations in three such genes and analyzing their transcriptional phenotypes (Section 7). This work establishes mouse lemur as a tractable primate model organism for genetic and molecular analysis, and it prioritizes primate genes, splice forms, physiology and disease for future study. The approach can be readily applied to other emerging model organisms.

## RESULTS

### 1. Organism-wide scRNA-seq uncovers new genes

Our organism-wide scRNA-seq dataset provides a massive amount of transcriptomic sequence information: ∼2×10^12^ base pairs (∼10^12^ bp high-quality reads from 10x and SS2 each) distributed throughout the ∼2.5×10^9^ bp *M. murinus* genome, providing ∼10^4^-fold average coverage of the transcriptome (∼2.5×10^8^ bp of NCBI-annotated transcripts). We reasoned that such deep coverage of the transcriptome across most cell types of most organs would enhance gene detection, gene structure (intron-exon) definition^24^, and gene annotation beyond current state-of-the-art methods that rely primarily on phylogenetic sequence comparisons and, when available, bulk RNA sequencing, as done for the current *M. murinus* genome annotations (NCBI Microcebus Murinus Annotation Release 101, Ensembl Genome Browser v100). Here we used our scRNA-seq transcriptomic data in both a systematic approach to detect unannotated genes across the genome, and in a targeted approach to uncover genes in historically challenging loci.

For systematic gene detection, we used the hidden Markov model approach described in an associated paper^25^ to identify across the genome Transcriptionally Active Regions (TARs), locations with significant read coverage in our scRNA-seq datasets (Fig. 1a). This revealed that TARs comprise 13% (3.3×10^8^ bp) of the Mmur 3.0 genome assembly^22^, with most TARs (87%, 2.8×10^8^ bp, 11% of genome) mapping to previously annotated genes (aTARs) (Fig. 1b). The rest of the TARs (13%, 4.2×10^7^ bps, 1.7% of genome) mapped to unannotated regions (uTARs), suggesting they are genes with unassigned identities. uTARs that are differentially-expressed across cell types account for 2.4±1.5% (mean±s.d.) of the unique sequencing reads per cell, up to 18.5% in skin sweat gland cells (Fig. 1c, Table 1). These differentially-expressed uTARs are likely biologically significant because, in many tissues, from uTAR cellular expression patterns alone, we could distinguish cell types with a consistency that approached that using annotated genes or aTARs (Fig. S2a-c). Likewise, dimensional reduction of the testes scRNA-seq data using uTAR features alone reconstructed the full spermatogenesis program, mimicking the result with annotated genes^6^ (Fig. 1d-g, Fig. S2d).

**Figure 1.**
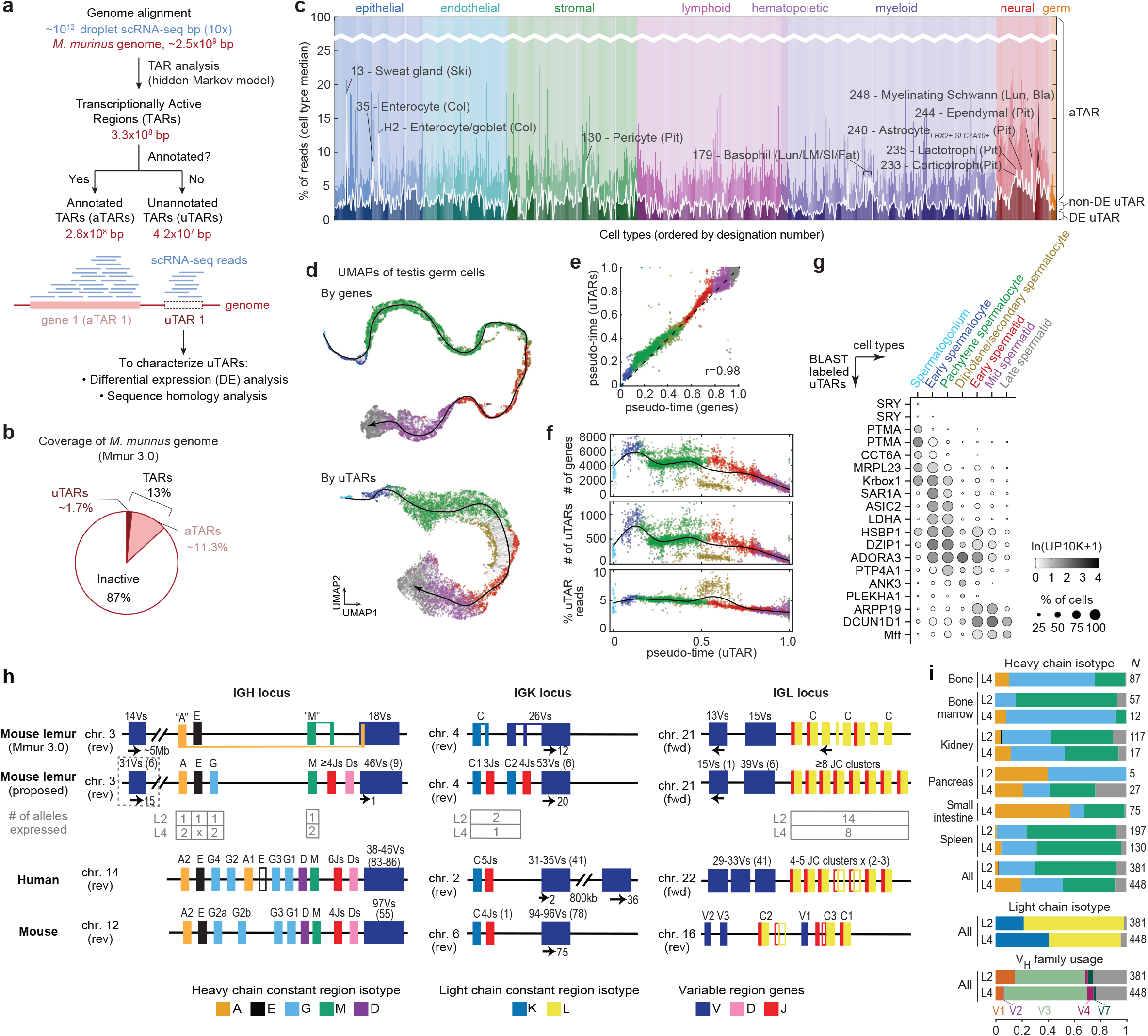
Organism-wide scRNA-seq uncovers new mouse lemur genes. **a**. Schematic of the analysis used to identify transcriptionally active regions (TARs) of the genome from 10x scRNA-seq data and to characterize the differentially-expressed, unannotated TARs (uTARs). **b**. Pie chart showing percent of the mouse lemur genome (base pairs) that is aTARs, uTARs, or transcriptionally inactive. **c**. Stacked bar plot showing for each atlas cell type the median percentage of transcript reads categorized as aTARs, differentially-expressed uTARs (DE uTAR), and non-differentially-expressed uTARs (non-DE uTAR). Cell types are ordered by designation number (see accompanying Tabula Microcebus manuscript^6^) and color coded by tissue compartment. Examples of cell types enriched for DE uTARs are indicated with their designation number and tissue of origin in parentheses. **d**. UMAP visualization of mouse lemur male germ cells from testis (L4, 10x dataset), with embedding based on expression of either the annotated genes^6^ (top) or uTARs alone (bottom). Each datapoint (cell) is color coded by its annotated stage of spermatogenesis (color code in panel g). Black lines with arrowheads indicate the pseudotime developmental trajectory of spermatogenesis, and fine gray lines show alignment of each data point to the trajectory. **e**. Comparison of male germ cell pseudotime trajectory coordinates based on expression of annotated genes (x-axis) or of uTARs (y-axis). Dashed black line, 1:1 relationship; *r*, Pearson’s correlation coefficient. **f**. Number of annotated genes (top) and uTARs (middle) expressed, and percent uTAR reads of total TARs (bottom), in each male germ cell, ordered along the uTAR-derived pseudotime trajectory, showing in uTARs the expected progressive transcriptional downregulation of genes during spermatogenesis. **g**. Dot plot showing average expression (ln(UMI_*g*_/UMI_total_ ^*^1e4 +1), abbreviated ln(UP10K+1), indicated by circle color) and percent of cells (circle size) expressing selected DE uTARs (names based on subsequently identified sequence homology) across the indicated male germ cell types ordered by developmental stage. Identical gene names indicate multiple uTARs aligned to the same gene in another species. **h**. Previously unannotated or incorrectly annotated immunoglobulin (Ig) genes identified by scRNA-seq of B and plasma cells. Schematics of mouse lemur (top), human^29,148^ (middle) and mouse^29^ (bottom) Ig loci for heavy chain (*IGH*, left) and kappa (*IGK*, center) and lambda (*IGL*, right) light chains, located on the forward (fwd) or reverse (rev) strand of the indicated chromosomes (chr) and colored as in key. Top lemur line shows annotation in Mmur 3.0’s NCBI Annotation Release 101; line below shows revised annotation. Filled boxes, constant (A, E, G, D, M for *IGH*, C for *IGK* and *IGL*), variable (V), joining (J) and diversity (D) regions; open boxes, pseudogenes. Above V regions are the estimated number of functional V genes (varies per individual) and in parentheses, the estimated number of V pseudogenes that lack transcripts. Gray dashed box, smaller V cluster ∼5Mb downstream of constant region in *IGH* locus which may be an assembly error (main V cluster is upstream). Arrows indicate genes oriented opposite to direction of constant regions, and those with numbers indicate subset of those genes that are flipped. Tables below mouse lemur locus indicate number of expressed alleles for each constant region isotype in each analyzed lemur. **i**. Barplot showing fraction of atlas B and plasma cells (SS2 dataset) by their expressed Ig heavy chain (top) and light chain isotype (middle), and heavy chain variable domain (V_H_) family member (bottom), colored as in panel h. Gray indicates unassigned isotype. N, number of cells analyzed. Data are shown separately for L2 and L4. Fractions for heavy chain isotypes are also shown separately for organs with ≥5 cells and reveal tissue specialization (e.g., *IGA*-expressing cells prominent in small intestine and pancreas). Note V_H_ gene families related to human *IGHV1, 3* and *4*, show the broadest expression, as in human and mouse^29,149,150^; however, light chain isotype *IGL* is more commonly expressed than *IGK*, in contrast to human and mouse B cells where *IGK* predominates^151^.

We sought to determine the gene identities of uTARs. We first showed that TAR analysis has high sensitivity for detecting previously annotated genes (Fig. S2e). For example, of the 5000 NCBI-annotated mouse lemur genes with the highest cell type expression variance in our 10x scRNA-seq dataset, TARs captured 98% (4884 genes), and of the 3904 genes annotated by Ensembl but not NCBI, TARs captured 44% (1728 genes, mostly non-protein coding genes, Table 1). TARs also captured 88% (376 genes) of the 425 primate-selective/mouse absent genes (Section 6, Table 8). We then searched for homologs in other species of the 4003 differentially-expressed uTARs (Table 1). DNA sequence (blastn) searches identified homologous genes for 2368 (59%) of these uTARs and transcript/protein sequence searches (nf-predictorthologs^26,27^) identified protein-coding (DIAMOND blastp) hits for 3185 (80%), noncoding (Infernal cmscan) hits for 45 (1%), and both protein-coding and noncoding hits for 231 (6%) (Fig. S2f). Some of these uTARs are conserved genes missed by the conventional annotation pipeline, such as dozens of genes present in human but missing from NCBI and Ensembl’s annotations of Mmur 3.0 (e.g., *GSTA3* in tendon cells of the bone, *TIGD1* in *CD4*+ T cells, *SPRR2G* in suprabasal epidermal cells, see Table 1).

We also used our scRNA-seq datasets to aid gene discovery in challenging loci. Immunoglobulin (Ig) genes are exceedingly difficult to annotate by conventional approaches because they comprise large arrays of related and rapidly evolving genes and gene segments, some extremely short (e.g. D-region segments are as short as 10 bp) but widely spaced in the genome, brought together by V(D)J recombination during B lymphocyte development to create antibody diversity^28^. Genomic mapping of Ig transcripts in the 829 B and plasma cells in our SS2 dataset revised *IGA* and *IGM* gene structures, uncovered D and J gene clusters, tripled the number of identified V genes (from 32 to 92), and identified 15 unexpressed V genes as likely pseudogenes (Fig. 1h, *IGH* locus). Some expressed V genes mapped to a large V gene cluster located ∼5 MB upstream of the rest of the *IGH* locus, suggesting it is an orphan V gene cluster^29,30^. This atlas-enhanced genome annotation shows that mouse lemur Ig heavy chain locus has a similar overall organization to that of human, but it is substantially streamlined with only a single constant region of each heavy chain isotype and no *IGD*, an evolutionarily plastic isotype lost in many lineages^31,32^. Hence mouse lemur provides a simplified model for understanding Ig gene rearrangement, expression, and functions. Similar analyses of Ig light chain genes corrected the structure of *IGK* and *IGL* constant regions, uncovered duplicated *IGK* J gene clusters and additional *IGL* JC clusters, and doubled the number of identified V genes at each locus (from 26 to 59 genes at *IGK*, 28 to 61 at *IGL*) (Fig. 1h, *IGK* and *IGL* loci). Thus, organism-wide scRNA-seq is a powerful way of detecting missed genes throughout the genome and at historically challenging regions like Ig loci.

### 2. Organism-wide scRNA-seq defines gene structures and splicing

To enhance gene structure and splice site definition, we used SICILIAN^33^ to uncover potential splice junctions from sequence reads that mapped with high confidence to discontinuous positions along the genome (Fig. 2a-b, Table 2). The current mouse lemur genome annotation (Mmur 3.0, NCBI Refseq Annotation Release 101, genome size ∼2.5×10^9^ bp) has 212,198 previously assigned splice junctions, 41% fewer than human (358,924 junctions in RefSeq hg38, genome size ∼3.1×10^9^ bp) and 33% fewer than mouse (319,497 junctions in RefSeq mm10, genome size ∼2.7×10^9^ bp), suggesting that thousands of lemur splice junctions remain to be discovered. Application of SICILIAN to our 10x and SS2 scRNA-Seq datasets computationally supported nearly all (98%, 202,802 junctions) of the currently annotated splice junctions; however, annotated junctions account for only 9.4% of SICILIAN-identified junctions (Category A in Fig. 2b-d). Newly-identified junctions include 67,672 junctions that had both 5’ and 3’ splice sites separately annotated previously (Category B, e.g., exon skipping) and 274,991 junctions between an annotated and a novel splice site (Category C); both types were supported by substantial reads (on average, 3,711 (Category B) and 346 (Category C) unique reads per junction across the dataset). SICILIAN also detected new junctions between two novel splice sites (Category D) and junctions that map to unannotated genes (Category E), but these were supported by fewer reads (98 and 121 reads/junction, respectively) suggesting some could result from natural “noise” in splicing^34,35^ or are highly cell-type specific.

**Figure 2.**
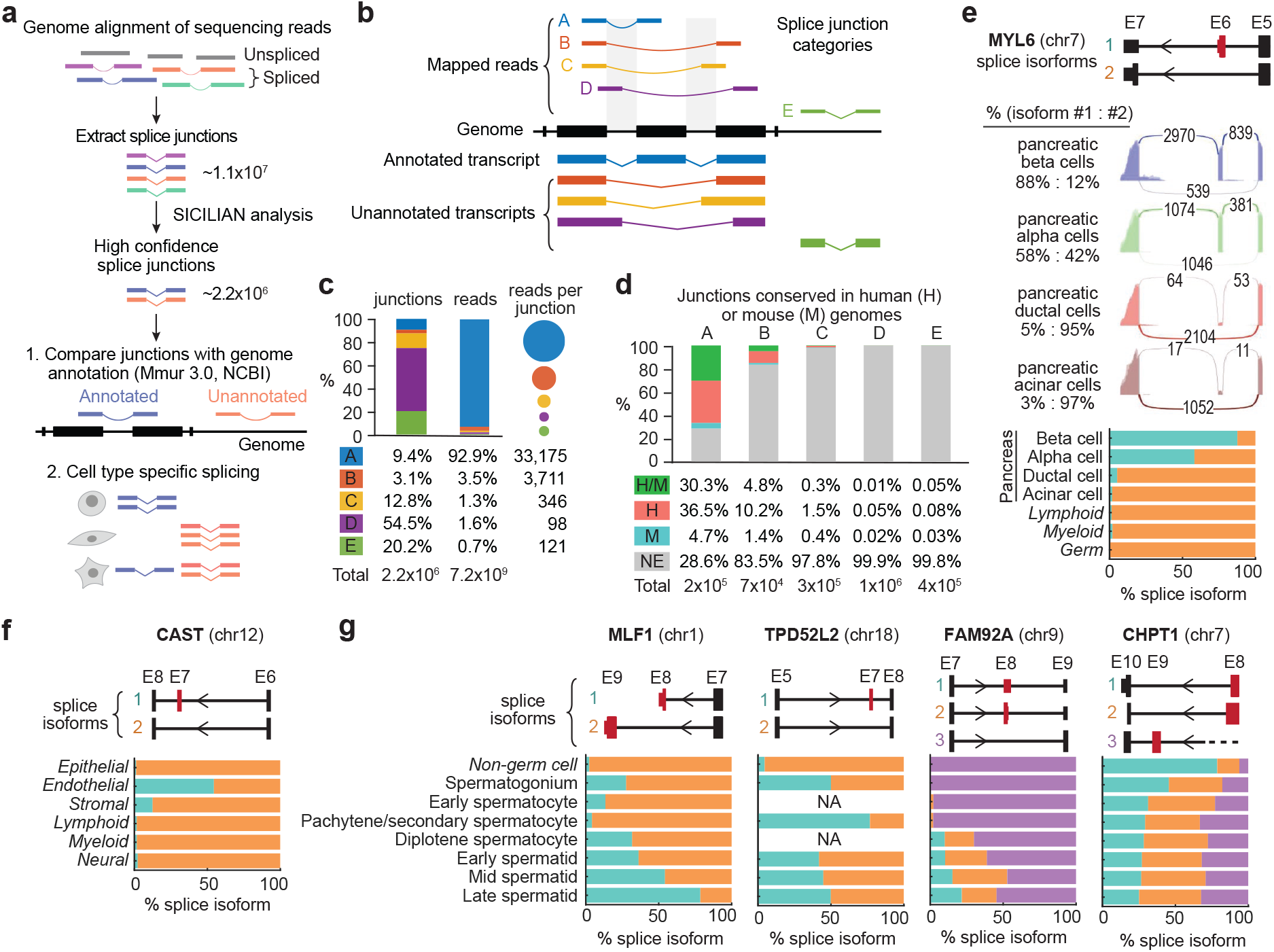
SICILIAN analysis of organism-wide scRNA-seq defines mouse lemur gene structures and splicing. **a**. Schematic of mRNA splicing SICILIAN analysis. Bars, exonic sequences; lines, intronic sequences **b**. Splice junction categories based on annotation status. Category A: Previously annotated splice junction. Category B: Novel junction between two annotated exon boundaries (e.g., novel exon skipping events). Category C: Novel junction between an annotated exon boundary and an unannotated location within the gene. Category D: Novel junction between two unannotated locations within a gene. Category E: Novel junction outside annotated gene boundaries. **c**. Percent of splice junctions, scRNA-seq reads, and average number of reads per junction for each junction category. **d**. Percent of splice junctions in each category that are conserved in both human and mouse genomes (HM), only in human (H), only in mouse (M), or neither (NE). **e**. Cell-type-specific alternative splicing of *MYL6*. Top, structure of *MYL6* transcripts (isoforms 1, 2) from exon 5 (E5) to exon 7 (E7) (NCBI annotation, detail in Fig. S3a; arrowhead, transcription direction 5’→ 3’); alternatively spliced exon 6 (E6) is red. Middle, mapped scRNA-seq read buildups for *MYL6* at E5 to E7 in indicated cell types; connecting arcs show number of sequence reads across the junction. Bottom, percent of isoform 1 (cyan) and isoform 2 (orange) in indicated cell types (combined 10x and SS2 scRNA-seq reads from all individuals); cell types across atlas are shown in Fig. S3a. **f**. Barplots as in panel e showing compartment-selective (endothelial, stromal unique) alternative splicing of *CAST*. Isoform 1 (cyan), exon 7 included; isoform 2 (orange), exon 7 excluded. Individual cell types in Fig. S3b. **g**. Barplots showing developmentally-regulated alternative splicing of *MLF1, TPD52L2, FAM92A*, and *CHPT1* during spermatogenesis and in comparison with atlas-wide non-germ cells. NA, no spliced transcript of gene in corresponding cell type. Dashed line, insufficient reads covering corresponding region of transcript. Individual cell types in Fig. S3c-f.

Over 85,000 of the lemur splice junctions are conserved in human but missing in mice (Fig. 2g, Table 2). Among the newly-detected junctions, nearly 19,000 are conserved in human and/or mouse, most of which (59%) belong to category B (Fig. 2d). These results suggest that organism-wide scRNA-seq combined with rigorous statistical methods such as SICILIAN robustly detect RNA splicing and gene structure in a newly characterized genome, and can be used to prioritize splice junctions and isoforms for further study, such as those that are present in primates but missing in mice.

We next identified cell type-specific splicing events through a MANOVA-based differential splicing analysis. This identified 545 genes that are most differentially spliced across cell types in any tissue (Table 3). For example, myosin light chain *MYL6*, a ubiquitously-expressed but poorly characterized gene^36^ can be alternatively spliced to either include or skip exon 6; in most cell types both isoforms are produced but the ratio of the two isoforms can differ dramatically (Fig. 2e, Fig. S3a). In pancreatic β and α cells, most transcripts include exon 6, whereas in ductal and acinar cells, as well as most immune and germ cells, almost all transcripts exclude it. *CAST*, a regulator of membrane fusion, also exhibits differential splicing, with exon 7 included in about 50% of transcripts in endothelial cell types but almost always skipped in other cells (Fig. 2f, Fig. S3b). In addition, numerous genes including *MLF1, TPD52L2, FAM92A*, and *CHPT1* show sperm-specific splicing patterns and are differentially spliced during spermatogenesis (Fig. 2g, Fig. S3c-f).

### 3. Organism-wide scRNA-seq aids gene annotation

Gene identity assignments in newly sequenced genomes have traditionally relied on phylogenetic sequence comparisons and chromosomal position, with each gene assigned a name corresponding to that of the characterized homolog in other species with greatest sequence similarity and conserved gene order along the chromosome, indicating a direct evolutionary relationship (ortholog). However, such analyses can be ambiguous, returning no significant homologies or uncovering multiple homologs with similar levels of sequence identity and evolutionary relatedness, obscuring the true ortholog^37^. Hence, about a quarter of the genes (∼7,600) in the current mouse lemur NCBI genome annotation (assembly Mmur 3.0, NCBI Refseq Annotation Release 101) have been assigned only a locus identifier (e.g. “Loc “ or “orf”) and no formal gene name/symbol or description (“uncharacterized genes”), and another quarter (∼8,000) have been assigned a locus identifier and initial description based on sequence homology but no gene name/symbol (“unnamed genes”) (Fig. 3a). The fractions of these unnamed and uncharacterized genes in the current lemur genome annotation are much greater than those of human and mouse. We reasoned that organism-wide gene expression profiles could complement the classical approaches to gene orthology assignment and naming by identifying the sequence homolog(s) with the most conserved expression pattern (the “expression homolog”).

**Figure 3.**
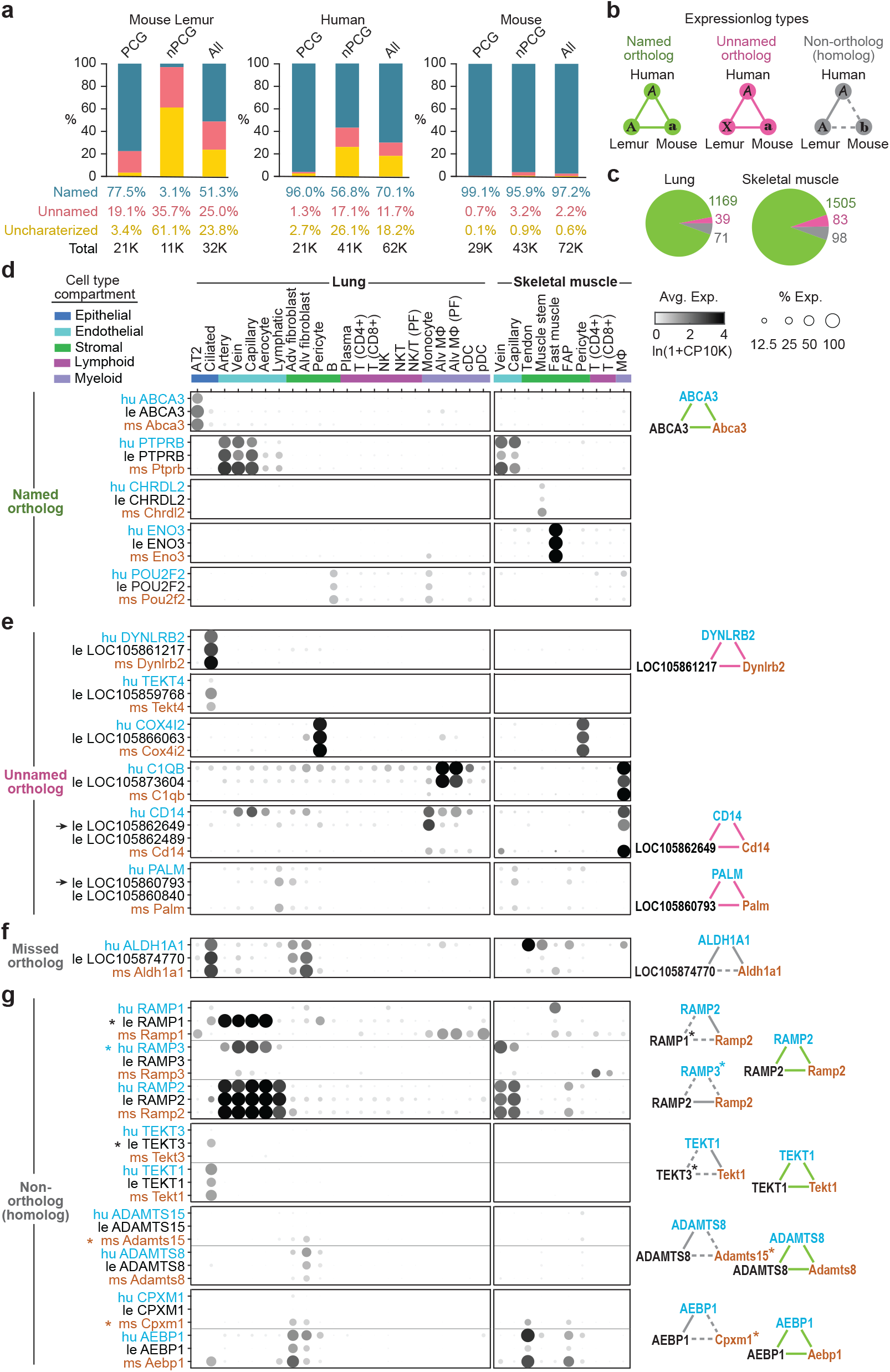
SAMap analysis of atlas scRNA-seq data aids mouse lemur gene annotation. **a**. Percent of genes that are named (with gene symbol), unnamed (only a locus identifier, e.g., “Loc “ or “orf”, and a suggested gene description), and uncharacterized (unnamed and with no gene description) in mouse lemur (left; genome assembly Mmur 3.0, NCBI annotation release 101), human (middle; assembly GRCh38.p13, NCBI annotation release 109), and mouse (right; assembly GRCm38.p6, NCBI annotation release 108), shown for all genes in genome (all) or for protein-coding genes (PCG) or non-protein-coding genes (nPCG). **b**. Schematic of human-lemur-mouse expression homolog triad types based on similarity of their cell type specific expression profiles, genome-assigned orthology (according to NCBI and Ensembl ortholog databases), and naming status of mouse lemur gene. Solid line, expression homologs with previously assigned gene orthology; dashed line, expression homologs without assigned orthology. Left (green): expression homolog triad involving assigned orthologs in all three species, and lemur ortholog is named accordingly. Middle (pink): expression homolog triad involving assigned orthologs in all three species, but lemur ortholog remained unnamed. Right (gray): expression homolog triad involving homologous (calculated by sequence homology^39^) genes that are currently not assigned orthologs for at least one of the species, regardless of the naming status of lemur gene. **c**. Pie charts showing percent of named ortholog (green), unnamed ortholog (pink), or non-ortholog (gray) expression homolog triads identified in the lung and skeletal muscle scRNA-seq datasets, respectively. **d**. Dot plot showing average expression for five selected expression homolog triads of the “named ortholog” type across human, lemur, and mouse lung and skeletal muscle cell types. In each triad (boxed), human gene ortholog (blue) has “hu” prefix, lemur (black) has “le” prefix, and mouse (orange) has “ms” prefix. Cell types are ordered by compartment; Adv (adventitial), Alv (alveolar), MΦ (macrophage), cDC (conventional dendritic cell), pDC (plasmacytoid dendritic cell), FAP (fibroadipogenic progenitor cell), PF (proliferating). Diagram (right) shows an example expression homolog triad (format of panel b; line color indicates expression homolog type). **e**. Selected expression homolog triads of “unnamed ortholog” type. Note lemur genes have only locus identifiers and no gene symbol. In bottom two examples (*CD14, PALM*), two lemur orthologous genes with identical gene description were assigned by NCBI (i.e., monocyte differentiation antigen CD14-like for *CD14*, paralemmin-1 for *PALM*), but only one is detected as an expression homolog (arrows), suggesting it is the true ortholog. **f**. Selected expression homolog triad of “non-ortholog” type that likely represents a missed orthology assignment. Note lemur gene is assigned as an ortholog of human *ALDH1A1* but not of mouse *Aldh1a1* in NCBI or Ensembl. **g**. Selected expression homolog triads of “non-ortholog” type. In each example, all relevant expression homolog triads are shown (including corresponding “named ortholog” triads); asterisk indicates the outlier non-orthologous gene. For example, three *RAMP* expression homolog triads are detected: two “non-orthologs” (i.e., hu *RAMP2* - le *RAMP1*^*^ - ms *Ramp2*, hu *RAMP3*^*^ - le *RAMP2* - ms *Ramp2*) and one “named ortholog” (hu *RAMP2* - le *RAMP2* - ms *Ramp2*). The shared expression pattern of lemur *RAMP1* and human *RAMP3* with *RAMP2* suggests that lemur RAMP1 and human RAMP3 have evolved to engage in similar physiological functions as the species-conserved RAMP2 or instead to modulate RAMP2-mediated ligand signaling in lung endothelial cells.

We used SAMap^38,39^ to find, for each mouse lemur gene, the mouse and human homologs with the most similar patterns of cell type specific expression across 32 orthologous cell types in lung and muscle, carefully curated in the same way for all three species. This identified 1,279 expression homolog triads in lung and 1,686 in muscle (Table 4), most of which (91% and 89% in lung and muscle respectively) are triads of named orthologous genes across the three species, substantiating their assignments as orthologs by traditional approaches (e.g., *ABCA3* is selectively expressed in lung AT2 cells from all three species) (Fig. 3b-d, Fig. S4). In addition, we identified 39 (3%) lung and 83 (5%) muscle orthologous gene triads that show conserved expression patterns but for which the lemur locus remains unnamed by NCBI, strongly suggesting that the identified lemur gene is the true ortholog and should be named accordingly (Fig. 3b-c, e). This also includes instances where multiple unnamed lemur loci are assigned the same gene description by NCBI, ambiguity likely resulting from the traditional annotation pipelines relying principally on sequence homology. For example, both lemur loci *LOC105862649* and *LOC105862489* share the description “monocyte differentiation antigen CD14-like” in NCBI. By comparing their expression patterns across species, we identify *LOC105862649* as the likely lemur *CD14*, which is expressed in lemur myeloid cells similar to human and mouse *CD14* (Fig. 3e). In contrast *LOC105862489* is sparsely expressed (<10 cells in the atlas) so may be a pseudogene. In other instances, we identified expression homolog triads with incomplete orthology assignments such as lemur locus *LOC105874770* (Fig. 3f), which shares expression in lung ciliated and fibroblast cells with the human ortholog *ALDH1A1* as well as the mouse homolog *Aldh1a1*, though the mouse gene has not been assigned as an ortholog to the lemur gene. The similar expression pattern between mouse and lemur provides strong evidence that these two genes are orthologs currently missed by standard annotation pipelines.

Surprisingly, the analysis also identified a small fraction of genes (71/1279 (6%) in lung, 98/1686 (6%) in muscle) whose expression patterns were not conserved with their assigned orthologs (Fig. 3b-c, g). For example, *RAMP1* is highly expressed in lemur lung endothelial cells, however the gene is only sparsely expressed in human lung and is expressed in mouse myeloid cells instead. When compared to its homologs, we found that lemur *RAMP1* shares a lung expression pattern most similar to *RAMP2*, which is selectively expressed in endothelial cells across all three species. This suggests that lung endothelial and myeloid cells have species-specific responses to hormones including CGRP and adrenomedullin through the RAMP/CALCRL receptor complexes^40–42^. Similarly, we identified such expression pattern changes for human *RAMP3* (top expression homolog being *RAMP2*), lemur *TEKT*3 (expression homolog *TEKT1*), mouse *Adamts15* (expression homolog *ADAMTS8*), and mouse *Cpxm1* (expression homolog *AEBP1*) (Fig. 3g). These exemplify rare instances where species-specific adaptations have dissociated gene expression patterns from their conserved sequence/structure^6^. Examining expression homologs thus provides another dimension for orthology assignment, gene naming, and exploring evolutionary diversification in expression and function, which can be further enhanced by expanding the analysis to additional organs and species.

We also used the organism-wide atlas to enhance annotation of the major histocompatibility complex (MHC) (Fig. S5a-e), encoding antigen-presenting proteins critical for vertebrate adaptive immunity^43^. The MHC is extremely difficult to annotate because of its evolutionary plasticity^44–48^, with some genes among the most polymorphic in the genome^49^ due to mutations and gene duplications/deletions that individualize immune systems and their response to infection. Allele-specific expression analysis of MHC class II genes across the atlas established gene copy number and distinguished major (highly and broadly expressed) class II genes (*DQA, DQB, DRA*, and *DRB*) from minor genes expressed at lower levels and in fewer cells (*DMA, DMB, DPA, DPB*) and putative pseudogenes not expressed at all (*DOA, DOB*) (Fig. S5c, h). Similar analysis of MHC class I genes distinguished a cluster of non-expressed pseudogenes (chromosome 6) from a functional cluster of 11 expressed genes on chromosome 20q that includes four with high and widespread expression we designate “classical” (*Mimu-168, -W03, -W04, -249*) and three previously thought to be pseudogenes (*Mimu-180ps, -229ps, - 239ps*) based on sequence analysis (Fig. 4a-c, h). Additional details are provided in Supplementary Results and Guethlein et al^50^.

**Figure 4.**
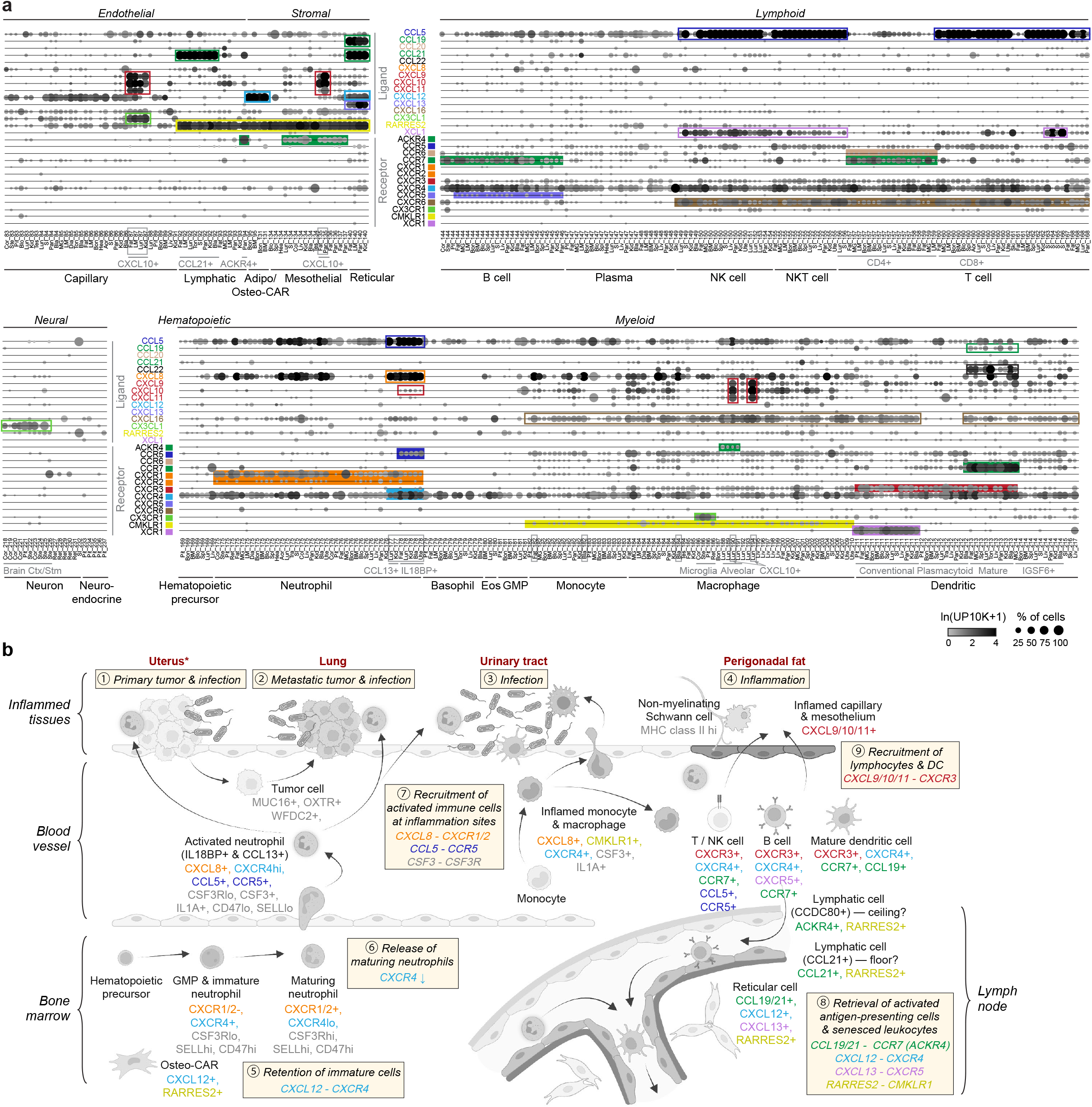
Mapping expression of chemokine signaling and systemic immune responses. **a**. Dot plot of average expression of selected chemokine receptors and their primary cognate ligands across immune cell types and other major interacting cell types in the atlas (10x dataset). (Figs. S6c shows epithelial cell types and all genes analyzed, and Table 6 provides the full list of gene interactions). Filled boxes highlight representative cell types highly expressing a receptor, and open boxes of the same color highlight representative cell types highly expressing cognate ligand. Note cell-type-stereotypical expression of many chemokine ligands and/or receptors, which guides specific immune cell trafficking under normal physiology (see Section 4 and Supplementary Results), as well as inflammation and disease related expression in several cell types (boxed in gray and examples illustrated in panel b). **b**. Schematic summary of multi-organ inflammatory processes, exemplified in L2, who was diagnosed with endometrial cancer (1) with metastatic spread to lung (2) (see Section 5) and secondary bacterial infection in both organs. L2 also exhibited suppurative cystitis (3) and suspected inflammation in perigonadal fat (4). We detected accompanying local and systemic immune responses, and the likely underlying genes and immune signals (indicated as *ligand — receptor* and colored as in panel a). These include a systematic increase in circulating maturing neutrophils (see Section 4 and Fig. 5d). These maturing/immature cells and other immune progenitors (e.g., granulocyte-monocyte progenitors (GMP)) are normally retained in the bone marrow by surrounding niche cells (e.g., osteo-CAR cells) through CXCL12-CXCR4 signaling^101,152,153^ (5). Upon infection, maturing neutrophils are released from the bone marrow, presumably through down regulation of *CXCR4* and upregulation of *CSF3R, CXCR1/2*^*55,56,154*^ (6). At the site of infection, two subtypes of activated neutrophils (designated as *IL18BP*+ and *CCL13*+ respectively) were found expressing high levels of chemoattractants that can recruit additional immune cells (7) (see Fig. 5b-d, Fig. S8e-f, Supplementary Results). These activated neutrophils also express markers of senescence including *CXCR4* and decreased levels of the “don’t eat me signal” (*CD47*), facilitating their trafficking to lymph node, bone marrow, and spleen for antigen presentation and/or clearance^71,72^, guided by subtypes of lymphatic endothelial and reticular stromal cells expressing *RARRES2, CCL21*, and/or *ACKR4* (8) (see panel a). In the bladder and perigonadal fat, inflamed/activated monocytes and macrophages were found to express high levels of cytokines *CSF3, CXCL8*, and *IL1A* (7) (see panel a, Fig. S9d, Supplementary Results). In perigonadal fat, inflamed capillary and mesothelial cells were identified expressing interferon-γ induced *CXCL9/10/11* which attract *CXCR3*-expressing immune cells^155,156^ (9). Non-myelinating Schwann cells in perigonadal fat also expressed elevated levels of MHC class II genes (see Fig. S5h), suggesting an inflammation state^157^ (9). Schematic created with BioRender.com.

Thus, organism-wide scRNA-seq is a valuable means of refining gene annotation and orthology assignments. This utility in gene annotation as well as in gene (Section 1) and splice site (Section 2) discovery provide a powerful complement to phylogenetic sequence comparison in creating the highest quality reference genome for a new model organism.

### 4. Mapping immune gene expression, development and function across the body

Very little is known about the cell or molecular biology of lemurs, but the organism-wide transcriptomic atlas greatly expedites a cellular and molecular understanding of its physiological processes. Here we show how the atlas illuminates lemur immune function, a critical biomedical process with significant differences between human and mouse^51^, and the accompanying manuscript illustrates this for endocrinology^42^. We first mapped global expression patterns of three key sets of immune genes (MHC, Ig/B cell receptors, chemokines), and then examined immune cells across the body to characterize their development, dispersion, and activation. These analyses supported general immune action and functions in the lemur but also revealed unexpected and primate-specific features of immune specialization.

Classical class I MHC genes were highly and broadly expressed (Fig. S5f-h), supporting a widespread role in presenting peptides derived from cytosolic proteins to *CD8+* T cells^52^. But expression varied substantially between compartments (highest in endothelial and immune, intermediate in stromal and epithelial, low in neural and germ compartments), and even within a compartment there were significant differences among cell types (Fig. S5f-h). For example, *CXCL10+* capillary cells and lung capillary aerocytes showed the highest class I gene expression in the entire atlas, and non-myelinating Schwann cells were a striking exception to the generally low expression across the neural compartment, suggesting these cell types play special immune roles protecting the lung and peripheral nervous system against intracellular pathogens. Class II MHC genes were more specifically expressed, notably in “professional” antigen-presenting cells (dendritic cells, macrophages, B cells) (Fig. S5f, h), supporting their role in presenting fragments of engulfed extracellular pathogens to helper (*CD4+*) T lymphocytes^53^. But they were also expressed across the endothelial compartment of all tissues, similar to humans but not rodents^54^, and at especially high levels in several specialized capillary subtypes (Fig. S5f, h). There was little expression in the stromal, epithelial, and neural compartments, with the notable exception of two stem cell niche cells (adipo-CXCL12-abundant reticular (CAR), osteo-CAR cells), and some mesothelial and lung epithelial (ciliated, AT2) cells (Fig. S5h). These high-expressing non-immune cell types presumably play “non-professional” roles in alerting the immune system to extracellular pathogens.

Mapping expression of the identified Ig/B cell receptor genes across the atlas established many classical features of B cell development and function in lemurs. These include expression in each B cell of a dominant Ig heavy chain and light chain isotype (Table 5, Fig. 1i), heavy chain class switching during B cell development with clonal expansion and distribution to different organs (Fig. S6a-b), heavy chain isotype tissue specialization (Fig. 1i), and definition and characterization of the complementarity-determining antigen binding region of the heavy chain (CDRH3) (Fig. S6a).

Expression of the chemokines receptors (24 genes) and their cognate ligands (32 genes) (Table 6) provides insight into the control of immune cell trafficking in health and disease (Fig. 4a-b, Fig. S7a-f). While the ligands were broadly expressed across non-germ compartments, the receptors were mostly restricted to the immune populations (Fig. S7a-b). We identified specific cell types (adipo-CAR, osteo-CAR cells) that expressed chemokine *CXCL12* so may retain hematopoietic progenitors (which express its receptor *CXCR4*) within bone marrow and regulate release into circulation of maturing immune cells as they downregulate receptor expression^55,56^ (Fig. 4a-b, Fig. S7d). We also identified epithelial cell types of the skin, GI tract, and bladder that express chemokine *CCL20* so can recruit receptor *CCR6*-expressing immune cells (Fig. S7c, Fig. 4a), as reported in skin of psoriasis patients^57,58^; cortical and brainstem neurons that express *CX3CL1* so could recruit *CX3CR1*-expressing microglia to the brain^59,60^ (Fig. 4a); and cell types and chemokines that may traffic diverse immune cells to lymph nodes (see Supplementary Results).

In addition to these local immune interactions, we were able to globally map the mouse lemur hematopoietic program from progenitors in bone marrow through their maturation, dispersal and differentiation throughout the organism, plus their activation in specific target tissues. The integrated lemur immune cell UMAP (Fig. 5a, Fig. S8a-d) reconstructs the developmental trajectories of major hematopoietic lineages. Below we molecularly dissect the neutrophil lineage.

**Figure 5.**
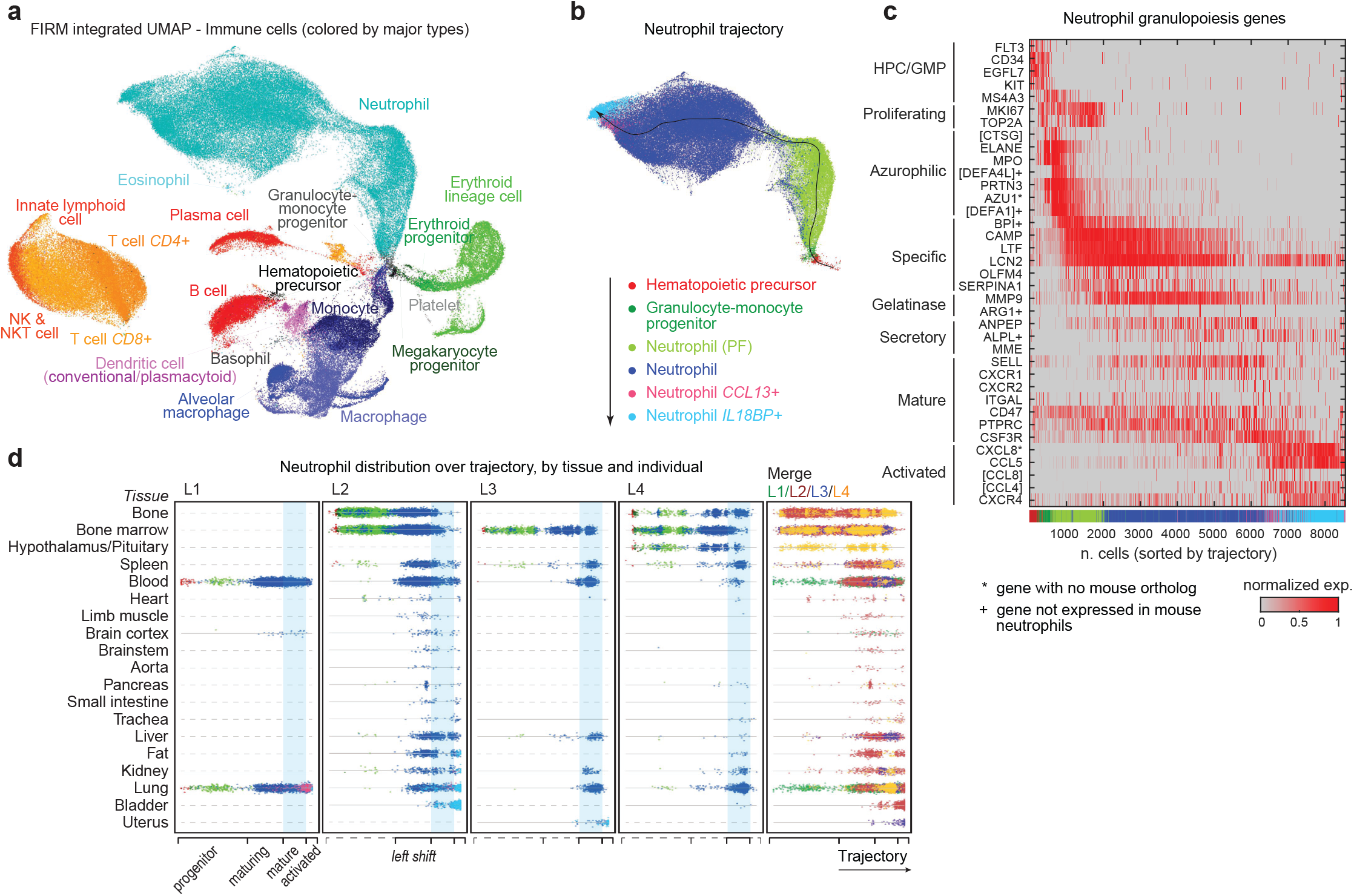
Organism-wide mapping of immune cell development and activation. **a**. Integrated UMAP of atlas immune cells (10x and SS2 datasets from all tissues and individuals, integrated by FIRM) with cells (dots) positioned by their overall transcriptional similarity and color coded by major immune cell type. Same UMAP highlighting proliferation state, sequencing method, individual, and tissue of origin is shown in Fig. S8a-d. **b**. Neutrophil trajectory as in panel a with cells colored by indicated neutrophil progenitor/cell type designation. Black line, pseudotime trajectory, with arrow indicating maturation direction, determined by expression of neutrophil markers; thin gray lines, individual cell alignments to trajectory. **c**. Heatmap showing relative expression (normalized to stable maximum (99.5 percentile)) of orthologs of human neutrophil markers in lemur neutrophil lineage cells (10x dataset). Cells (columns) are ordered by pseudotime trajectory in b, from progenitors (left) to (activated neutrophils (right) (see bar at top colored by cell type designation). Because non-activated neutrophils are highly abundant, only 10% are shown (uniformly subsampled along trajectory) for visualization purposes. Note the human-like sequential expression of these markers including granulopoiesis genes; azurophilic (primary) granules (*AZU1, MPO, ELANE*) in early stages, followed by specific (secondary) granules (*LTF, CAMP, LCN2*), then gelatinase granules (*MMP9, ARG1*), and finally the secretory vesicles (*ALPL, MME*) in mature neutrophils. ^*^, marker genes without a mouse ortholog; +, marker genes known to not be expressed in mouse neutrophils. [], description of gene identified by NCBI as a gene locus ([CTSG], LOC105866609; [DEFA4L], LOC105881499; [DEFA1], LOC105881500; [CCL8], LOC105885739; [CCL4], LOC105881712). **d**. Partitioning of neutrophil lineage cells along pseudotime trajectory (x-axis) by tissue of origin (y-axis) and mouse lemur individual (L1-L4, merged in right panel and colored by individual). Cells (dots) are colored as in b. Light blue stripes, approximate trajectory range of non-activated, mature neutrophils, the main circulating (blood) neutrophil population in health. Note L1 had neutrophil progenitors and immature cells in circulation (blood), indicating dysregulation of granulopoiesis; L2 had maturing neutrophils in blood (clinically known as a “left shift”); and L1-3 all had activated neutrophils in peripheral inflamed tissues (also see Fig. S8e-f), likely in response to infection and malignancy. Gray dashed lines, organs not sequenced in the individual.

Neutrophils are an abundant class of circulating leukocytes that ingest microbes and release granules containing enzymes that kill them. We used human/mouse neutrophil markers (*CSF3R+, MSR1-*) to identify ∼59,000 developing, proliferating, and mature lemur neutrophils across the atlas and recapitulate their full trajectory (Fig. 5b). This revealed sequential expression of granulopoiesis genes (Fig. 5c), mimicking the time course of different granule production at each stage of human neutrophil maturation^61,62^. This includes antimicrobial enzymes absent or lowly expressed in the corresponding mouse granules, such as defensins (*DEFA1, DEFA4*), bactericidal/permeability increasing protein (*BPI*), alkaline phosphatase (*ALPL*), and arginase (*ARG1*)^63^. Lemur neutrophils also express multiple human neutrophil genes (*AZU1, IL32, TCN1, FCAR* (*LOC105877181*), *S100A12*, and chemokines *CCL14, CCL16, CXCL8*) not present in the mouse genome^63–65^ (Fig. 5c, Fig. S7f, and see below Fig. 7f, Fig. S12i).

Mapping the tissue locations of neutrophils along the trajectory in each profiled individual revealed local inflammatory sites of neutrophil activation and turnover, along with global feedback regulation of the hematopoietic program (Fig. 5d). The expected distribution of neutrophils in health is earliest progenitors (myeloblasts, promyelocytes, myelocytes, metamyelocytes) and maturing (band) neutrophils predominantly localized to bone marrow, with blood and all other tissues enriched for mature, unactivated neutrophils, as observed for lemur 4 (L4). However, activated neutrophils, at the extreme end of the developmental trajectory, were found in L2’s lung, bladder, kidney, and perigonadal fat, as well as L1’s lung and L3’s uterus (Fig. 5d, Fig. S8e-f), focal sites of inflammation associated with infection and/or malignancy as confirmed by histopathology (Fig. 4b, Section 5, and Supplementary Results). These advanced neutrophils showed downregulation of mature neutrophil markers (e.g., *SELL/CD62L, MMP9, CXCR1, ITGAL*) that facilitate extravasation, and induction of chemokines that promote homing to sites of inflammation and recruitment of additional neutrophils (*CXCL8/IL8, CCL5/RANTES*)^66–70^ (Fig. 5c, Fig. S8f). Some of these genes are markers of neutrophil aging and trafficking to lymph node (increased *CXCR4*, decreased *SELL* and *CD47* (“don’t eat me” signal)), suggesting antigen-presentation and/or clearance following activation^71,72^ (Fig. 5c, Fig. 4b). Subclustering of these activated neutrophils revealed two subpopulations (*CCL13+* and *IL18BP+*), with differing tissue distributions across lemur 1-3 (Fig. S8e-f, Fig. 5d, Supplementary Results), suggesting that local factors can drive distinct neutrophil activation pathways. We also uncovered global responses to neutrophil activation. L2 had a leukocytosis (32.1 K/ul, normal range 4.5-11 K/uL in humans) dominated by neutrophils (91%), which were shifted toward more immature stages of the developmental trajectory (Fig. 5d), a molecular demonstration of the classical “left shift” seen in human blood smears when immature/maturing neutrophils of bone marrow are released into circulation to replenish the mature neutrophils recruited to an infection site^73^ (Fig. 4b). L1 also showed a distinctive global neutrophil pattern, with neutrophils from across the developmental trajectory including early progenitors in circulation (Fig. 5d), presumably from dysregulation of granulopoiesis by widespread fibrous osteodystrophy in his skeletal bones identified by histopathology. These global and local changes mirror classical human inflammatory responses, and shed light on the molecular signaling regulating primate neutrophil activation and renewal.

We similarly mapped development and trafficking across the body of the monocyte/macrophage lineage, which showed dozens of distinct, tissue-specific macrophage subtypes, including many novel subtypes and several locally activated monocyte/macrophage subtypes (Fig. S9, Supplementary Results). In contrast, mature T, NK, NKT, and innate lymphoid cells formed a single isolated cluster, belying their well established functional distinctions; similar patterns were found for B cells and plasma cells (Fig. 5a, Fig. S8g). This suggests lymphocyte development is rapid with few standing intermediates, or that tissues containing such intermediates were not among those profiled.

Thus, systematic examination of organism-wide immune cell distributions and expression patterns provides a rich and dynamic portrait of the mouse lemur immune system, revealing many cellular and molecular aspects of its development and function including human-like features differing from mouse, such as the presence and expression pattern of a dozen critical neutrophil genes (e.g., granule enzymes and chemokines) and of MHC class II genes in endothelial cells. These results support the potential of mouse lemur as a primate model for immunology and infectious diseases.

### 5. Cellular and molecular characterization of primate diseases and physiology

One reason for establishing mouse lemurs as a model organism is for investigations of primate biology and disease. The lemurs analyzed here were elderly and had spontaneous human-like pathologies revealed by full necropsies^23^. For example, both females (L2, L3) had endometrial cancer (Fig. 6a-e), the most common malignancy of the female reproductive tract and fourth most common cancer in U.S. women^74^, with increasing incidence and mortality over the past decade attributed to an aging population and increasing obesity^75,76^. Type 1 endometrial carcinomas are of low-grade endometrioid histology and typically estrogen-induced with a favorable prognosis under standard therapy, whereas type 2 tumors are of high-grade endometrioid, serous or clear cell histology, commonly estrogen-independent and lack durable therapies^77,78^. Animal models are limited: mice do not naturally acquire endometrial cancer, and although rats do, they and the engineered mouse models generally resemble type 1 rather than the high-grade, intractable type 2 tumors^78^. The spontaneous mouse lemur tumors were uncovered in L2 as a novel lung epithelial cell type (Fig. 6f) that curiously expressed high levels of oxytocin receptor (*OXTR*) (Fig. S10), a gene known to be highly expressed in female reproductive tissues and brain^79,80^. Comparison to cells across the atlas revealed their similarity to uterine epithelial cells (Fig. 6g-i, Fig. S10); necropsies established the diagnosis of primary uterine endometrial carcinoma with lung metastases (L2) or with local spread to mesenteric lymph nodes (L3)^23^. Organism-wide atlases thus enable identification of the primary site of cancers of unknown origin, which comprise ∼2% of all human cancers^81^.

**Figure 6.**
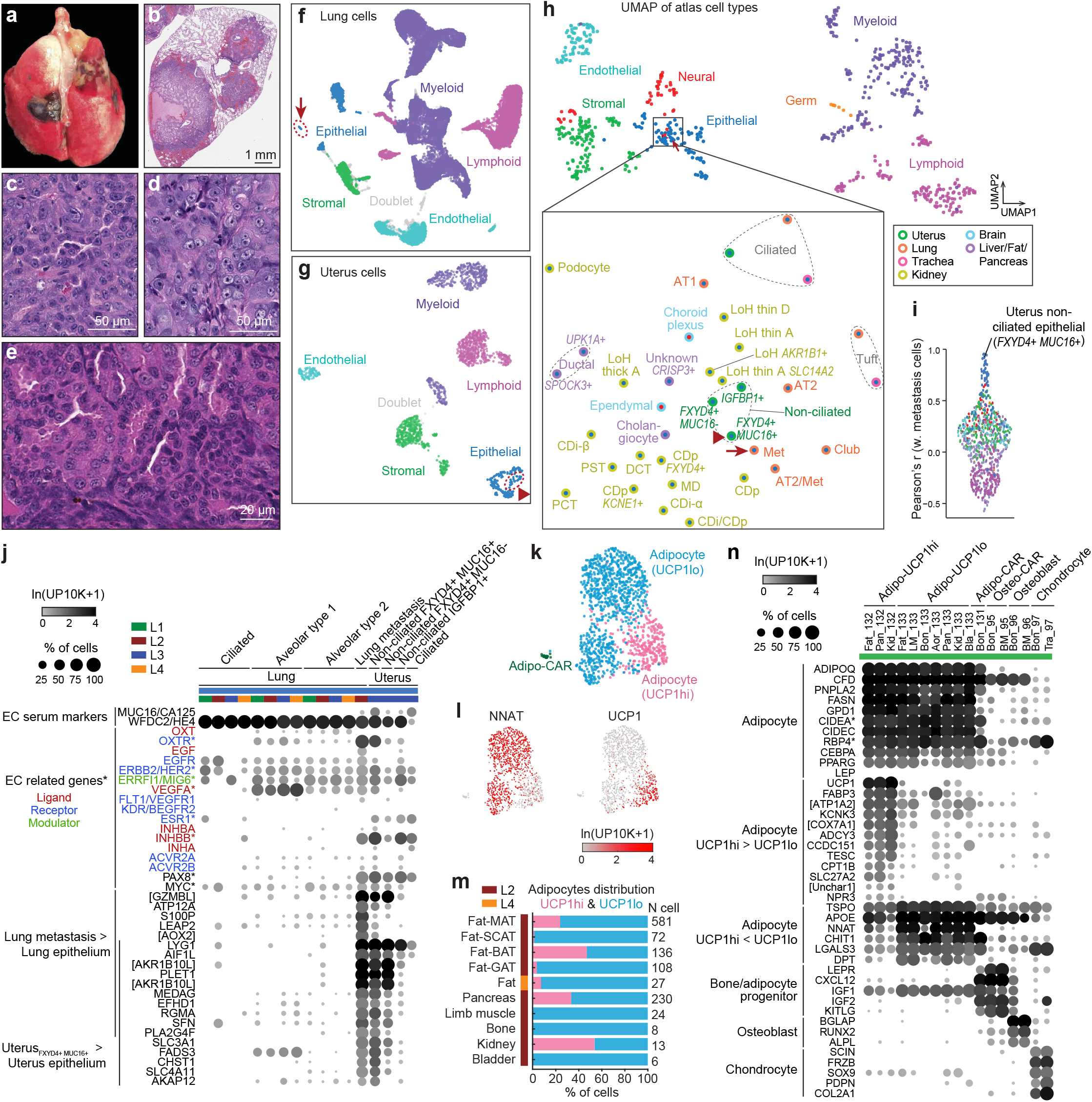
Cellular and molecular characterization of mouse lemur diseases and physiology. **a**. L2 intact lung showing metastatic endometrial tumor nodules on surface. **b-c**. Hematoxylin-and-eosin-stained (H&E) sections of L2 lung showing tumor nodules extending into lung parenchyma^23^ (b) and close-up of tumor cells (c). **d-e**. H&E section of primary endometrial tumor in the uterus of L2 (d), which metastasized to the lung, and of L3 (e), which metastasized locally. Full micrographs available at the Tabula Microcebus web portal. **f**. UMAP of lung cells (L1-L4, 10x and SS2 datasets) integrated by FIRM and colored by compartment (and by cell type designation in Fig. S10a). Note unique cluster (arrow) of epithelial cells, identified as metastatic tumor cells. **g**. UMAP of uterine cells (L3, 10x dataset) colored by compartment (and by cell type designation in Fig. S10b). Note cluster of uterine epithelial cells (arrowhead), designated as non-ciliated epithelial cell of uterus (*FXYD4+ MUC16+*), presumed to be the primary endometrial tumor cells. **h**. UMAP of all atlas cell types according to their overall transcriptional similarity (L1-L4, 10x dataset)^6^. Data points are cell types (unique combination of tissue and free annotation), colored by compartment. Inset (boxed) shows close-up of indicated UMAP region with cell types colored by compartment (circle fill) and tissue of origin (circle border), and labeled with their cell type designation. Note metastatic tumor cells (Met, arrow) from L2 lung and adjacent (molecularly similar) uterine non-ciliated epithelial cells (*FXYD4+ MUC16+*, arrowhead) from L3, suggesting uterine origin of the metastatic tumor. Cell type abbreviations are Lung: Met (uterine metastasis); Kidney: PCT (proximal convoluted tubule), PST (proximal straight tubule), LoH thin D (loop of Henle thin descending limb), LoH thin A (loop of Henle thin ascending limb), LoH thick A (loop of Henle thick ascending limb), DCT (distal convoluted tubule), CDp (collecting duct principal cell), CDi-α (alpha intercalated cell), CDi-β (beta intercalated cell), MD (macula densa). **i**. Sina plot showing Pearson’s correlation coefficients between L2 lung metastatic cells and other atlas cell types (10x and SS2 datasets, colored by compartment). Note the high correlation with uterine non-ciliated epithelial cells (*FXYD4+ MUC16+*). **j**. Dot plot showing average expression in lung and uterine epithelial cell types, separated by lemur individual, of endometrial (and ovarian) cancer (EC) serum marker genes and genes (indicated by ^*^) known to be amplified, overexpressed, or mutated in EC with their cognate ligands (red), receptors (blue), and/or modulators (green)^84,87–89,158–164^. Also plotted are representative genes with enriched expression in lung metastasis compared to other lung epithelial cell types, and genes enriched in uterine *FXYD4+ MUC16+* cells (presumed primary tumor) compared to other uterine epithelial cell types. [], description of gene identified by NCBI as a gene locus ([GZMBL], LOC105864431; [AOX2], LOC105856978; [AKR1B10L], LOC105857399 and LOC105860191). **k-l**. FIRM-integrated UMAP of adipocytes and adipo-CAR cells (10x and SS2 datasets) colored by cell type (k) or expression of the indicated genes (l). Note that adipocytes formed two main populations, distinguished by expression of classical white (e.g., *NNAT*, panel l left) and brown (e.g., *UCP1*, panel l right) adipocyte markers, and designated as *UCP1*lo and *UCP1*hi, respectively. *UCP1*lo population formed two subclusters in the UMAP that differed only in total gene and UMI counts per cell, and not in expression of any biologically significant genes (Fig. S11a-c). **m**. Distribution of *UCP1*hi (pink) vs. *UCP1*lo (blue) adipocytes in indicated fat depots and organs with ≥5 adipocytes (10x and SS2 datasets, all from individual L2, except combined fat depots from L4). N cell, number of adipocytes detected in each depot/organ. **n**. Dot plot of expression of indicated cell type markers and differentially-expressed genes in *UCP1*hi and *UCP*1lo adipocytes, adipo-CAR cells, osteoblasts, osteo-CAR cells, and chondrocytes (L1-L4, 10x dataset). Note classical brown adipocyte marker *CIDEA* and white adipokine *RBP4* (both indicated by ^*^) are surprisingly equally expressed across all adipocytes. [], description of gene identified by NCBI as a gene locus ([ATP1A2], LOC105862687; [COX7A1], LOC105876884; [Uncharacterized 1], LOC105854963).

The presumptive primary tumor cells in the uterus of L3, based on co-expression of the classical human endometrial and ovarian cancer serum markers Cancer Antigen 125 (*CA125/MUC16*) and *HE4/WFDC2*^*82–84*^, showed enriched expression of *OXTR*, as well as *MYC* and *ERBB2* (Fig. 6j), two genes commonly amplified, overexpressed, or mutated in human type 2 endometrial tumors^85,86^. In addition, these tumor cells expressed inhibin-βB (*INHBB*) and thus likely produced INHBB homodimer (activin B) (Fig. 6j), which correlates with higher grade endometrial tumors and is known to promote cancer cell migration and invasion^87,88^. The lung metastasis also expressed *ERBB2*, and gained expression of its binding partner *EGFR* and specifically the ligand *EGF*, suggesting progression to autocrine mitogenic signaling during metastasis. In addition, the metastatic cells showed altered expression in several hormone ligands and receptors^42^, including loss of estrogen receptor (*ESR1*) (Fig. 6j), which correlates with more advanced human tumors^89^. Endometrial cancer of these lemurs thus molecularly and histologically mimics the more aggressive form of the human disease, including its propensity to metastasize; however, further studies of the lemur cancer will be necessary to validate this. The mouse lemur presents a promising model to explore susceptibility factors, pathogenetic mechanisms, and therapies - particularly anti-angiogenic^90^ (VEGFR), anti-EGF/EGFR^91^, and endocrine^92,93^ (e.g., ESR1) therapies given expression of these potential targets in both the lemur and human tumors (Fig. 6j). Conversely, human therapies might help control the lemur disease^94^.

One notable aspect of mouse lemur physiology is their dramatic annual oscillations in body weight, metabolism, and body temperature as they enter a hibernation-like (torpor) state during the dry, resource-poor winter season, providing a valuable model for primate seasonal (circannual) rhythms and regulation of metabolism, adipose biology, and body weight^11,15,95–97^. We analyzed four major mouse lemur fat depots (subcutaneous adipose tissue (SCAT), mesenteric (MAT), interscapular brown (BAT), and perigonadal (GAT)), and identified hundreds of adipocytes that expressed canonical adipocyte markers including lipid biosynthetic and metabolic genes (e.g., *PNPLA2, FASN, GPD1, CIDEC*) and adipokines (*ADIPOQ, CFD*)^98,99^ (Fig. 6k-n, Fig. S11a-b, f). With these markers we also identified rare adipocytes in seven other tissues (Fig. 6m, Fig. S11a, d). In total, we captured 1,231 adipocytes and adipo-CAR cells, most of which (>95%) express mature adipocyte markers (*CEBPA, GPD1, PPARG*)^98^ (Fig. 6n).

Mouse lemur adipocytes showed two surprising features. Although they express most well-established adipocyte markers, they curiously did not robustly express the classical adipocyte hormone leptin (*LEP*), which is highly expressed by human and mouse adipocytes and regulates food intake, energy expenditure, and body weight^100^ (Fig. S11f). *LEP* expression was detected in only 0.6% of mouse lemur adipocytes (7 of 1,231 cells) and at a low level (average of 3.2 transcripts per 1e4 reads), and also sparsely and at even lower levels in several unrelated cell types. However, its receptor *LEPR* is selectively and highly expressed in a similar cellular pattern as in humans and rodents^42,101^ (Fig. S11f). Perhaps *LEP* expression is inducible in lemur adipocytes depending on season, age, diet, or body weight^102^, or some occult cellular source (or another gene) has usurped its function.

The second surprising aspect of mouse lemur adipocytes is the blurring of the distinction between white and brown adipocytes. Aside from the bone adipo-CAR cells that may be adipogenic progenitors (Fig. 6n), the bulk of the adipocytes formed two continuous populations (Fig. 6k-l, Fig. S11a-c). These populations were distinguished by expression of Uncoupling Protein 1 (*UCP1*), the canonical thermogenic brown adipocyte marker^103^. We designate the *UCP1*-high population in mouse lemur as “brown-like” because they also express higher levels of known thermogenesis regulators including genes involved in fatty acid metabolism (e.g., *CPT1B, SLC27A2, FABP3*), mitochondrial respiration (*LOC105876884* (*COX7A1*)), transport ATPase (*ATP1A2*/*LOC105862687*), and adrenergic signaling mediated lipolysis (*KCNK3*)^104^ (Fig. 6n). We designate the *UCP1*-low population as “white-like” because they show enriched expression of many white adipocyte genes including *APOE, NNAT*, and *DPT*, despite their expression (albeit low) of the brown-defining gene *UCP1*^*105–107*^. Further blurring the white-brown distinction, both the classical brown marker *CIDEA* and the classical white adipokine *RBP4* were equally expressed across all adipocytes^108,109^ (Fig. 6n). These mixed molecular signatures suggest that the distinction between white and brown adipocytes is less strong in *M. murinus*, and the observed continuum between the populations suggests potential interconversion between white-specific lipid storage and brown-specific thermogenesis, affording functional plasticity to accommodate the dynamic energy needs of this seasonally-cycling primate.

There was no exclusively brown-like fat depot among the depots surveyed: each site contained exclusively white-like adipocytes or a mix of both (Fig. 6m). And, different depots did not cluster separately and did not differentially express any biologically significant genes, except for gonadal adipocytes, which show higher levels of *S100A8*/*9*/*12, IL1B*, and metallothionein genes *LOC105866476* (*MT2A*) and *LOC105866554* (*MT1E*) (Fig. S11a, e), known in humans and rodents to correlate with inflammatory and feeding status and insulin-resistance^110–113^. It will be critical to systematically study the seasonal changes in adipocyte gene expression in each depot to explore adipocyte plasticity, how it is controlled, and its role in seasonal physiology and behavior.

### 6. Expression patterns of primate genes missing in mouse

Mouse lemurs could be especially valuable for the study of human genes that are absent, or expressed differently^6^, in other model organisms such as mouse. By comparing lists of orthologous protein-coding genes in human, lemur, and mouse (Table 7, integrating NCBI, ENSEMBL and MGI homology assignments), we identified 539 human genes for which there are orthologs in lemur (425 orthologs annotated in NCBI) but not mouse (Fig. 7a, Table 8), what we will refer to here for simplicity as “primate-selective” (PS genes). At least twenty-four of the PS genes are known to cause human disease or phenotypes^114^ (Table 8). Others play important roles in human physiology, such as motilin (*MLN*) and its receptor (*MLNR*) in gastrointestinal motility^115^, *CD58* in antigen presentation to T cells (see below), and *FCAR* (IgA receptor), *CXCL8*, and *S100A12* in the inflammatory response. Gene set enrichment analysis showed PS genes are enriched in transcription factor activity and regulation (GO term: 0001067, 0000977) and herpes simplex virus 1 infection (KEGG: 05168), including many that encode zinc finger proteins (Table 9). Nearly all (94%) of the 425 NCBI-annotated PS genes were expressed in the lemur atlas. Their global expression patterns across 739 lemur cell types are shown in Fig. S12a-b (Table 8). Some were selectively expressed (or depleted) in specific compartments (166 genes) and/or in specific organs (99 genes) (Fig. 7a-e, Fig. S12c-h). Many were specific to male germline cells, immune cells and neurons, indicating substantial evolutionary gene plasticity in these compartments. Many PS genes exhibited similar expression patterns in human and lemur, including chemokine *CCL16*, neutrophil granule proteins *AZU1* and *TCN1*, as well as many other immune-related genes, such as *CD1B, CD1E, IL32*, and *KLRF1* (*NKp80*) (Fig. 7f, Fig. S12i, Table 8). Notably, some of these PS genes have functions that remain to this day poorly characterized, such as endothelium-specific transmembrane protein *TM4SF18*, myeloid-specific *SIGLEC14*, broadly-expressed *TCEAL4*, and sperm and progenitor-specific *SMKR1, CT83, DDX53, C5orf58/C23H5orf58* (human/lemur symbols), *C12orf54/C7H12orf54, C16orf82/C17H16orf82* (Fig. 7f, Fig. S12i). Mouse lemur provides a system to dissect the functions of these genes in organismal physiology, development, and disease.

**Figure 7.**
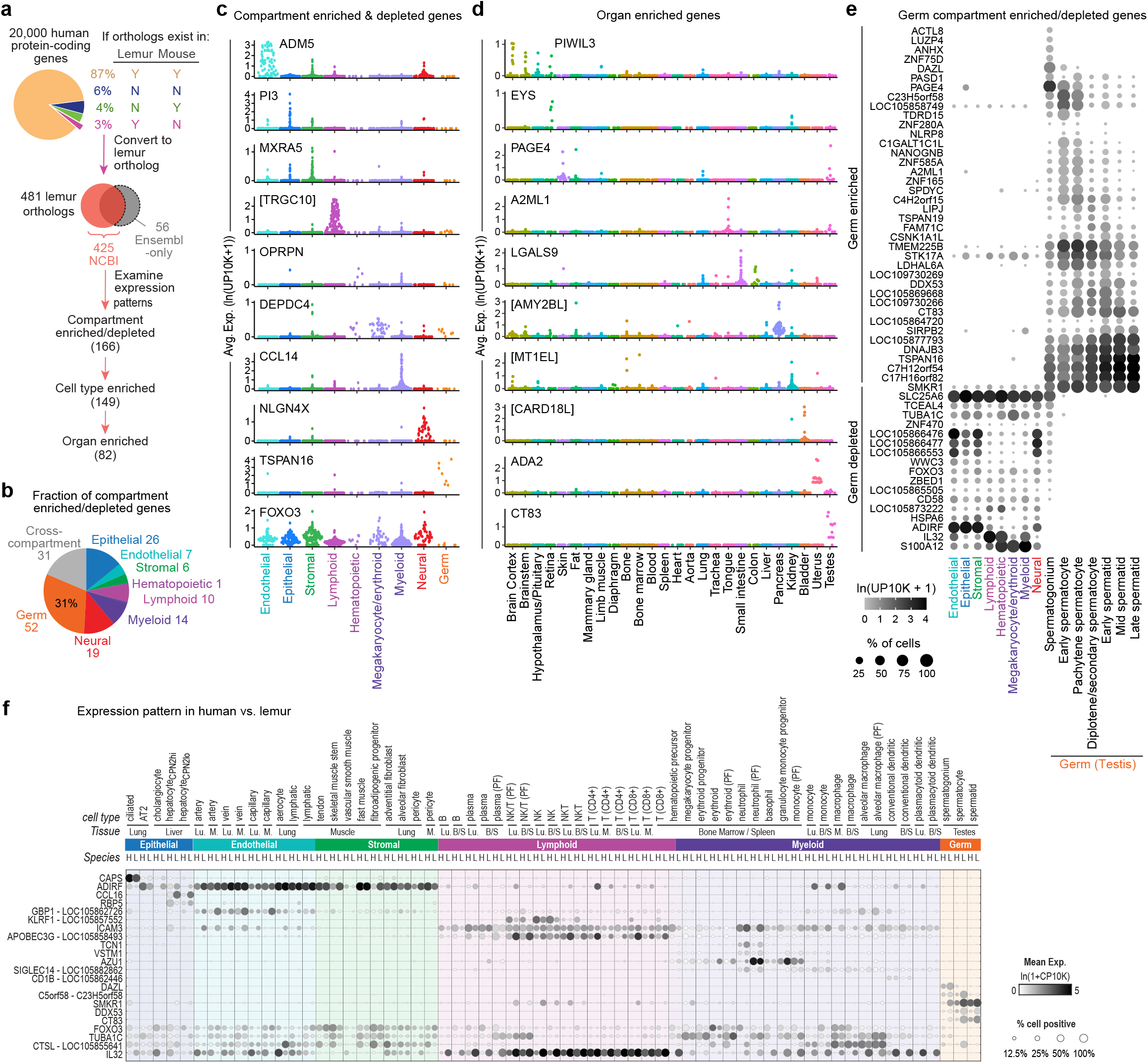
Expression patterns of primate selective genes in mouse lemur. **a**. Percent of the ∼20,000 human protein-coding genes with identified ortholog(s) (by NCBI, Ensembl, and/or MGI) in lemur and mouse. The 539 (3%) human genes that share an ortholog only with lemur (referred to as primate selective (PS) genes) correspond to 481 lemur genes (329 annotated by both NCBI and Ensembl, 96 by NCBI only, and 56 by Ensembl only). Expression patterns of the 425 NCBI-annotated lemur orthologs were examined and categorized. **b**. Pie chart showing fraction of PS genes (166) enriched (or depleted) in a specific tissue compartment. Genes enriched (or depleted) in more than one compartment are indicated as ‘cross-compartment’. Note about one-third (31%) of the genes are enriched (or specifically depleted) in the germline compartment. **c-d**. Sina plots showing expression of representative PS genes that are compartment-enriched (or specifically depleted for *FOXO3*) (c) or organ-enriched (d) across atlas cell types (10x dataset), with cell types (data points in figure) grouped by compartment (c) or by organ of origin (d). Note that organ-enriched genes are almost always enriched in specific cell types of the tissue, for example *LGALS9* in small intestine and colon enterocytes, *EYS* in retinal cones, and *ADA2* in uterine epithelial cells. [], description of gene identified by NCBI as a gene locus ([TRGC10], LOC105878255; [AMY2BL], LOC105863954; [MT1EL], LOC105866478; [CARD18L], LOC105862464). **e**. Dot plot showing average expression of PS genes that are enriched or depleted in germ compartment across non-germ compartments (averaged across all cells in the compartment) and different germ cell types (10x dataset). [], description of gene identified by NCBI as a gene locus ([H2BC12], LOC105858749; [Uncharacterized 1], LOC109730269; [AK1], LOC105869668; [HSFX4], LOC109730266; [SPANXN4], LOC105864720; [MT2A], LOC105866476, [MT2AL], LOC105866477, [MT1XL], LOC105866553, [RPL36AL], LOC105873222). **f**. Dot plot showing cross-species expression of selected PS genes in 63 orthologous cell types in human and lemur, as analyzed in the evolutionary comparison analysis of the accompanying Tabula Microcebus manuscript^6^. Rows are orthologous genes, indicated with the respective human gene symbols and followed by the respective lemur gene symbol if different. Columns are cell types, displayed as doublets of the respective expression in human (H) and lemur (L). Cell types are ordered first by compartments, then by tissue, and finally by species. Note the conserved expression patterns of these genes in both species, with many genes known to regulate important functions, particularly immune functions, such as chemokine *CCL16* in hepatocytes, granule proteinsazurocidin (*AZU1*) and transcobalamin 1 (*TCN1*) in neutrophils, cytokine-stimulating *KLRF1* (*NKp80*) in NK/NKT cells, antigen-presenting *CD1B* and *CD1E* in conventional (and not plasmacytoid) dendritic cells, and interleukin 32 (*IL32*) in T, NK, and NKT lymphocytes.

### 7. Identification and characterization of natural mutations using the cell atlas

A critical step in establishing an emerging model organism is development of methods for analyzing the phenotypes and function in vivo of individual genes, especially those of great interest like the PS genes described above. We used the transcriptomic atlas to identify and analyze the first nonsense mutations and their molecular phenotypes in mouse lemur (Fig. 8a, Fig. S13a-b). Short-read, whole genome sequencing was performed to uncover natural mutations in the genomes of the lemurs that were transcriptomically profiled for the atlas. Single-nucleotide polymorphisms (SNPs) and short insertion-deletion mutations (indels) were then systematically mapped by comparing the obtained sequences to the reference genome Mmur 3.0. We focused on genes carrying putative null nonsense alleles that were segregating in the laboratory colony and present in only one or two of the profiled individuals, then examined them for transcriptional phenotypes in our cell atlas data.

**Figure 8.**
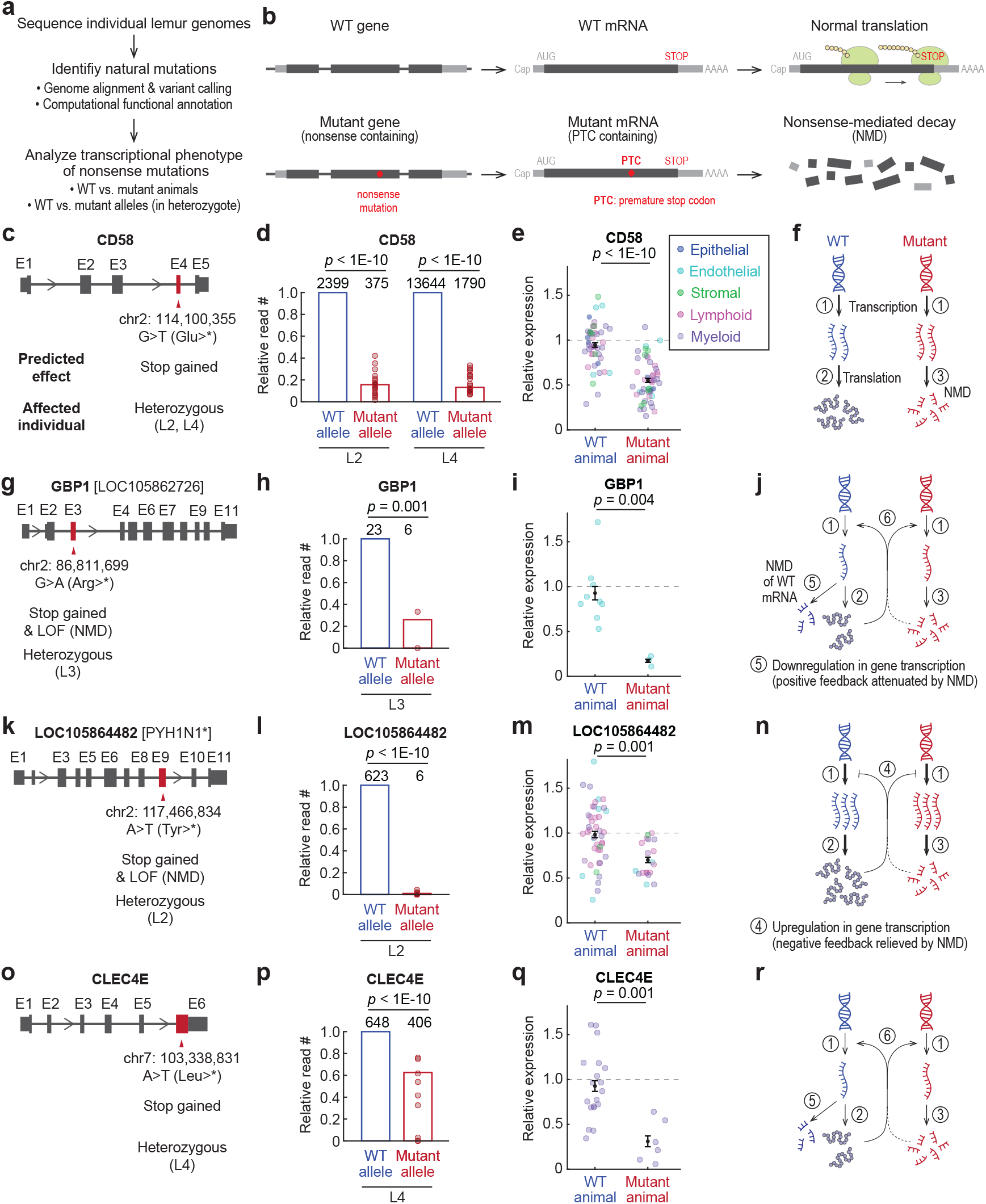
Identification and analysis of nonsense mutations and their transcription phenotypes. **a, b**. Overview of the analysis (a) and schematic of the nonsense-mediated RNA decay (NMD) pathway (b). **c-r**. Nonsense mutations and their transcriptional consequences for four lemur immune genes: CD58, a ubiquitously expressed CD2-binding T cell activator (c-f); GBP1, an interferon-inducible GTPase expressed in endothelial cells; LOC105864482 (PYH1N1 homolog), an interferon-inducible protein abundant in T and NK cells (k-n); and CLEC4E, an innate immune regulator expressed in neutrophils and monocytes (o-r). Three of these genes (CD58, GBP1, PYH1N1) are present in lemur and human but missing in mouse (see Fig. S13c for the cross-species expression patterns). (**c, g, k, o)** The identified nonsense mutation and its predicted effect (see also Fig. S13a), and the affected individual(s). Exons (E, affected exons in red). Note only two of the mutations (g, k) were computationally predicted from the sequence to cause NMD (see Methods), but varying degrees of NMD were observed for all four mutations. (**d, h, l, p)** Relative transcript read counts (10x datasets, values normalized to the WT value in the same individual) in affected individuals for reads aligned to WT (blue bars) vs. the mutant (red bars) alleles. Data points show values for all tissues (10x batches) with detected reads that aligned to the mutation site. Numbers above bars, total read counts for the indicated allele and individual. Note that transcript reads analyzed are only those that aligned to the mutation site, a small fraction of all sequencing reads for the gene (see Fig. S13b). *p*-values, one-tailed binomial tests (combining reads from all tissues). (**e, i, m, q)** Relative expression levels (normalized to the mean expression level of the gene in WT animals, averaged across all cells) of the affected genes in WT vs. mutant animals. Data points show different cell types (10x datasets), colored by compartment as indicated. Cell types with fewer than 35 cells were excluded. Error bars, mean and 95% confidence intervals, estimated across cell types. *p*-values, two-tailed student *t*-tests. (**f, j, n, r)** Schematic models of how the nonsense mutation affects expression of the corresponding gene. **f**. Simple model in which NMD affects only the mutant and not the WT *CD58* transcript; thus ∼90% depletion of the mutant transcript (d) results in ∼45% reduced transcripts in heterozygous mutants (e). **n**. NMD pathway reduces mutant *LOC105864482* transcripts, and the gene exhibits compensatory upregulation in transcription; thus, despite almost complete (99%) elimination of the mutant transcripts (l), the heterozygotes showed only a ∼30% decrease in total gene transcripts (m). **j, r**. NMD pathway partially destroys both mutant and WT transcripts of *GBP1* (j) and *CLEC4E* (r), or there is attenuation of a positive-feedback loop; thus transcriptional reduction in heterozygotes vs. WT animals is greater than expected from the simple model (n).

Most of the identified nonsense mutations were heterozygous, so we leveraged our scRNA-seq data to discriminate transcripts from each allele to quantify the effect of nonsense-mediated mRNA decay (NMD) (Fig. 8b). For nearly all autosomal genes, both alleles are transcribed at similar levels^116^. But in an individual with a heterozygous nonsense mutation, the mutant (nonsense mutation-containing) transcript should be selectively degraded by the NMD pathway and hence underrepresented in the transcriptomic profiles relative to the corresponding wild type (WT) mRNA, with the magnitude of difference reflecting the efficiency of NMD degradation of the mutant RNA. In this way, we compared the number of scRNA-seq reads containing the nonsense mutation to that of the wild type sequence in the profiled heterozygous individuals.

Fig. 8c-r shows the transcriptional phenotypes of heterozygous nonsense mutations we identified in four genes of interest whose human orthologs function as immune regulators. Two of the genes (*CD58*, T cell CD2 ligand; *GBP1*, interferon-inducible GTPase in innate immunity) are among the PS genes described above that are present and expressed similarly in lemur and human but absent in mouse (Fig. S13c). The third, *LOC105864482*, is a homolog (46% protein sequence identity) of human interferon-inducible protein *PYH1N1*, with orthologs restricted to primates (and flying lemurs, a close primate relative)^117^. For all three genes, we found that the nonsense transcript reads were substantially depleted compared to the WT transcripts in the same individual (74-99%, Fig. 8d, h, l), suggesting efficient recognition and destruction of the mutant RNA by the NMD pathway. For the fourth gene, *CLEC4E*, an immune regulator conserved across all three species, nonsense mutant transcript reads were only 37% depleted (Fig. 8p). This indicates less efficient degradation of the mutant RNA by the NMD pathway, consistent with the location of the nonsense mutation in the last exon of the gene (Fig. 8o), which is known to prevent or reduce NMD activity^118^. In this manner, the transcriptomic atlas facilitated both identification and quantification of the most direct molecular consequences (NMD-mediated RNA destruction) of these nonsense mutations.

The atlas also allowed determination of the indirect consequences of the nonsense mutation on expression of the WT allele and overall expression of the gene, by comparing total transcript levels of the gene in heterozygous versus WT individuals. For *CD58*, heterozygous mutants exhibited ∼45% less *CD58* expression than WT individuals (Fig. 8e), the level expected based on the observed ∼90% depletion of the mutant transcript (Fig. 8d). This suggests that transcription of the WT allele was unaffected by the nonsense-containing transcripts (Fig. 8f).

However, for *LOC105864482*, the heterozygous lemur showed only a ∼30% overall reduction in *LOC105864482* expression relative to WT individuals (Fig. 8m) despite almost complete (99%) elimination of the mutant transcript (Fig. 8l). This suggests compensatory upregulation of the gene (Fig. 8n). In contrast, the *GBP1* and *CLEC4E* heterozygous mutants showed more than the expected reduction in their overall expression (Fig. 8h-i, p-q), suggesting that NMD somehow also reduces (in trans) transcripts of the respective WT allele^119,120^, or there is attenuation of a positive feedback loop (Fig. 8j, r). Thus, the transcriptomic atlas enabled identification and transcriptomic characterization of nonsense mutations in mouse lemur genes, demonstrating the direct and secondary consequences of the mutations at the transcript level that highlight gene-specific differences in regulation of the NMD pathway.

## DISCUSSION

Mouse lemurs have many desirable characteristics for a primate model organism, including their easy lab husbandry, short generation time, and genetic proximity to human. Here we exploited an extensive transcriptomic atlas of mouse lemur^6^ to establish a foundation for genetic and molecular studies of mouse lemur, and to prioritize genes, splice junctions, physiology, and disease for future study, which have historically taken decades to achieve for model organisms.

First, we used the massive transcriptomic dataset to uncover mouse lemur genes, define gene structures, and aid in gene naming. This identified and named/orthologized thousands of mouse lemur genes (Section 1, 3) and hundreds of thousands of splice forms (Section 2) missed by the conventional genome annotation pipelines, including genes in the most variable, repetitive, and difficult to annotate loci (e.g., Ig and MHC genes). This provides a comprehensive genomic foundation for future mouse lemur studies.

Second, we used the full-body transcriptomic atlas to analyze mouse lemur physiology and disease with cellular and molecular precision. Our systematic analysis of immune cells and genes provides a deep understanding of the lemur immune system, including molecular mapping of immune cell development, trafficking, tissue-specialization, and local and global inflammatory responses to infection (Section 4). Likewise, systematic analysis of hormone and receptor gene expression^42^ provides a comprehensive view of the lemur endocrine system and unprecedented insight into general properties of endocrine networks. And, by analyzing the transcriptomic data in conjunction with the extensive clinical metadata and histopathology collected for each individual^23^, we ascertained rich molecular portraits of lemur disease such as the full pathogenic sequence of uterine adenocarcinoma, from primary tumor to metastasis, secondary infection, and the local and global inflammatory responses (Section 5). “Molecular cell autopsies” like this herald a new era of pathology, providing both local and systems-level understanding of the pathogenic mechanisms of disease. This approach is extendable to other model organisms and disease processes, including human pathologies.

Third, we identified high priority areas for study in mouse lemur, most particularly its genes, physiology, and diseases that are conserved in human, or unique in lemur, but absent or substantially divergent in mice. These include molecular features of the immune program (Section 4), of lemur adipocytes that exhibit dramatic seasonal rhythms, and of metastatic endometrial cancer that resembles the human malignancy (Section 5). Our classical autopsies uncovered a variety of other human-like pathologies including cataracts, osteoarthritis, chronic kidney disease, and amyloidosis^23^, and prior studies identified an Alzheimer’s-like neurodegenerative disease common in aged mouse lemurs^121^. A top priority for future studies is the more than 400 primate genes that are missing in mice, especially those that are already implicated in human disease (Section 6). And beyond the missing genes are the many genes present in mouse but whose splice isoforms (Section 2) or cellular expression patterns^6^ we found differ substantially from those of lemur and human.

Finally, we describe an experimental pipeline for reverse genetic analysis of mouse lemur genes and biology. We identify the first null alleles in mouse lemur, including naturally-occurring nonsense mutations in three of the top priority genes — primate immune genes missing in mice — and use the cell atlas to characterize the transcriptional effects of the mutations (Section 7). In parallel, systematic forward screens for mouse lemur morphological, physiological, and disease phenotypes have identified eight human-like cardiac arrhythmias and recently succeeded in mapping the disease gene for one of them (sick sinus syndrome), a transporter with cell autonomous, primate-specific function in regulation of the sinus node pacemaker (Chang et al., accompanying manuscript). Both forward and reverse genetic approaches are thus now possible for mouse lemur.

This work establishes mouse lemur as a tractable model organism for genetic and molecular dissection of primate genes, splice forms, physiology and disease. The experimental framework developed here can be readily applied to other emerging model organisms to establish molecular and cellular foundations for exploring their unique biology. However, unlike the original mouse cell atlas obtained from cell profiles of many genetically and environmentally homogenous animals^122,123^, atlases for emerging model organisms will generally be created from fewer individuals and ones of different ages and from different genetic and environmental backgrounds, like this atlas and our recent human cell atlas^124^. Under these conditions, as shown here, it is especially valuable to profile most or all tissues from each individual, and to provide extensive metadata for each individual, such as its distinguishing features, life history, and clinical data including full autopsies and histopathology. This creates both a working reference cell atlas — crucial for understanding normal physiology and for evolutionary comparisons — but also uncovers and provides insight into valuable pathological variants, which must then be validated and explored more deeply in follow up studies. There is also great value in using the same profiling strategies and analytical approaches for each organ and organism to ensure fidelity of evolutionary comparisons. This becomes increasingly important as organism-wide evolutionary analyses expand beyond the three species compared here (lemur, human, mouse) to other emerging model organisms and ultimately the full tree of life.

## Supporting information

Supplementary results

## ACKNOWLEDGMENTS

This work was supported by funding from The Chan-Zuckerberg Biohub, Howard Hughes Medical Institute and Vera Moulton Wall Center for Pulmonary Vascular Disease; unrestricted grant from Research to Prevent Blindness and NEI P30-EY026877 to the Stanford Department of Ophthalmology to Albert Y. Wu; The Hong Kong University of Science and Technology (start-up grant R9364), The Hong Kong University of Science and Technology Big Data for Bio Intelligence Laboratory (BDBI), and The Chau Hoi Shuen Foundation (R9056) to Angela Ruohao Wu; NSF-DBI-1701984 to Anne D. Yoder; funding from Developmental and Stem Cell Biology Graduate Program to Aris Taychameekiatchai; funding from Hong Kong Research Grant Council (16307818, 16301419, 16308120, 16307221, C6021-19E), the Hong Kong University of Science and Technology (startup grant R9405), and The Hong Kong University of Science and Technology Big Data for Bio Intelligence Laboratory (BDBI) to Can Yang; NIH grant R01 GM122951 and R35 GM136433 to Margaret Fuller; funding from Shanghai Sailing Program (21YF1410600) to Jingsi Ming; NIA 1K99AG066963 to Thomas Ambrosi; NIH DP2AI138242 to Iwijn De Vlaminck; National Sciences and Engineering Research Council of Canada fellowship PGS-D2 to Michael F.Z. Wang; NIH R35 GM139517, R01 GM116847, R35 GM139517, NSF MCB1552196 to Julia Salzman; NSF Graduate Research Fellowship DGE-1656518 and a Stanford Graduate Fellowship to Julia Olivieri; Cancer Systems Biology Scholars Fellowship (Grant R25 CA180993) and Clinical Data Science Fellowship (Grant T15 LM7033-36) to Roozbeh Dehghannasiri; Urology Care Foundation Research Scholar Award Program and AUA Western Section Research Scholar Fund II to Hosu Sin; Stanford Graduate Fellowship/HHMI/NIH CMB Training Grant to Yue Zhang; American Cancer Society Postdoctoral Fellowship to SoRi Jang; Walter V. and Idun Berry Postdoctoral Fellowship to Andrea R. Yung; NSF Graduate Research Fellowship and Stanford Graduate Fellowship to Youcef Ouadah; the Wu Tsai Neurosciences Institute Interdisciplinary Scholar Award to Shixuan Liu; European Community’s 7th Framework Programme (FP7/2007-2013) under grant agreement Nbr 278486 (DEVELAGE), from the Fonds Unique Interministériel and Région Languedoc-Roussillon under grant agreement Nbr 110284 (DiaTrAl), and from the Fondation Plan Alzheimer (PRADNET) to Jean-Michel Verdier and Corinne Lautier; NIH R01 AI024258 to Peter Parham and Lisbeth A. Guethlein; NIH R01DC016892 to Wan-Jin Lu; NIH P30DK116074 to Yan Hang; NSF Graduate Research Fellowship to Connor V. Duffy; Postdoctoral Fellowships from the DFG (NE 2006/1-1) and California TRDRP (25FT-0011) to Patrick Neuhöfer; a NovoNordiskFonden Start Package grant (0071116) to Antoine de Morree; funding from Independent Research Fund Denmark (DFF–5053-00195) and Lundbeck Foundation (R232-2016-2459) to Jean Farup; NIH AG068667, AR073248 and AG036695 to Thomas A. Rando; funding from Wu Tsai Neurosciences Institute to Tony Wyss-Coray; NSF BCS 0647402 to Liza Shapiro and E. Christopher Kirk.

## METHODS

### Data availability

Tabula Microcebus data (including cell coordinates of FIRM-integrated UMAPs and MHC gene expression data) and cross-species datasets are in Figshare (https://figshare.com/projects/Tabula_Microcebus/112227). Raw sequencing data (fastq files) are available at https://app.globus.org/file-manager?origin_id=c9fc0a15-54a0-4182-8d64-fd8afc12f1fc&origin_path=%2F. Tabula Microcebus web portal (https://tabula-microcebus.ds.czbiohub.org/) also allows for interactive visualization of the scRNA-seq data with cellxgene and the histological images of the mouse lemur tissues analyzed.

### uTAR analysis to identify unannotated genes

To uncover unannotated transcriptionally active regions (uTARs), we employed the workflow developed by Wang et al.^25^ for scRNA-seq data that identifies transcriptionally active regions (TARs), genome regions with abundant transcript alignments, using a previously published tool groHMM^125^. Briefly, all mouse lemur 10x datasets were aligned to genome assembly Mmur 3.0 using STAR with default parameters, without gene annotation indexing, and transcribed regions were predicted using the groHMM tool. TARs within 500 bps of another were combined into a single TAR and kept if it was expressed in at least 2 cells of the droplet-based atlas. The detected TARs were then separated into annotated TARs (aTARs) and unannotated TARs (uTARs), based on whether the region is currently annotated as a gene in NCBI Annotation Release 101 of Mmur 3.0. This identified that in the mouse lemur genome (2487 Mbps), aTARs and uTARs cover 284 and 42 Mbps, respectively.

To filter out transcriptional and sequencing noise from biologically significant uTARs, we then examined if a uTAR is differentially-expressed across cell types by Wilcoxon rank-sum test. This analysis was performed separately for each tissue and individual lemur. A differentially-expressed uTAR (DE-uTAR) is defined as a uTAR with a significant Bonferroni-corrected *p*-value <0.05 from the Wilcoxon rank-sum test, expressed in ≥25% cells of a cell type (cells of the same free annotation), and cell type average expression level is ≥ 1.65 (e^0.5^) times the average of other free annotation cell types in the same tissue. Some of the uTARs passed the differential expression test in multiple cell types and/or tissues. Together, a total of 4003 unique DE-uTAR were identified. To infer their identity, we applied BLASTn on each of the DE-uTAR against the nucleotide collection (nt) database (with a threshold of maximum e-value of 0.01 and a minimum bit score of 50), using either the entire length of the uTAR or the peak coverage region (full width at half maximum region around the absolute peak in coverage after Gaussian smoothing within the uTAR location). Occasionally, multiple uTARs aligned to the same gene in another species. The genome location and inferred homology of all DE-uTARs as well as their expression levels across the atlas 10x cells are provided in Table 1. There were 30 DE-uTARs with a BLASTn result that corresponded to one of the 2055 human genes without a mouse lemur ortholog, which are likely genes missed from NCBI’s and Ensembl’s annotations of Mmur 3.0 (Table 1). Note, 14 out of these 30 genes have no mouse ortholog, suggesting these are PS genes.

DE-uTARs were also classified as protein-coding or noncoding using Orpheum (Botvinnik et al., 2021), with sequences containing 95% or more k-mers (k=9) matching a reference database of mammalian proteins from UniProt assigned as putatively protein-coding, and otherwise, assigned as noncoding. Coding sequences were then annotated using DIAMOND blastp^126^ and noncoding sequences were annotated using Infernal cmscan (https://www.ebi.ac.uk/Tools/rna/infernal_cmscan/). Some uTARs had both coding reads and non-coding reads, potentially representing incompletely spliced transcripts or untranslated regions.

To detect the developmental trajectory of the sperm lineage cells in the testes 10x dataset by the uTAR expression data, we followed the same procedure as detecting the trajectory by gene expression data, which is described in the accompanying manuscript^6^. The analysis included a total of ∼50,000 uTARs that have transcript reads in the testis dataset. Each uTAR was treated as a gene and data were first normalized for scRNA-seq library size (to 10,000 total uTAR transcripts per cell) and natural log-transformed. The top highly-variable uTARs (∼1500) were then used for principal component analysis. The top 20 principal components that were not driven by extreme outlier data or immediate early genes were used to construct a 2D UMAP using cell-cell Euclidean distances as input. The pseudotime developmental trajectory was then identified as the density ridge of the data in this 2D UMAP by automated image processing.

Cells were assigned to the trajectory by the shortest connecting distance. The cells’ pseudotime trajectory coordinates were linearly normalized such that the trajectory started at 0 and ended at 1, and then were compared to the pseudotime coordinates derived from the gene-based trajectory using Pearson’s correlation. The uTAR expression data were similarly pre-processed (normalized, scaled, and UMAP embedded) in other analyses, including when comparing the UMAP cell distribution patterns of the 10x colon dataset and in the silhouette coefficient analysis (Fig. S2a-b).

To measure the consistency of cell distribution patterns for 10x datasets embedded in a UMAP using the gene, aTAR, and uTAR-expression space, silhouette coefficient values were calculated for each dataset (separated by tissue and lemur individual, and sequencing channel). Cells were grouped according to the expert cell type designation (free annotation). The silhouette coefficient value for each cell *i* in a dataset was calculated as *s*(*i*) = (*b*(*i*) − *a*(*i*))/ *max*{*b*(*i*),*a*(*i*)}, where *a*(*i*) is the mean in-group distance (mean distance of cell *i* to the other cells in the same cell type) and *b*(*i*) is the minimal out-group distance (minimal distance of cell *i* to any cell in the tissue of a different cell type). The cell silhouette coefficient values were then averaged across each dataset to derive the dataset-averaged silhouette value - an overall score representing how well each cell type co-clusters and separates from other cell types in the UMAP embedded space, with higher positive values representing better separation. The silhouette coefficient values were calculated separately using the cell-to-cell distances in UMAPs based on the gene expression, aTAR expression, and uTAR expression spaces, and then compared with a box plot (Fig. S2a).

The lists of genes for each category in Fig. S2e were obtained as follows. Lists of the top n variable NCBI-annotated genes were derived by applying variance-stabilizing transformation to the entire 10x dataset and selecting the genes with top n transformed variance. The list of the genes annotated by Ensembl only (not by NCBI) were detected by comparing the genomic positions of mouse lemur genes annotated in the two databases, searching for the Ensembl-annotated gene with no overlap in NCBI. The PS genes are derived as in methods, and listed in Table 8.

To determine the amount of transcriptomic sequence information provided by the mouse lemur scRNA-seq atlas datasets, we estimated the total number of sequenced base pairs (bp) that mapped to the mouse lemur genome. For the 10x datasets, the total number of aligned paired-end reads (∼1.63×10^10^) was multiplied by the number of bp per read (90), equating to 1.46×10^12^ bp, though this number is inflated by PCR duplicates. However, summing the number of unique molecular identifiers (UMIs) across all cells in the atlas (∼1.22×10^9^) and multiplying it by the number of bp per read (90), equated to 1.10×10^11^ bp – 10-fold less than the estimate with total reads. For the SS2 datasets, the total number of aligned paired-end reads (1.11×10^10^) was multiplied by the number of bp per read (100), equating to 1.11×10^12^ bp (unique reads are not possible to identify in SS2 datasets). In comparison, the total number of bulk RNAseq bp used by NCBI for gene prediction/annotation of Mmur 3.0 equates to ∼3.4×10^11^ bp (across ∼3.4×10^9^ total reads), which was calculated by summing the number of sequenced bp for each RNAseq run (all tissues and generic samples included) – information obtained from the Sequence Read Archive (Biosample ID numbers and corresponding SRA link available at https://www.ncbi.nlm.nih.gov/genome/annotation_euk/Microcebus_murinus/101/). However, this estimate is inflated because it includes bp from unaligned reads and does not correct for PCR duplicates.

### SICILIAN splicing analysis

To identify splice junctions of mouse lemur transcripts, we employed SICILIAN, a statistical method for unbiased and annotation-free detection of splice junctions^33,127^. SICILIAN computes for each discontinuous sequence read, a statistical confidence value based on features that influence sequence alignment (e.g., sequence entropy of the read, sequence mismatches, number of mapping locations for the read in the genome), and then these read-level statistical values are aggregated to compute a final confidence score (empirical *p*-value) for each potential splice junction.

Raw sequencing reads from the atlas (both 10x and SS2 datasets) were first aligned to the mouse lemur genome assembly Mmur 3.0 by the STAR algorithm with parameters chimSegmentMin = 12, chimJunctionOverhangMin = 10, chimOutType = “WithinBAM SoftClip Junctions”, and default values for the rest of the parameters. SICILIAN first extracted spliced reads that mapped to discontinuous regions of the genome. These reads could provide evidence for candidate splice junction sites but may also reflect sequencing noise or alignment error. For each of these potentially-spliced scRNA-seq reads, SICILIAN estimated a read-level confidence score, quantifying the probability that the alignment of the read is true. It then incorporated all the read-level scores for the reads aligned to each potential splice junction and computed an empirical *p*-value for the junction. The empirical *p*-value was computed for each SS2 cell or 10x channel within the dataset, then the median was calculated for these empirical *p*-values across the dataset. Junctions with median empirical *p*-value <0.1 were selected for follow-up analysis. The threshold 0.1 was determined as the optimal point in the receiver operating characteristic (ROC) curve based on simulated data. The *p*-value threshold was based on a benchmarking analysis in the SICILIAN method paper (see ROC in Fig. 1E)^33^ which identified that a threshold of 0.15 maximizes discovery sensitivity and specificity (Youden’s index) using bulk simulated RNA-seq data. Here, we prioritized specificity to identify high-confidence novel junctions and therefore used a more stringent threshold (0.1). We performed SICILIAN analysis separately on the reads from each lemur individual and scRNA-seq method (10x and SS2).

The detected splice junctions were compared to the junctions in transcripts annotated in the NCBI’s Annotation Release 101 of mouse lemur genome assembly Mmur 3.0. The detected splice junctions were categorized into five types (see Fig. 2b). Type A refers to junctions that matched an annotated splicing pattern. Type B-D refers to junctions that aligned to an annotated gene but the specific splicing pattern is unannotated; type B contains junctions in which both the donor (5’) and acceptor (3’) splice sites are annotated but not previously paired (e.g., unannotated exon skipping); type C contains junctions in which one site is annotated and the other is not (e.g., annotated donor site but unannotated acceptor site); type D contains junctions in which both splices sites are unannotated. Type E refers to detected junctions that do not align to any annotated gene.

To examine if the detected splice junctions are conserved in human or mouse genomes, we used the UCSC LiftOver tool from the UCSC genome browser^128^ and computed the fraction of junctions annotated in mouse and/or human by considering a junction as conserved only if it had successful LiftOver conversion to the other genome (i.e., both 5’ and 3’ splice sites in the lemur genome mapped successfully to unique coordinates in the other genome).

To identify cell-type specific splicing events, we performed MANOVA analysis separately on the 10x data from each tissue. Cell types with fewer than 10 cells or junctions present in fewer than two cells were removed from the analysis. To highlight splicing events with global effects, we analyzed the genes with at least two spliced reads mapping to the gene in each cell type in the tissue. Note, however, our approach can easily be extended to include more genes expressed in only a subset of cell types. Let *C* be the set of cells, and *J* be the set of junctions for this gene. Consider cell *m* ∈ *C* and junction *i* ∈ *J* for a particular gene. Let 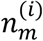 be the number of reads mapping to junction *i* in cell *m*. Then define the fraction of junctional reads mapping to junction *i* in cell *m* as: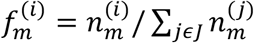. Then calculate the dataset average fraction of junction *i* as: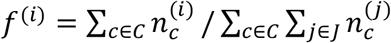. Then let the scaled z score for junction *i* in cell *m* be defined as: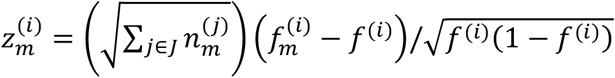. MANOVA analysis was then performed using the cellular z scores of each junction for the gene as input and the cell type (free annotation) as output. This analysis generates, for each gene, a *p*-value of all cell types in the tissue having the same multivariate mean junctional expression. Benjamini-Hochberg correction was then applied to all *p*-values. To identify a list of candidate junctions with the most significant cell type differential splicing, a stringent threshold (corrected *p*-values less than 10e-16) was used, which resulted in 545 junctions.

### SAMap analysis to study conservation of gene expression patterns across species

To compare expression patterns of homologous genes across human, lemur, and mouse, we used the SAMap method developed by Tarashansky et al.^38,39^. In addition to the lemur 10x cell atlas datasets, mouse and human 10x scRNA-seq datasets were retrieved from Tabula Muris Senis^122^ and Tabula Sapiens^124^, respectively. We applied SAMap to these datasets to compare the cell type expression patterns of homologous genes across the three species. While the SAMap algorithm does not require cell type labels, having comparable annotations simplifies the interpretation of the mapping results. Therefore, we limited the analysis to the tissues (lung and skeletal muscle) that were reannotated using the same standards as described in the accompanying Tabula Microcebus manuscript^6^. This cross-species data with new unified cell type designation can be found on Figshare https://figshare.com/projects/Tabula_Microcebus/112227.

SAMap was used to simultaneously map genes and cells across the three species (human, lemur, and mouse). For each pairwise combination of species, SAMap first detects homologous genes by bi-directional BLAST of the transcriptomes of the two species, as annotated by Ensembl and NCBI. A cross-species gene-to-gene graph is then generated with edges connecting a gene in one species and a homologous gene in the other species and edge weights assigned as sequence similarity of the gene pairs. The homology graphs from all pairwise comparisons of species are combined into one, tripartite adjacency matrix. Using this initial gene graph, SAMap projects the three single-cell RNAseq datasets into a joint, lower-dimensional manifold representation. This joint manifold allows estimation of similarity between cells and genes across species. Note that a simplified version of such cell or gene similarity quantification is to only consider the one-to-one orthologs while ignoring the many-to-one, one-to-many, and many-to-many orthologous genes as well as the non-orthogonal relationship between non-orthologous but homologous genes, both of which are incorporated by the SAMap method. Next, the expression correlations between homologous genes are calculated in the initial joint manifold to reweight the edges of the gene-gene homology graph. Using the reweighted homology graph as the new input, SAMap then iterates until convergence to generate a final joint manifold. The expression correlation between homologous genes of the two species calculated across the joint manifold quantifies the similarity of the two genes’ expression patterns. Homologous gene pairs with an expression correlation higher than 0.3 are deemed expression homologs - homologs that share similar expression patterns across mapped cells. Triads of mapped expression homologs from human, lemur, and mouse datasets were identified.

We then examined, for each expression homolog triads, whether the three gene pairs were assigned as orthologous genes by NCBI and/or Ensembl (see Table 7). We further examined for each expression homolog triad whether the lemur gene is named or unnamed in NCBI Annotation Release 101 of the mouse lemur genome assembly Mmur 3.0 (with only a locus identifier, e.g., “Loc “ or “orf”). Expression homolog triads were then categorized into three types (Fig. 3b). Type “named ortholog” refers to triads that consist of 3 orthologous gene pairs and the lemur gene is named accordingly. Type “unnamed ortholog” refers to triads of 3 orthologous gene pairs but the lemur gene is unnamed. Type “non-ortholog” refers to triads of less than 3 orthologous gene pairs, regardless of the naming status of the lemur gene. Table 4 lists the expression homologs detected in this study.

### Ig/B cell receptor (BCR) analysis

To improve the annotation of the mouse lemur Ig/BCR loci, we first used BLAST^129^ with human BCR genes (retrieved from ImMunoGeneTics (IMGT), https://www.ebi.ac.uk/ipd/imgt/hla/) to search for unannotated variable and constant region genes (e.g., *IGG*) in the mouse lemur genome (Mmur 3.0, NCBI Annotation Release 101). We then built a custom reference database from these retrieved mouse lemur Ig genes as well as the human IMGT sequences to extract the transcripts and assemble the Ig sequence for each of the 829 B and plasma SS2 cells analyzed using BASIC^130^. Ig sequences obtained by BLAST searches of the transcriptomes from a subset of mouse lemur atlas B cells were added to the custom reference database to further improve alignment. Constant regions from both the heavy and light chain contigs assembled by BASIC were aligned to the reference database using BLAST and the best hit (with at least 80nt of overlap) was used to assign the isotype (i.e. *IGA, IGG, IGM*, or *IGE* for the heavy chain, and *IGK* or *IGL* for the light chain) for each cell. Putative V and J gene families and the *CDR3* sequences from both the heavy and light chain contigs were identified using IgBlast^131^. In some cases, BASIC was not able to generate a contig for the heavy and/or light chain, therefore the isotype was not assigned for these cells (52 and 10 cells for the heavy and light chains, respectively). In cases where BASIC was unable to assemble a single continuous contig from both constant and variable region ends of either the heavy and/or light chain, we submitted to BLAST/IgBLAST the two contigs constructed from each end (94 and 100 cells for heavy and light chains, respectively) or the only constructed contig from one end (147, 1, 28, and 2 cells with only heavy chain constant, heavy chain variable, light chain constant, and light chain variable contigs, respectively). In rare cases, the contigs constructed from the variable and constant region ends from a single cell returned different hits. Constant region isotypes were not assigned for such cells with different BLAST hits (16 and 13 cells for heavy and light chains, respectively). Similarly, V gene families were not assigned for cells with different IgBLAST hits (alignment quality V score >100) from the constant and variable contigs (3 and 59 cells for heavy and light chains, respectively). These discrepancies may be caused by doublets of B cells/plasma cells (though applying the program Scrublet^132^ with default parameters identified only three of the 80 cells with two different BLAST/IgBLAST hits as possible doublets) or more likely, reflect dual expression as more recently appreciated in Shi et al.^133^. CDRH3 lengths were calculated as the number of amino acids between the canonical C at the 5’ end of the sequence and the 3’ sequence WGXG where X is any amino acid. *CDRL3* (including lambda and kappa chains) lengths were calculated as the number of amino acids between the canonical C at the 5’ end of the sequence and the 3’ sequence FGXG or WGXG where X is any amino acid.

All Ig sequences assembled through BASIC were then used to determine the minimum number of constant and variable region alleles in the mouse lemur genome. These sequences were aligned using MAFFT^134^ and then manually corrected using Geneious Prime 2021.1.1 (https://www.geneious.com). Because somatic mutation patterns in *IGV* genes can render a single V gene indistinguishable from separate but closely related alleles, we estimated a minimum number of V genes based on the number of V loci that appeared in the current assembly of the genome. Long read sequences covering this region would help determine the true number of V gene loci.

Clonal lineages were identified as groups (N≥2) of cells within a single individual with identical CDRH3 and CDRL3 lengths, the same light chain isotype and CDR3 identities of at least 80% for both the light (CDRL3) and heavy (CDRH3) chains.

### MHC gene expression analysis

Methods to examine mouse lemur MHC gene structure, extract MHC gene expression from the atlas, and re-annotate the MHC genes are detailed in an accompanying paper^50^. In brief, raw fastq files from 10x scRNA-seq of blood from each of the individuals were mapped against an MHC reference sequence extracted from the Mmur 3.0 genome assembly according to the BAC sequences^135,136^, as well as the known expressed mouse lemur class I Mimu W01-04^137^ using bowtie2^138^. The mapping results were assessed for mismapping of reads, allelic variation and possible presence of additional genes, by manual inspection using the Integrative Genomics Viewer (IGV)^139^ and Geneious Prime 2021.2.2. The fastq files were also ‘probed in silico’ by searching for reads containing sequences specific to the known genes. This confirmed the absence of expression of particular MHC genes. For the analysis of expression levels, a reference specific to each individual was used with bowtie2 to map the reads extracted from the raw fastq files for each tissue. The sequence used was restricted to the final 600 bp comprising exon 5 through the 3’UTR to avoid complications from potential recombinant sequences. Manual inspection of the results from the blood 10x scRNA-seq files was used to determine a mapping quality (MAPQ) threshold for each gene. The Sequence Alignment Map (SAM) file from the mapping was converted to a BAM file and then divided into individual BAM files for each gene. These individual files were then filtered to remove reads below the MAPQ threshold. For the remaining reads, the cell barcode and UMI were counted. The expression level is normalized as read counts/10000 UMIs and then natural log transformed. Counts for each MHC gene (raw and normalized) are available in the metadata for every h5ad file in Figshare (https://figshare.com/projects/Tabula_Microcebus/112227). SS2 data was not used due to limited data available regarding allelic variation and recombination between alleles and/or genes that is prevalent in the MHC. Lacking phase information for the SS2 data made it impossible to accurately assign all sequences to specific genes (only the terminal 3’ 600-650 bp could be assigned with confidence to a particular class I gene). Discarding the upstream information would have biased the expression level results. Thus, we chose to focus on the 10x dataset where the majority of the sequences were obtained from a single region that falls within 600-650 bp of the 3’ end and therefore could be unambiguously assigned to a specific gene.

### Chemokine ligand and receptor expression analysis

A list of human chemokine receptors was compiled from the literature^140,141^ and their cognate ligands were obtained from CellPhoneDB^142^(Table 6). We included the four atypical chemokine receptors which induce G-protein-independent downstream signaling^143^, as well as the chemerin (*RARRES2*) receptors (*CMKLR1, GPR1, CCRL2*) given their established dual role in immune and adipokine chemoattraction^144^. The chemokine *CXCL17*, without a known receptor, was also included. Of the 25 identified receptors, a corresponding lemur ortholog annotated in NCBI was found for all except *CCR2*. Of the 45 cognate ligands, a corresponding lemur ortholog was identified for 32. The log normalized expression level of each of the lemur orthologs across all cell types in the 10x dataset is summarized in Fig. 4a and Fig. S7. Cell type expression levels for each gene were then binarized (i.e., expressed or not expressed) based on absolute and relative thresholds for the purpose of building an interaction network, per below. First an absolute threshold was applied requiring a gene be expressed at non-zero levels in at least 5% of cells and with an average expression level of at least 0.5 across all cells from that cell type. Second, a relative threshold was applied: for each gene, a ceiling expression level was defined as the expression level of the 99th percentile of all cell types that passed the first threshold (to remove outliers with abnormally high expression levels). Cell types with receptor gene average expression level above 5% of the ceiling were deemed to be expressing the receptor. A higher threshold (20%) was applied for ligand genes given that ligands are diffusible and therefore require high levels to be functional.

To build a chemokine interaction network across all cell types in the atlas, connections (edges) were drawn between cell types (nodes) expressing a ligand and cell types expressing the cognate receptor, allowing self-loops where a cell type expressed both ligand and corresponding receptor. Connections between cell types from different organs (other than blood) were excluded given the short effective intercellular communication distances of chemokine signals. Note edges are directed such that cell type A expressing a ligand and cell type B expressing the cognate receptor formed a separate edge from cell type B expressing a ligand and cell type B expressing the cognate receptor. Multiple connections in the same direction between two nodes (i.e., two cell types with more than one receptor-ligand interaction) were counted as a single edge.

Network density was calculated as the number of edges identified divided by the total number of possible edges in the network: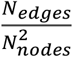. The density was calculated separately for the network of interactions across all cell types in the atlas, only immune cell types, only non-immune cell types, and between immune and non-immune cell types.

### Cross-organ immune cell analysis

Immune subpopulations were identified and annotated by systematic subclustering of the lymphoid and myeloid compartment in each tissue for every individual, then adjusted by inspection using cellxgene after integration of tissues across all individuals to ensure consistency of cell type labeling, as described in the accompanying Tabula Microcebus manuscript^6^. Clusters branching off the main group were labeled with a differentially-expressed gene (e.g., neutrophil (*CCL13*+), neutrophil (*IL18BP+*), B cell (*SOX5*+)) and cells expressing proliferative markers (*MKI67, TOP2A*) were appended ‘PF’ (e.g., B cell (PF)). For macrophages, their identities as tissue-resident macrophages based on published marker genes (see Table 1 in the accompanying Tabula Microcebus manuscript^6^) was indicated by appending the corresponding name (e.g. macrophage (Kupffer cell), macrophage (microglial cell)), given that clear distinction from monocyte-derived macrophages was challenging (with the exception of lung tissue-resident alveolar and monocyte-derived interstitial macrophages which were confidently distinguished based on canonical markers and labeled as such). Identification of differentially-expressed genes for each subpopulation was performed using the Wilcoxon rank-sum test, selecting genes with log fold change >=1 and *p*-value <0.05, after adjustment by the Benjamini-Hochberg correction.

For the cross-organ monocyte/macrophage analysis, all granulocyte-monocyte precursors, monocytes, and macrophages from the atlas were extracted for further analysis. Data were integrated across the four lemur individuals and then across the scRNA-seq methods (10x and SS2 datasets) using the FIRM algorithm^145^ to correct for batch effects. In this integrated UMAP, monocyte populations co-clustered across tissues, whereas macrophage populations were generally separated by tissue. We therefore tried additional FIRM integration across tissues, however tissue-specific separation of macrophage types and L2’s bladder monocytes remained; hence we did not perform tissue-level FIRM integration for the final UMAP to avoid potential computational bias from overcorrection. We then examined expression of known monocyte and macrophage markers reported in the literature as well as the distribution of monocytes and macrophages from each tissue in the FIRM-integrated UMAP (Fig. S9). In addition to the tissue-specific/resident populations highlighted in Fig. S9a, we found that pancreatic and heart macrophages also formed separate populations; however these were excluded from further analysis because they likely resulted from technical issues: the differentially-expressed genes for pancreatic macrophages were found to be broadly expressed in other cell types of the same tissue (signal spreading), and the heart sample had overall fewer transcripts per cell (lower quality).

The neutrophil developmental trajectory was based on embedding of neutrophils in the FIRM-integrated UMAP of the entire atlas (as described in the accompanying manuscript^6^). This resulted in co-clustering of neutrophils by individual and tissue, allowing for recapitulation of the developmental trajectory. We also tried FIRM integration of neutrophils alone (by individual and scRNA-seq method), however that resulted in separation of neutrophils by tissue and individual, which was largely driven by batch effect (no biologically meaningful differentially-expressed genes were identified across most clusters). This suggests that neutrophils are more molecularly homogeneous across tissues compared to other cell types such as macrophages. The trajectory was obtained by an in-house algorithm that detects the density ridge of the cell distribution on the FIRM integrated UMAP embedding, as described in the accompanying Tabula Microcebus manuscript^6^, with the direction of the trajectory assigned manually based on expression of neutrophil maturation markers. Similarly to the neutrophils, the FIRM-integrated UMAP of B cells and plasma cells showed global separation of plasma cells and B cells, however further cell separation was driven by batch effect. In the atlas-wide UMAP, the clear separation between B cells and plasma cells precluded further trajectory analysis.

### Endometrial cancer analysis

Uterine cancer was identified by scRNA-seq (Section 5) and later confirmed by histopathology in both females (L2, L3). Both had metastases, with L2 spread to the lung and L3 to an intra-abdominal lymph node. We analyzed lung metastasis in L2 and primary tumor in L3. L2’s uterus was not analyzed by scRNA-seq because we were unaware of the tumor at the time of tissue harvest. For L3, we were unable to sequence the metastasis given the liquefactive necrotic nature of the tissue.

To compare the novel lung epithelial cell cluster in L2 (retrospectively identified as endometrial tumor cells metastasized from the uterus) with all other cell types of the atlas, we examined the correlation scores (Fig. 6i) and UMAP embedding (Fig. 6h) of their gene expression profiles, by the methods described in the accompanying Tabula Microcebus manuscript^6^. Here, we extracted results of the metastatic tumor cell type. In brief, to calculate the cell type pairwise correlation scores with the metastatic tumor cell type, atlas data was first integrated across individuals, tissues, and scRNA-seq methods (10x and SS2) using FIRM^145^ and FIRM-generated principal component coefficients were calculated for each cell. The coefficients were then averaged across all cells of a cell type and used to calculate the Pearson’s correlation scores between every atlas cell type and the metastatic cells. The lung cell type in L2 that is a hybrid of metastatic and AT2 cells, which could be doublets of the two cell types, (though Scrublet^132^ only identified one of these five cells as a possible doublet) was excluded from the analysis. To generate the cell type UMAP (Fig. 6h), gene expression levels were averaged across cells for each cell type (10x dataset, excluding low quality cell types and ones represented by <4 individual cells, expression levels were normalized (0 to 1 scale) to the maximal value of each gene across all cell types, and the normalized cell type gene expression matrix projected onto a 2D space with cosine distances between pairs of cell types as input.

Differential gene expression analysis was performed on lung metastatic cells versus all other lung epithelial cells and on uterine *FXYD4+ MUC16+* epithelial cells versus all other uterine epithelial cells (10x datasets) using the Wilcoxon rank-sum test (*p*-value <0.05, after adjustment by the Benjamini-Hochberg method), and representative examples selected for Fig. 6j.

### Adipocyte analysis

Adipocytes and adipo-CAR cells were extracted from the FIRM-scaled and integrated data of the entire atlas (10x and SS2 datasets, see accompanying Tabula Microcebus manuscript^6^. The top 3,000 highly variable genes in the FIRM-transformed gene counts table of adipocytes and adipo-CAR cells were selected, and dimensionality reduction by PCA performed (top 13 PCs) to generate a 2D UMAP of adipocytes and adipo-CAR cells. Differential gene expression analysis on the *UCP1*hi versus *UCP1*lo adipocyte populations (L2 and L4, 10x dataset) was performed using Wilcoxon rank-sum test (*p*-value <0.05, after adjustment by the Benjamini-Hochberg method), and representative genes selected for Fig. 6n. Similarly, differential gene expression analysis was performed between the adipocytes of each fat depot of L2 (BAT, MAT, SCAT, GAT; 10x dataset), and the top 10 genes enriched in each depot selected for Fig. S11e.

Most of the adipocytes in the atlas were isolated from fat depots, for which the tissue dissociation protocol was designed to enrich for the stromal vascular fraction (SVF) and exclude adipocytes (Supplementary Methods in accompanying Tabula Microcebus manuscript^6^). Most were from L2 (Fig. S11a),whose adipocytes in fat depots surrounding several tissues (e.g., kidney, spine, uterus) were generally small, predominantly multilocular, densely stained, and mitochondrial rich (Fig. S11d), features of brown or beige adipocytes in human and mouse, and intermingled with small, unilocular adipocytes with a single lipid droplet, which resemble white adipocytes. In contrast, L3 and L4 adipocytes were generally larger and most were unilocular (Fig. S11d). These may be harder to capture by current scRNA-seq protocols so may have contributed to the lower yield of adipocytes for L3 and L4.

### Identification of PS genes and analysis of their expression patterns in lemur and human

The list of human and lemur orthologous genes with no corresponding mouse ortholog was compiled by merging human, mouse lemur, and mouse homology assignments from NCBI, Ensembl, and Mouse Genome Informatics (MGI) databases by a similar method used to compile the list of one-to-one-to-one gene orthologs for the comparison of cell types across the three species in the accompanying Tabula Microcebus manuscript^6^. We began by compiling all human protein-coding genes annotated in NCBI (taxonomy ID: 9606), then merged the corresponding mouse lemur and mouse orthologs from NCBI (gene_info.gz and gene_orthologs.gz from https://ftp.ncbi.nlm.nih.gov/gene/DATA/, February 2020/August 2023). We next added Ensembl gene ID numbers, gene names, and lemur/mouse ortholog assignments from Ensembl Biomart (Ensembl Genes version 99, February 2020), using the Ensembl gene ID (variable ‘Gene_stable_ID’) for each NCBI gene ID (variable ‘NCBI_gene_ID ‘) in Ensembl Biomart. MGI mouse gene ID numbers, gene names, and ortholog assignments (none provided for lemur) from Jackson Laboratory (HOM_MouseHumanSequence.rpt from http://www.informatics.jax.org/downloads/reports/, Feb 2020) were added using the MGI homology ID (variable ‘HomoloGene_ID’) attributed to each NCBI gene ID (variable ‘EntrezGene_ID’) in the MGI database. The Online Mendelian Inheritance of Man (OMIM) genetic disorder phenotype associated with each human gene (genemap2.txt from https://www.omim.org/downloads January 2022, variable ‘Phenotypes’) was added using the gene name (variable ‘Approved_Gene_Symbol’) in the OMIM database.

A human gene was identified as sharing an ortholog with lemur if at least one such assignment was made by either Ensembl or NCBI, and/or as sharing an ortholog with mouse if at least one such assignment was made by NCBI, Ensembl, or MGI. This resulted in 539 human genes with an assigned lemur, but no mouse ortholog (i.e., PS genes), which corresponded to 388 unique lemur Ensembl gene IDs and to 425 unique lemur NCBI genes IDs (not all orthologs are annotated in both NCBI and Ensembl). Note that gene orthology assignments from NCBI, Ensembl, and MGI are being updated periodically, thus these numbers may change in the future. Transcripts were detected for 401 of the 425 PS NCBI-annotated genes, and their expression patterns across all lemur atlas cell types (10x dataset) were visualized in heat maps and dot plots and qualitatively categorized by whether their expression was enriched (higher or restricted expression) or depleted in one or more tissues/organs or compartments (see Table 8, Fig. S12). Expressed genes that did not show any of these expression patterns were categorized as ‘not enriched in any category’.

Gene set enrichment analysis of the 539 PS genes was performed using gprofiler2 in R^146^, searching for overrepresented gene sets (relative to all human annotated genes) in Gene Ontology terms, biological pathways, regulatory DNA elements, human disease gene annotations, and protein-protein interaction networks, using default parameters (e.g. user_threshold = 0.05, correction_method = “g_SCS”).

We further analyzed evolutionary conservation in the expression patterns of the PS genes that have one-to-one orthology mapping between human and lemur (Table 8). Analysis followed a similar pipeline as described in the accompanying Tabula Microcebus manuscript^6^, where we compared across human, lemur, mouse, and macaque using one-to-one orthologs (i.e., not including any PS genes). We analyzed cells from the lung, skeletal muscle, liver (epithelial cell only), testis (germ cell only), as well as bone marrow and spleen (immune cells only). To unify cell type annotation, data of different species were integrated by Portal using ∼15,000 one-to-one orthologs, and cells were reannotated for consistent designation across all species. Here, we applied the same cross-species cell type annotation and compared between human and lemur only, with lemur data from the Tabula Microcebus study^6^ and human data from the Human Lung Cell Atlas^140^ (lung), Shami et al.^147^ (testis) and Tabula Sapiens^124^ (rest tissues).

With manual curations, we identified a total of 398 PS genes with one-to-one orthology mapping between human and lemur. Note that NCBI and Ensembl occasionally have inconsistent orthology assignments. For example, one database may assign a one-to-one mapping whereas the other database may assign a one-to-many mapping. In such cases, we prioritized NCBI mapping but also maximized coverage by retaining the orthologs with identical gene symbols or description in both species. Next, we analyzed 346 of the PS genes that were reported in all scRNA-seq datasets described above. Because the number of annotated genes were different in human and lemur, we normalized the transcript counts of the PS genes against the background of all one-to-one orthologs, and then log transformed the expression levels (i.e., ln(1+UP10K)). For each PS gene, mean expression (*E*_max_) in the maximally expressed cell type in each species was quantified. Next, we filtered for PS genes with notable expression across the analyzed cell types, requiring *E*_max_>0.5 in each species or *E*_max_>0.1 in each species and *E*_max_>1.5 in at least one species. This resulted in a total of 93 PS genes for which we quantified their expression pattern similarity between human and lemur. Mean cell type expression levels across the orthologous cell types were normalized by *E*_max_ of the same species and Pearson’s correlation coefficients between human and lemur were calculated and reported in Table 8. Majority (55%, 51/93) of the analyzed PS genes had a correlation coefficient above 0.5, indicating conservation in their expression patterns between lemur and human.

### Analysis of natural mutations

We performed whole genome sequencing (WGS) for each of the four atlas lemurs (L1-4) along with 31 additional lemurs originating from the same lab colony (median 54x coverage). Methods for the WGS and analysis pipeline are detailed in an accompanying manuscript (Chang et al., accompanying manuscript). Briefly, genomic DNA libraries were generated by Tn5-based tagmentation, and indexed and PCR-amplified for 150 bp paired-end short-read Illumina sequencing. Sequencing reads were aligned to the current Mmur 3.0 genome assembly (NCBI RefSeq assembly accession: GCF_000165445.2) and germline variants identified using the Sentieon® DNAseq® workflow. Variants were annotated and filtered, and functional impact predictions made using the SnpEff & SnpSift toolbox. This resulted in ∼45 million total variants across all 35 lemurs. To identify functionally-significant rare nonsense variants relevant for this study, we first filtered for 3 criteria: 1) allele frequency less than 0.5; 2) base call quality greater than 99.9%; 3) homozygous or heterozygous variants present in at least one of the atlas lemurs (L1-4). This resulted in 6905 variants. Next, we refined this list by filtering for variants where their respective genotypes were identified in all sequenced animals, focusing on nonsense variants by looking at those predicted to cause frameshift mutations, alterations in the stop codon, as well as variants computationally predicted to cause NMD. This narrowed our final list to 713 unique variants found in 713 genes (1 per gene).

To analyze the transcriptional impact of these nonsense variants, we compared cell type-specific gene expression of the affected gene within the four atlas lemurs. We prioritized genes that were abundantly expressed and potentially functionally-important (e.g., absent in mouse). Cell-type specific scRNA-seq reads were identified by their 10x barcodes, parsed from the original post-alignment BAM files for each atlas lemur and counted using Samtools (version 1.16.1) across the respective gene. This enabled discernment of the number of reads with the reference allele versus those with the alternative allele at the variant locus, along with the total number of reads mapping to the gene. Quantifying and analyzing the differing allelic expression patterns of these genes, in the presence and absence of the variant, allowed us to identify nonsense variants linked to significant reductions in gene expression.

## TABLES

**Table 1. Differentially-expressed-uTARs and their homologous genes and cellular expression levels**

Table 1a (DE uTAR characteristics) lists the 4003 unique differentially-expressed (DE) uTARs detected in the atlas. Columns include uTAR identifier (unique uTAR ID that appends genomic position, strandedness (+/- strand), and total count of aligned reads followed by a “-0”, indicating the TAR region is a uTAR), fasta (uTAR location in the genome in the FASTA format, i.e., chromosome ID followed by the start and end locations), fastaPeak (full width at half maximum region around the absolute peak in coverage after Gaussian smoothing within the uTAR location), tissue_cell type_all (list of cell types that the uTAR is differentially-expressed in each atlas tissue), num_tissue_cell type (number of cell types the uTAR is differentially expressed), tissue_cell type_top (the cell type with the most significant *p*-value in the differential expression test), avg_logFC (log(e) fold increase of the expression in the indicated top cell type compared to other 10x cell types in the same tissue), p_val_adj (*p*-value of enriched expression in the indicated top cell type, adjusted by Bonferroni correction), pct.1 (percent of cells in the indicated top cell type expressing the uTAR), pct.2 (proportion of other 10x cells in the same tissue expressing the uTAR), blastShort (most frequent significant BLASTn result for the entire length of the uTAR), blastShort_peak (most frequent significant BLASTn result for the peak region in the uTAR). The table indicates whether the DE-uTAR nucleotide sequence has homology with a human gene that has no annotated mouse or lemur orthologs (is_H=TRUE) suggesting it is a PS gene missed in the current lemur NCBI annotation; or with a human gene that has a mouse ortholog and no annotated lemur orthologs (is_HM=TRUE) suggesting it is a conserved gene across all three species missed in the current lemur annotation. The table also highlights DE-uTARs that overlap in genomic position with the mouse lemur Ensembl annotation but that is missing in NCBI (is_EnsemblOnly=TRUE).

Table 1b (uTAR cellular expression) lists the identity of each cell (cell ID, tissue source, individual source, cell ontology, and free annotation, as described in https://tabula-microcebus.ds.czbiohub.org/whereisthedata) and its TAR characteristics including for each cell the percentages of reads (UMIs) detected by the TAR analysis that are annotated TARs (percent_aTAR) as well as unannotated TARs (percent_uTAR) separated by differentially-expressed uTARs (percent_uTAR_DE) and not differentially-expressed uTARs (percent_uTAR_nonDE).

Table 1c (cell x uTAR count table) lists the UMI count matrix of the DE-uTARs across all 10x cells, with each entry being a non-zero element of the matrix followed by the corresponding cell index (index_cell), uTAR index (index_uTAR), and number of UMI (n_reads). The uTAR and cell indexes are as indicated in Table 1a and 1b, respectively.

**Table 2. Spice junctions and categories identified by the SICILIAN analysis**

Table lists all splice junctions detected by SICILIAN. Each row corresponds to a distinct junction and columns are: junction_ID (unique name for each junction providing chromosome name, gene name, coordinate, and strand orientation for the 5’ and 3’ sides of the junction), num_reads_total (total number of reads mapped to the junction across all 10x and SS2 datasets), chromosome_refseq_ID (RefSeq ID for the chromosome), chromosome_name (chromosome name), junction_5SS (5’ splice site coordinate), junction_3SS (3’ splice site coordinate), gene_5SS (gene name for the 5’ side of the junctions), gene_3SS (gene name annotation for the 3’ side of the junctions), strand (strand orientation), conservation (conservation status with human and/or mouse as defined in Fig. 2d), category (annotation category as defined in Fig. 2b).

**Table 3. Genes differentially spliced across cell types**

Table of genes with statistically significant splicing differences across cell types in the same tissue for at least two lemurs, detected by MANOVA test using the 10x atlas dataset. Each row lists a unique combination of gene, tissue source, and lemur individual source. Additional columns include pval_zscore_adj (the p-value resulting from the MANOVA test) and num_cell_types (number of cell types analyzed with the MANOVA test in the given tissue and individual for this gene).

**Table 4. Expression homolog triads of human, lemur, and mouse detected by SAMap**

Table lists expression homolog triads detected by SAMap analysis separately on the lung and skeletal muscle cross-species datasets (listed in column “Dataset”). Each row represents an expression triad with its type (‘Type_expression_homolog’), gene symbols in the three species (‘huGene’, ‘leGene’, ‘msGene’) including its species prefix, status of lemur gene annotation in NCBI (‘is_leGene_Named’), number of gene pairs in the triads that are orthologs (‘N_ortholog_pair’) or non-orthologs (‘N_non-ortholog_pair’), pairwise orthology relationship between species (‘huleOrthology’, ‘humsOrthology’, ‘lemsOrthology’), expression pattern correlation score between gene pairs (‘huleCorr’, ‘humsCorr’, ‘lemsCorr’), and the cell types with selective expression of the gene in respective species (‘huCellType’, ‘leCellType’, ‘msCellType’).

**Table 5. Expression of immunoglobulin isotypes in atlas B cells and plasma cells**

Table lists atlas B cells and plasma cells (SS2 dataset) with the immunoglobulin isotype identified by the scRNA-seq profile of each cell, including the expressed constant region isotype (heavy_constant_isotype, light_constant_isotype) and variable region family member (heavy_variable_family, light_variable_family) of heavy and light chains, as well as the complementarity-determining region (CDR3) amino acid sequence and length of the heavy (CDRH3_aa_sequence, CDRH3_aa_length) and light chains (CDRL3_aa_sequence, CDRL3_aa_length). Blank entries in these columns indicate cells for which the isotype could not be assigned (see Methods). Pairs or triplets of cells identified as part of the same clonal lineage, based on their CDRH3 and CDRL3 sequence similarity, are given the same number in column ‘clone_number’.

**Table 6. Chemokine receptor and cognate ligand genes**

Table lists the analyzed lemur orthologs designated by NCBI (‘receptor_lemur_ortholog’, ‘ligand_lemur_ortholog’) of the 25 human chemokine receptors and their cognate ligands, retrieved from CellPhoneDB (March 2024). The source of each curated ligand-receptor interaction is given under ‘cellphoneDB_annotation_strategy’, ‘cellphoneDB_curator’, and ‘cellphoneDB_source’ (CellphoneDB output, ‘curated’ indicates it was annotated by CellPhoneDB developers), and the type of signaling interaction is listed under ‘cellphoneDB_classification’. For receptors with multiple ligands, the primary ligand(s) (when known) are noted under ‘comments’.

**Table 7. Human genes and their lemur and mouse orthology assignments**

Table 7a (Summary of human orthology assignments) summarizes the orthology status of all human protein-coding genes annotated in NCBI (19,966), beginning with the NCBI gene ID, synonym, gene description, and gene type. The presence of corresponding mouse lemur orthologs assigned by NCBI and/or Ensembl (as detailed in Methods) is indicated in columns ‘HumanLemur_NCBI_HomologyType’ and ‘HumanLemur_Ensembl_HomologyType’, respectively. Similar columns are provided for corresponding mouse orthologs assigned by NCBI, Ensembl, and/or MGI. For every human gene, column ‘any_Lemur_homology’ summarizes if there is at least one lemur ortholog assigned by NCBI and/or Ensembl and column ‘any_Mouse_homology’ summarizes if there is at least one mouse ortholog assigned by NCBI, Ensembl, and/or MGI. Human genes that are primate-selective/mouse-absent (539) are identified in the column ‘HumanLemurOrthogs_NotMouse’ (1: ‘any_Lemur_homology’ = yes and ‘‘any_Mouse_homology’ = no, 0 for all other entries).

Table 7b (All human orthology assignments) lists all human protein-coding genes annotated in NCBI (19,966) with corresponding NCBI gene ID, synonym, external database cross references (dbXrefs), chromosome location, map location, gene description, gene type, as well as corresponding Ensembl gene ID, gene name and OMIM associated phenotype description. The presence of corresponding mouse lemur orthologs assigned by NCBI and/or Ensembl (as detailed in Methods) is indicated in columns ‘HumanLemur_NCBI_HomologyType’ and ‘HumanLemur_Ensembl_HomologyType’, respectively, with the corresponding NCBI gene ID, gene name, and synonym as well as Ensembl gene ID and gene name listed for lemur genes with an assigned ortholog in either of these databases. Similar columns are provided for the corresponding mouse orthologs assigned by NCBI, Ensembl, and/or MGI. Rows are duplicated for each human NCBI gene with more than one corresponding Ensembl gene, or that is assigned to more than one human/mouse ortholog.

**Table 8. PS genes and their expression patterns in the mouse lemur and human**

List of mouse lemur PS genes annotated in NCBI (425), starting with the corresponding NCBI gene ID, synonym, external database cross references (dbXrefs), chromosome location, gene type, gene description, and, if present, the associate human OMIM genetic disorder phenotypes. For each gene, the compartments and tissues that have enriched expression of the gene are listed in columns ‘Lemur_enriched_compartment’ and ‘Lemur_enriched_tissue’, and summarized in column ‘Lemur_expression_pattern_summary’. Cell types showing selective expression in the enriched compartments/tissues are noted in column ‘Lemur_enriched_celltype’. Compartments and tissues that have selectively depleted expression are indicated by brackets. The number of lemur cell types (10x dataset, excluding cells labeled as mix, doublets or low quality) with mean expression of the gene greater than zero (column ‘N_lemur_celltypes_wMeanExpGreaterThan0’), as well as the number of cell types with mean expression of the gene greater than zero and expressed in at least 1% of the cells in that cell type (‘N_lemur_celltypes_wMeanExpGreaterThan0_ExpIn1PCTOfCells’) are provided. The order of genes appearing in Fig. S12a-b heat maps are provided in column ‘Gene_order’. PS genes (aTARs) found using the TAR analysis pipeline (see Methods - uTAR analysis) are indicated in the ‘Gene_inTARanalysis’ column. The next three columns indicate whether the PS gene has one-to-one orthology mapping (‘HL_Orthology_one2one’), and if yes the corresponding human ortholog gene ID and symbol (‘Human_NCBI_GeneID’, ‘Human_NCBI_GeneName’). Finally, the last two columns indicate whether the PS gene was also present in the human datasets used for the cross species analysis (‘scRNAseq_reported’) and the correlation coefficient of its human vs. lemur cell type expression patterns (‘HL_Expression_R’).

**Table 9. Gene sets enriched in PS genes**

Gene set enrichment analysis (gprofiler2^146^ in R) for the 539 PS genes. Columns detailed in https://cran.r-project.org/web/packages/gprofiler2/vignettes/gprofiler2.html.

## SUPPLEMENTARY MATERIALS

### Legends for Supplementary Figures

**Figure S1. Sequence identity of the human genes to lemur versus that of rodents or higher monkeys**.

Scatter plot comparing pairwise gene sequence identity between human and lemur vs. human and mouse (a), human and rats (b), human and rhesus macaque (c), human and crab-eating macaque (d), or human and marmoset (e). Shown in each panel are genes with ortholog in each of the three species compared, assigned by the Ensembl homology database. Red lines indicate 1-1 relationship. The percentage and numbers indicated in each panel represent the genes on the lower right side of the red curve, i.e., with higher sequence identity between human and lemur than between human and the other species.

**Figure S2. uTAR analysis enhances mouse lemur genome annotations and uncovers potentially novel transcripts**.

**a**. Box plot of average silhouette coefficient values of the atlas datasets (separated by tissue and lemur individual, and sequencing channel) based on expression of annotated genes, aTARs, or uTARs. For each dataset, cells were grouped by their cell type designation and their silhouette value was calculated using the cell-cell distances in the 2D UMAP embeddings of either the gene, aTAR, or uTAR-expression space, and then averaged across all cells of the dataset (see Methods). Box, mean±s.d.; red triangles, an example dataset of L4 colon (10x) as shown in panel b. Note the positive uTAR-based silhouette values in most datasets, supporting effective clustering of cells according to cell types by uTARs alone.

**b**. UMAP visualization of L4 colon cells (10x dataset) embedded according to the expression of the annotated genes (top) or uTARs (bottom). Datapoints (cells) are colored by cell type designation (as in c).

**c**. Dot plot showing average expression of selected DE uTAR (labeled by identified sequence homology) across L4 colon cell types. Identical gene names indicate multiple uTARs aligned to the same gene in another species.

**d**. Expression of selected sperm cell type markers in germ cells from testis (L4, 10x dataset) ordered by the pseudotime developmental trajectory calculated by uTAR expression (as in Fig. 1d-g).

**e**. Percentage of genes detected by the TAR analysis (for indicated gene categories) as a function of the filtering threshold used to define cell type selective expression (i.e. TAR expression in any cell type ≥ e^threshold times that of the average of other cell types). Gene categories examined include: the top 100 (black), 2000 (dark gray), and 5000 (light gray) variably-expressed genes annotated in Mmur 3.0 NCBI annotation, all genes (blue), PS genes (yellow), and genes annotated in Mmur 3.0 Ensembl annotations but missing from NCBI (green).

**f**. Venn diagram of 4003 DE-uTARs with sequence homology to coding regions (>1 hit by DIAMOND blastp analysis) and/or non-coding regions (>1 hit by Infernal cmscan analysis), according to Nf-predictorthologs analysis.

**Figure S3. SICILIAN analysis reveals cell type specific alternative splicing**

Barplots showing percent of indicated splice isoforms of *MYL6* (a), *CAST* (b), *MLF1* (c), *TPD52L2* (d), *FAM92A* (e), and *CHPT1* (f) as in Fig. 2e-g but across all atlas cell types that have at least 10 cells with spliced transcripts of the gene and a total of ≥300 reads of spliced transcripts of the gene. Exceptions were made to spermatogonia (testes 250) of *TPD52L2* (1 cell, 8 reads), *FAM92A* (7 cells, 21 reads), and *CHPT1* (34 cells, 112 reads) and diplotene spermatocyte (testes 253) of *FAM92A* (37 cells, 50 reads), and *CHPT1* (88 cells, 148 reads). Cell types were labeled by their tissue source and designation number^6^, and colored by compartment. In the transcript annotations at top, splice isoforms are labeled with the NCBI Refseq ID and colored to correspond with barplot, with exons affected by alternative splicing shown in red.

Note for *CHPT1* (f), the two transcripts in dashed box cannot be distinguished using the atlas data due to insufficient reads covering dashed region of the transcript so reads of the two isoforms are combined in the analysis. Note *MYL6* and *CAST* show compartment-dependent patterns of splicing, and *MLF1, TPD52L2, FAM92A*, and *CHPT1* are differentially spliced in male germ cells (testes #250-256).

**Figure S4. Additional examples of expression homolog triads of the “named ortholog” type**

Dot plots as in Fig. 3d showing expression of additional expression homolog triads of the “named ortholog” type, identified from the lung (a) or skeletal muscle (b) datasets.

**Figure S5. Structure and global expression pattern of mouse lemur MHC class I and II genes**

**a**. Schematic of mouse lemur MHC locus on chromosome (chr.) 6. Filled rectangles, expressed genes; open rectangles, pseudogenes. Numbers below *DRB, DQA*, and *DQB* indicate the number of genes annotated by NCBI, though Guethlein et al.^50^ suggest a single gene for each family. Dashed lines, extended areas in genome assembly. g1, LOC105855356 in NCBI; g2, LOC105855357; g3, LOC105858107.

**b**. Schematic of MHC class I gene locus on mouse lemur chromosome 20. Top line, gene order as annotated by NCBI. The three genomic segments are separated by gaps (slanted lines) in Mmur 3.0 genome assembly. Second line, proposed reorganization of assembly based on rearrangement of three segments to match gene order of a sequenced BAC^136^ (third line; note gene content in Mmur 3.0 assembly and BAC differ due to haplotype variability in the individuals sequenced). Dashed lines connect corresponding genes. Fourth line, proposed revision of structure and annotation of this locus based on above and the expression pattern of these genes in the mouse lemur cell atlas. Dashed boxes, genes varying in either presence, copy number or expression status. *W03* and *W04* are sequences derived from the original study on the lemur MHC^137^ based on sequence similarity, *168, W03* and *W04* could represent divergent allelic variants (or separate genes). *202* and *202P* are a pair of phylogenetically related genes annotated by NCBI, possibly representing a recent duplication; however, there was no evidence supporting them as separate genes in atlas expression data, suggesting *202P* is a pseudogene or assembly error. g4, LOC105855949 in NCBI; g5, LOC105855951; g6, LOC105870766; g7, LOC105870764; g8, LOC105870765; g9, LOC105870769; g10, LOC105870767; g11, LOC105870762.

**c**. Number of putative alleles for each MHC gene in the four lemurs (L1-4), and number of genes annotated in the BAC and by NCBI. Note some class I sequences differ only by a few base pairs among haplotypes^50^, therefore the number of alleles in the panel represents our best estimate but may be inexact due to sequencing errors or other technical artifacts. Genes that exhibit copy number variation (C) or have at least one allele that is a pseudogene (P) are indicated below each panel. ?, insufficient number of reads for a reliable allele count. NA, gene absent from BAC sequence.

**d**. Comparison of the MHC class I and class II regions of mouse lemur with other primates and the mouse^48,135,136,165–167^. (chr.), chromosome on which the MHC regions are located. Dark blue, classical MHC class I genes (high, widespread expression); light blue, non-classical MHC class I genes (lower expression and/or tissue-specific expression), orange, MHC class II A genes; red, MHC class II B genes. Gray dashed box, genes with haplotypic variability in gene number or expression status. Cyan dashed boxes enclose mouse MHC haplotype T, M, and Q regions with expanded gene families.

**e**. Schematic representation of the three *Mimu-DRB* genes in current mouse lemur genome assembly (Mmur 3.0). *DRB1-10* is predicted to encode a full length DRB1 polypeptide, whereas *DRB1-1* and *DRB1-4* are incomplete and contain non-MHC sequences (gray shading). Note that the MHC sequences in *DRB1-1* and *DRB1-4* encode a functional DRB polypeptide, suggesting that these sequences were misassembled and belong together to form a second complete *DRB1* allele^50^. Thin vertical lines, positions that differ between the three sequences.

**f**. Sina plot showing summed expression of classical class I, non-classical class I, and class II MHC genes, respectively and averaged across individual cell types.

**g**. Sina plot showing ratio of summed non-classical to summed classical MHC class I expression.

Note consistently lower levels of non-classical vs. classical MHC class I expression across almost all atlas cell types (dots), except a few highlighted cell types in the neural and germ compartments.

**h**. Dot plot of average expression of each MHC gene across all molecular cell types in the 10x dataset, combining all individuals. Cell types are grouped by compartment and numbered as in accompanying Tabula Microcebus manuscript^6^.

**Figure S6. Immunoglobulin CDRH3 length and clonal analysis**

**a**. CDRH3 lengths (number of amino acids, aa) for L2 and L4 (SS2 data), as well as healthy humans and lab mice. N, number of cells for mouse lemurs and number of unique clones for humans and mice, including all isotypes. Human and mouse data courtesy of Scott Boyd’s group and Tho Pham^168,169^. CDRH3 lengths over 30 were rare in human and mouse and did not occur in the mouse lemurs analyzed and are therefore not displayed. Note CDRH3 length in lemur is generally shorter than in human and more comparable to that of mouse. Though it is known that the CDRH3 region length also varies with age, disease state, and B cell maturity to affect antigen-binding affinity, the functional relevance of inter-species variation remains unsettled^170^.

**b**. B and plasma cell clones identified by their CDRH3 sequence. Each clone is represented as a circle, with the outline color indicating the lemur from which the clone was found and the fill color indicating the heavy chain isotype(s) of the clone (colors as in Fig. 1h-i). All clones consisted of two cells except the larger circle (in spleen) which is a three-cell clone. Circles on an organ indicate the organ in which the constituent cells of the clone were found, and circles between organs indicate the two organs in which the constituent cells were found. Organ images were created by Mikael Häggström and Madhero88 and are in the public domain.

**Figure S7. Expression of chemokines and receptors across atlas cell types**

**a**. Heat map showing percent of chemokine ligands (n=32), typical receptors (20), and atypical receptors (4) expressed in each cell type separated by tissue (10x dataset). Cell types are ordered by their designation number^6^.

**b**. Rank of cell types based on the percent of chemokine ligands (left), typical receptors (center), and atypical receptors (right) expressed. Cell types (dots) are colored by tissue compartment.

**c**. Dot plot showing average expression of selected chemokine ligands and receptors across all epithelial cell types in the atlas (10x dataset), as formatted in Fig. 4a.

**d**. Global cell-cell interaction map based on chemokine signaling. Cell types are arranged by their designation number with the colored bars indicating the tissue compartment. Gray and orange curves link the ligand-expressing cell types to the cell types expressing the cognate receptor; Gray, all chemokine ligand-receptor pairs; Orange, an example chemokine ligand-receptor pair, CXCL12→CXCR4, where the ligand CXCL12-expressing cell types were additionally labeled yellow and receptor CXCR4-expressing cell types were indicated with black bars.

**e**. Lung-specific cell-cell interaction maps based on chemokine signaling, as in panel d. Lung and blood cell types were included. Each panel separately plots specific chemokine ligand-receptor pairs (indicated on the lower left) with curves connecting ligand-expressing cells to the receptor-expressing cells (black bars). ^*^, presumed activated immune cell types (see Section 4 and Supplementary Results).

**f**. Dot plot showing average expression across all cell types in atlas (10x dataset) of all chemokine receptors and cognate ligands analyzed (Table 6). Cell types are ordered by compartment, then cell type designation number^6^, then tissue. [], description of gene identified by NCBI as a gene locus ([CCL4], LOC105881712; [CCL8], LOC105885739; [CCL11], LOC105885804; [CCL13], LOC105859268; [CXCL4], LOC105860501).

**Figure S8. FIRM-integrated UMAP of all atlas immune cells and further characterization of neutrophils and B lymphocytes**

**a-d**. UMAP of all atlas immune cells as in Fig. 5a but colored by proliferation state (a), scRNA-seq method (b), individual (c), and tissue of origin (d).

**e**. UMAPs of lung neutrophils (10x dataset) of the indicated individuals, with cells colored by their molecular cell type designation. Note the two subtypes of activated neutrophils (*CCL13+* and *IL18BP+*) cluster separately from the main population of lung neutrophils (non-activated) in both L1 and L2.

**f**. Heatmap showing relative expression of the indicated differentially-expressed genes in each of the activated neutrophil subtypes. Bars at top show, for each neutrophil, its molecular cell type designation (bottom bar), individual lemur source (middle bar), and tissue source (top set of bars). Note both activated neutrophils were found in more than one individual and from multiple tissues. Single cell expression levels are normalized to the stable maximum (99.5 percentile) of that gene across cells along the neutrophil trajectory. The mature (non-activated) neutrophils are highly abundant so ones shown are 1/5 subsamples of non-activated neutrophils at late stage (>0.7) of the pseudotime trajectory (corresponding approximately to cells within blue stripes in Fig. 5d). [], description of gene identified by NCBI as a gene locus ([IFITM3L], LOC105874071; [Uncharacterized 1], LOC105867541; [CCL2L], LOC105859340 and LOC105885684; [CCL13], LOC105859268; [Uncharacterized 2], LOC105856756; [CCL3L], LOC105881608).

**g**. Dot plot showing average expression in the indicated B lymphocyte lineage cells in atlas (top) of marker genes for B cells, plasma cells, and top differentially-expressed genes in the *SOX5*+ B cell population compared to other B cells. Lemur B and plasma cells in the atlas appeared mostly molecularly homogenous, except for this population of *SOX5+* B cell population identified in pancreas (with nearby lymph nodes).

**Figure S9. Analysis of the monocyte and macrophage lineage development and tissue specialization**

**a**. UMAP of atlas immune cells of monocyte and macrophage lineage including progenitors, integrated by FIRM across different tissues and individuals (10x and SS2 datasets), by either major developmental stages (left) or major groups of tissue-specific/resident macrophages (right). Also note a separately-clustered population of activated monocytes (L2 bladder and perigonadal fat) indicated in the right panel (see also panel d).

**b**. Dot plot showing average expression of classical marker genes for hematopoietic precursors, granulocyte-monocyte progenitors, monocytes, macrophage, and tissue-resident macrophage markers across the atlas immune cells in the monocyte/macrophage lineage. Note some markers are shared between multiple cell types (see Table 1 in the accompanying Tabula Microcebus manuscript^6^). [], description of gene identified by NCBI as a gene locus ([CD14L], LOC105862649; [CD163], LOC105869074; [CD209L], LOC105885453).

**c**. Dot plot showing average expression of differentially-expressed genes of the indicated tissue-resident macrophage populations in the atlas, compared to all other macrophage populations. [], description of gene identified by NCBI as a gene locus ([LOC105878881], CD300C; [LOC105876782], HLA-DRB1L; [LOC105869752], HLA-DQA2; [LOC105869753], CCL3L; [LOC105872655], TLL; [LOC105866341], SIGLEC8; [LOC105882132], SIGLEC7; [LOC105873562], FCGR3AL; [LOC105871481], KRT76L; [LOC105863040], PRXL2A).

**d**. Dot plot showing expression of the top differentially-expressed genes of the separately-clustered monocytes from L2 bladder and perigonadal fat, compared to the other atlas monocytes. [], description of gene identified by NCBI as a gene locus ([Uncharacterized 1], LOC105869025).

**e**. UMAP as in panel a showing lemur expression of classical and non-classical monocyte markers as well as M1 and M2 macrophage markers for human and mouse. Human classical monocyte (*CD14++ CD16-*) markers: *CD14, CCR1, CCR2, CXCR1, CD11B/ITGAM, SELL/CD62L, CD163*; non-classical monocyte (*CD14+ CD16++*) markers: *CD16/FCGR3A, CD11C/ITGAX, CD64/FCGR1A, CX3CR1*; intermediate monocytes express intermediate levels of these markers^171–173^. Mouse pro-inflammatory monocyte (*Ly6c*hi) markers: *Ly6C, Ly6G/Gr-1, Ccr2, Sell/Cd62l*; patrolling monocyte (*Ly6c*lo) markers: *Cx3cr1* and *Spn/Cd43*^*171,173*^. In both human and mouse, M1 macrophage markers: *IL12A/B, IL23A, IL1B, IL6, TNF*; M2 macrophage markers: *IL10* and *TGFB1*^*171,174*^. Note lemur monocytes and macrophages either showed no clear pattern of these markers (plotted) or the respective genes remain unannotated in the lemur genome (i.e., *CCR2, CD64/FCGR1A, CD11C/ITGAX, Ly6c1, Ly6c2, Ly6G/Gr-1, CD43/SPN*). [], description of gene identified by NCBI as a gene locus ([CD14L1], LOC105862649; [CD14L2], LOC105862489; [CD16L], LOC105873562; [CD163], LOC105869074).

**f**. UMAP as in panel a with cells colored by lemur individual (left), scRNA-seq method (center), and tissue source (right).

**f**. UMAP as in panel a separately highlighting cells from each tissue, with cells colored by tissue (left), individual source (center), and molecular cell type designation (right).

**Figure S10. Further characterization of endometrial tumor metastasis cells in lung and uterine *FXYD4+ MUC16+* non-ciliated epithelial cells**

**a-b**. UMAPs of lung (a) and uterine (b) cells from atlas, as in Fig. 6f-g, with cells colored by epithelial cell type designation. Cells in gray are non-epithelial cells.

**c-f**. Same UMAPs of lung (c, e) and uterus (d, f) with cells colored to show expression of *OXTR* (c, d) and *MUC16* (e, f).

**Figure S11. Mouse lemur adipocytes and expression patterns of adipocyte differentially-expressed genes, selected adipokines, and their receptors**

**a**. FIRM-integrated UMAP of adipocytes and adipo-CAR cells (10x and SS2 datasets) as in Fig. 6k-l with cells colored by cell type designation, scRNA-seq method, individual lemur source, and tissue source, respectively.

**b**. UMAP as above colored by expression level of indicated adipocyte markers (*ADIPOQ, CIDEC*) and example differentially-expressed genes in *UCP1*lo (*CHIT1, APOE*) and *UCP1*hi (*FABP3, KCNK3*) adipocytes.

**c**. UMAP as above colored by the number of scRNA-seq reads per cell (UMIs, 10x dataset; transcripts, SS2 dataset, left) and number of genes detected per cell (right). Note the heterogeneity of *UCP1*lo population, which forms two subclusters distinguished by total read per cell and genes detected per cell, but not by any biologically significant differentially-expressed genes.

**d**. Micrographs of H&E-stained sections of fat tissues from L2 (left) and L4 (right) that are near the kidney (top) and paraspinal muscle (bottom). Scale bar, 50 μm (all panels). Full micrographs available online at the Tabula Microcebus web portal.

**e**. Dot plot of average expression of the top 10 differentially-expressed genes in each of the four fat depots: BAT, interscapular brown adipose tissue; GAT, perigonadal; MAT, mesenchymal; SCAT, subcutaneous (L2, 10x dataset). [], description of gene identified by NCBI as a gene locus ([Uncharacterized 1], LOC105856764; [Uncharacterized 2], LOC105867540; [COX7A1], LOC105876884; [Uncharacterized 3], LOC105867541; [MT2A], LOC105866476; [MT1E], LOC105866554; [CTRB1L], LOC105875474; [MAGEB16L], LOC105877758; [PRSS1L], LOC105873340; [IGLL1], LOC109729893; [IGLL5], LOC105882024; [RPS3], LOC105862350; [RPS20], LOC105874908; [RPS27L], LOC109731171; [RPL32], LOC105861123; [RPLP1], LOC105859117; [RPS15A], LOC105857549; [RPL29], LOC105863618; [FTL], LOC105870251).

**f**. Dot plot of expression of adipokines *LEP* and *ADIPOQ* as well as their receptors across the atlas cell types (L1-L4, 10x dataset). Note abundant and specific expression of *ADIPOQ* but lack of *LEP* expression in adipocytes. Curiously, *LEP* transcripts were detected in the *AKR1B1*+ kidney loop of Henle cells, mesothelial cells, and some vascular-associated smooth muscle cells, although at very low levels. In contrast, *LEPR* shows expected expression in various cell types including tendon cells, fibroblasts, and endothelial cells, and high *LEPR* expression is found in mesenchymal progenitor cell types such as osteo-CAR and adipo-CAR cells. Also note ubiquitous expression of *ADIPOR1* across almost all atlas cell types, and enriched expression of *ADIPOR2* in sperm lineage cells and adipocytes^42^.

**Figure S12. Expression patterns of PS genes across cell types and species**

**a-b**. Heat maps showing expression of the 425 PS genes across all lemur cell types in the atlas (10x dataset). Cell types are ordered in panel a by compartment, then cell type designation number, then tissue source, and in panel b by tissue source, then compartment, then cell type designation number. Genes are ordered (left to right) by compartment-enriched (or depleted) genes, broadly-expressed genes, and genes not detected in the atlas.

**c-h**. Dot plots showing average expression of compartment-enriched (or depleted) PS genes across lemur cell types in the atlas (10x dataset). Genes are organized by enriched compartments (epithelial (c), endothelial (d), stromal (e), immune (f), neural (g), germ (h)). Genes are shown in more than one panel if enriched in multiple compartments. See Table 8 for a full list of the PS genes and their description.

**i**. Dot plot showing cross-species expression of all PS genes, as in Fig. 7f. Rows are orthologous genes, indicated with the respective human gene symbols and followed by the respective lemur gene symbol if different. Columns are cell types, displayed as doublets of the respective expression in human (H) and lemur (L). Cell types are ordered first by compartments, then by tissue, and finally by species.

**Figure S13. Additional characterization of the nonsense mutation and the affected genes**

**a**. The length of preserved and C-terminus depleted portion of each mutant protein, predicted based on the position of the nonsense mutation in the gene. Numbers next to the bar indicate the total length of the protein (black) and the percent of the depleted portion (red).

**b**. Percent of transcript reads that cover the mutation position over all transcripts reads that align to the corresponding gene (10x datasets).

**c**. Dot plot showing cross-species expression of the four genes across the 63 orthologous cell types in human, lemur, and mouse (when gene present), as analyzed in the evolutionary comparison analysis of the accompanying Tabula Microcebus manuscript^6^. Rows are orthologous genes, indicated with the respective human gene symbols. ^*^ lemur symbol is LOC105864482. Columns are cell types, displayed as triplets of the respective expression in human (H), lemur (L), and mouse (M). Cell types are ordered first by compartments, then by tissue, and finally by species. Red cross indicates the gene missing in the mouse genome.

