## Supplementary figures for "Mouse lemur transcriptomic atlas informs primate genes, mutations, physiology, and disease"

**Fig. S1**

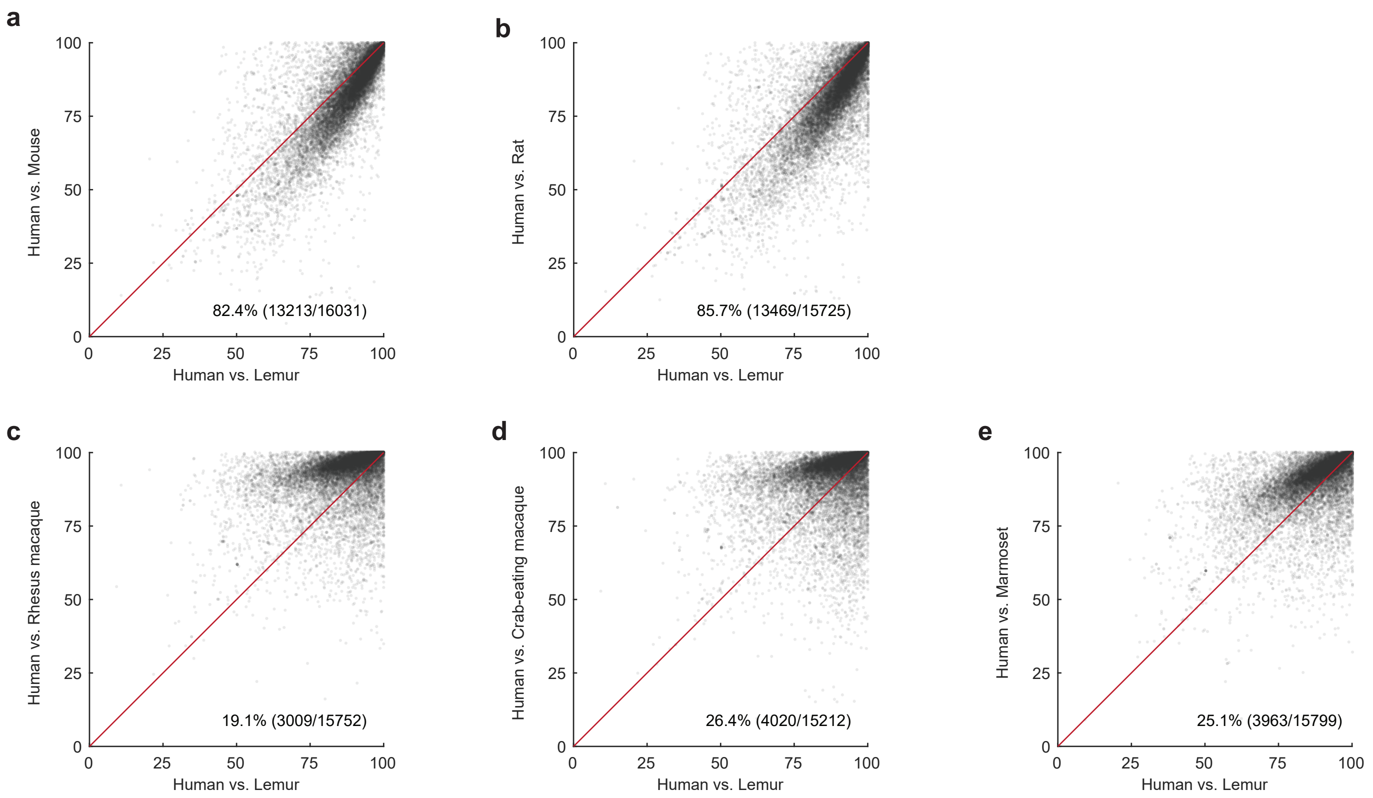
a
Human vs. Mouse
Human vs. Lemur
82.4% (13213/16031)
b
Human vs. Rat
Human vs. Lemur
85.7% (13469/15725)
c
Human vs. Rhesus macaque
Human vs. Lemur
19.1% (3009/15752)
d
Human vs. Crab-eating macaque
Human vs. Lemur
26.4% (4020/15212)
e
Human vs. Marmoset
Human vs. Lemur
25.1% (3963/15799)
0
25
50
75
100

Fig. S2

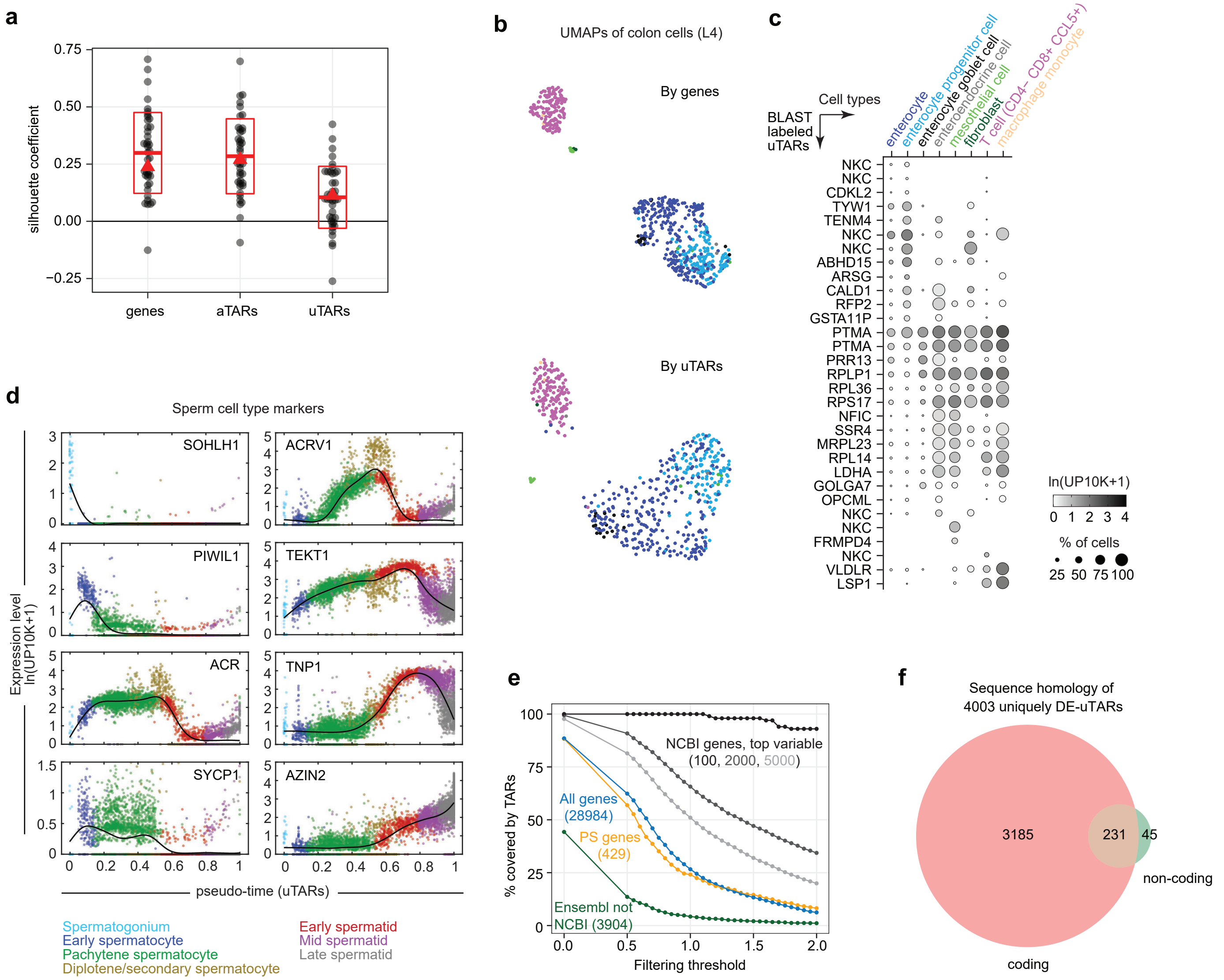
a
silhouette coefficient
0.75
0.50
0.25
0.00
−0.25
genes
aTARs
uTARs
b
UMAPs of colon cells (L4)
By genes
By uTARs
c
Cell types
BLAST labeled uTARs
enterocyte
enterocyte progenitor cell
enterocyte goblet cell
enteroendocrine cell
mesothelial cell
fibroblast
T cell (CD4− CD8+ CCL5+)
macrophage monocyte
NKC
NKC
CDKL2
TYW1
TENM4
NKC
NKC
ABHD15
ARSG
CALD1
RFP2
GSTA11P
PTMA
PTMA
PRR13
RPLP1
RPL36
RPS17
NFIC
SSR4
MRPL23
RPL14
LDHA
GOLGA7
OPCML
NKC
NKC
FRMPD4
NKC
VLDLR
LSP1
ln(UP10K+1)
0 1 2 3 4
% of cells
25 50 75 100
d
Sperm cell type markers
Expression level ln(UP10K+1)
SOHLH1
ACRV1
PIWIL1
TEKT1
ACR
TNP1
SYCP1
AZIN2
pseudo-time (uTARs)
Spermatogonium
Early spermatocyte
Pachytene spermatocyte
Diplotene/secondary spermatocyte
Early spermatid
Mid spermatid
Late spermatid
e
% covered by TARs
NCBI genes, top variable (100, 2000, 5000)
All genes (28984)
PS genes (429)
Ensembl not NCBI (3904)
Filtering threshold
f
Sequence homology of 4003 uniquely DE-uTARs
3185
231
45
non-coding
coding

Fig. S3

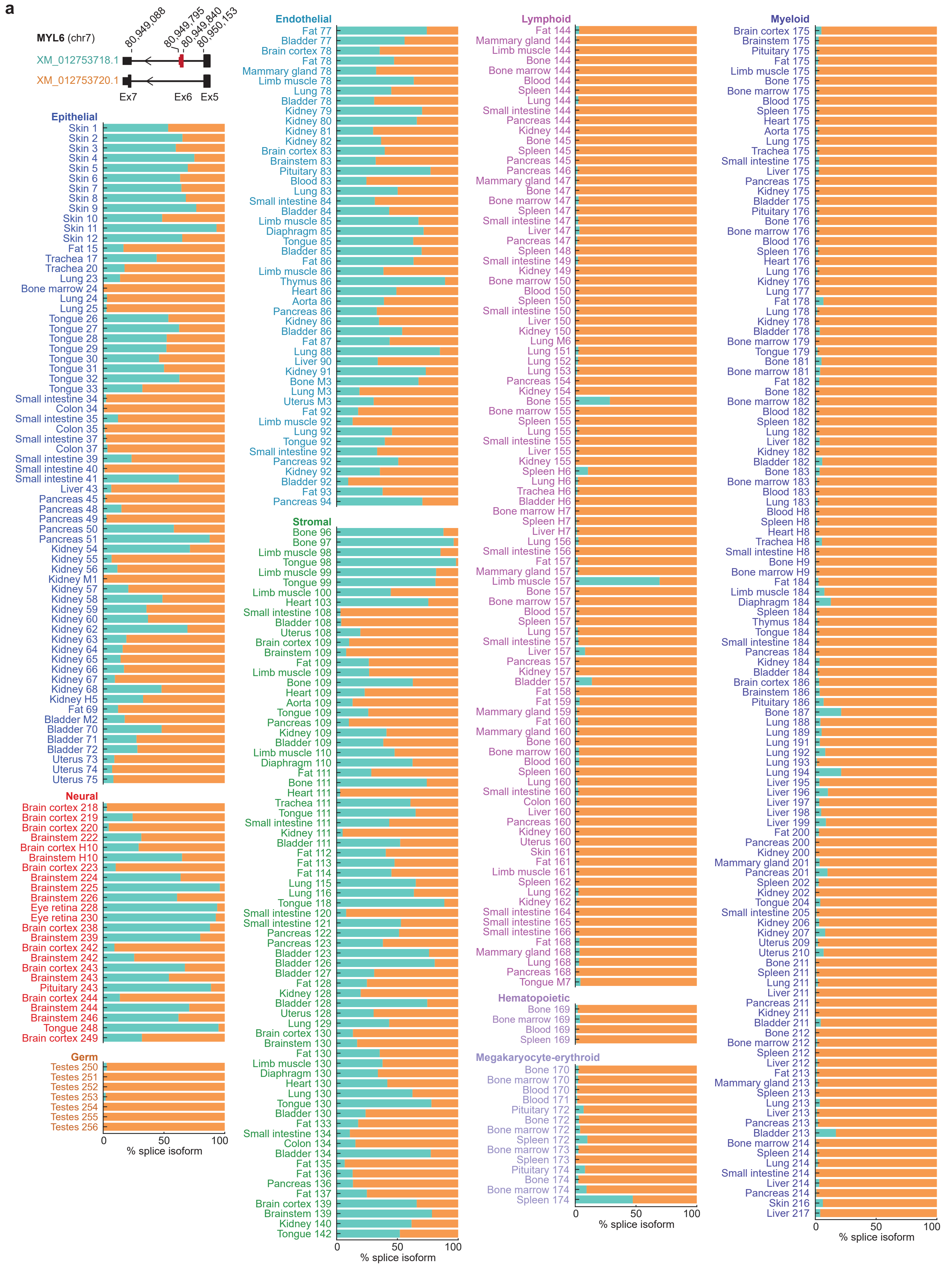
a
MYL6 (chr7)
80,949,088
80,949,795
80,949,840
80,950,153
XM_012753718.1
XM_012753720.1
Ex7
Ex6
Ex5
Epithelial
Skin 1
Skin 2
Skin 3
Skin 4
Skin 5
Skin 6
Skin 7
Skin 8
Skin 9
Skin 10
Skin 11
Skin 12
Fat 15
Trachea 17
Trachea 20
Lung 23
Bone marrow 24
Lung 24
Lung 25
Tongue 26
Tongue 27
Tongue 28
Tongue 29
Tongue 30
Tongue 31
Tongue 32
Tongue 33
Small intestine 34
Colon 34
Small intestine 35
Colon 35
Small intestine 37
Colon 37
Small intestine 39
Small intestine 40
Small intestine 41
Liver 43
Pancreas 45
Pancreas 48
Pancreas 49
Pancreas 50
Pancreas 51
Kidney 54
Kidney 55
Kidney 56
Kidney M1
Kidney 57
Kidney 58
Kidney 59
Kidney 60
Kidney 62
Kidney 63
Kidney 64
Kidney 65
Kidney 66
Kidney 67
Kidney 68
Kidney H5
Fat 69
Bladder M2
Bladder 70
Bladder 71
Bladder 72
Uterus 73
Uterus 74
Uterus 75
Neural
Brain cortex 218
Brain cortex 219
Brain cortex 220
Brainstem 222
Brain cortex H10
Brainstem H10
Brain cortex 223
Brainstem 224
Brainstem 225
Brainstem 226
Eye retina 228
Eye retina 230
Brain cortex 238
Brainstem 239
Brain cortex 242
Brainstem 242
Brain cortex 243
Brainstem 243
Pituitary 243
Brain cortex 244
Brainstem 244
Brainstem 246
Tongue 248
Brain cortex 249
Germ
Testes 250
Testes 251
Testes 252
Testes 253
Testes 254
Testes 255
Testes 256
0
50
100
% splice isoform
Endothelial
Fat 77
Bladder 77
Brain cortex 78
Fat 78
Mammary gland 78
Limb muscle 78
Lung 78
Bladder 78
Kidney 79
Kidney 80
Kidney 81
Kidney 82
Brain cortex 83
Brainstem 83
Pituitary 83
Blood 83
Lung 83
Small intestine 84
Bladder 84
Limb muscle 85
Diaphragm 85
Tongue 85
Bladder 85
Fat 86
Limb muscle 86
Thymus 86
Heart 86
Aorta 86
Pancreas 86
Kidney 86
Bladder 86
Fat 87
Lung 88
Liver 90
Kidney 91
Bone M3
Lung M3
Uterus M3
Fat 92
Limb muscle 92
Lung 92
Tongue 92
Small intestine 92
Pancreas 92
Kidney 92
Bladder 92
Fat 93
Pancreas 94
Stromal
Bone 96
Bone 97
Limb muscle 98
Tongue 98
Limb muscle 99
Tongue 99
Limb muscle 100
Heart 103
Small intestine 108
Bladder 108
Uterus 108
Brain cortex 109
Brainstem 109
Fat 109
Limb muscle 109
Bone 109
Heart 109
Aorta 109
Tongue 109
Pancreas 109
Kidney 109
Bladder 109
Limb muscle 110
Diaphragm 110
Fat 111
Bone 111
Heart 111
Trachea 111
Tongue 111
Small intestine 111
Kidney 111
Bladder 111
Fat 112
Fat 113
Fat 114
Lung 115
Lung 116
Tongue 118
Small intestine 120
Small intestine 121
Pancreas 122
Pancreas 123
Bladder 123
Bladder 126
Bladder 127
Fat 128
Kidney 128
Bladder 128
Uterus 128
Lung 129
Brain cortex 130
Brainstem 130
Fat 130
Limb muscle 130
Diaphragm 130
Heart 130
Lung 130
Tongue 130
Bladder 130
Fat 133
Small intestine 134
Colon 134
Bladder 134
Fat 135
Fat 136
Pancreas 136
Fat 137
Brain cortex 139
Brainstem 139
Kidney 140
Tongue 142
0
50
100
% splice isoform
Lymphoid
Fat 144
Mammary gland 144
Limb muscle 144
Tongue 144
Bone marrow 144
Blood 144
Spleen 144
Lung 144
Small intestine 144
Pancreas 144
Kidney 144
Bone 145
Spleen 145
Pancreas 145
Pancreas 146
Mammary gland 147
Bone 147
Bone marrow 147
Spleen 147
Small intestine 147
Liver 147
Pancreas 147
Spleen 148
Small intestine 149
Kidney 149
Bone marrow 150
Blood 150
Spleen 150
Small intestine 150
Liver 150
Kidney 150
Lung M6
Lung 151
Lung 152
Lung 153
Pancreas 154
Kidney 154
Bone 155
Bone marrow 155
Spleen 155
Lung 155
Small intestine 155
Liver 155
Kidney 155
Spleen H6
Lung H6
Trachea H6
Bladder H6
Bone marrow H7
Spleen H7
Liver H7
Lung 156
Small intestine 156
Fat 157
Mammary gland 157
Limb muscle 157
Bone 157
Bone marrow 157
Blood 157
Spleen 157
Lung 157
Small intestine 157
Liver 157
Pancreas 157
Kidney 157
Bladder 157
Fat 158
Fat 159
Mammary gland 159
Fat 160
Mammary gland 160
Bone 160
Bone marrow 160
Blood 160
Spleen 160
Lung 160
Small intestine 160
Colon 160
Liver 160
Pancreas 160
Kidney 160
Uterus 160
Skin 161
Fat 161
Limb muscle 161
Spleen 162
Lung 162
Kidney 162
Small intestine 164
Small intestine 165
Small intestine 166
Fat 168
Mammary gland 168
Lung 168
Pancreas 168
Tongue M7
Hematopoietic
Bone 169
Bone marrow 169
Blood 169
Spleen 169
Megakaryocyte-erythroid
Bone 170
Bone marrow 170
Blood 170
Blood 171
Pituitary 172
Bone 172
Bone marrow 172
Spleen 172
Bone marrow 173
Spleen 173
Pituitary 174
Bone 174
Bone marrow 174
Spleen 174
0
50
100
% splice isoform
Myeloid
Brain cortex 175
Brainstem 175
Pituitary 175
Fat 175
Limb muscle 175
Bone 175
Bone marrow 175
Blood 175
Spleen 175
Heart 175
Aorta 175
Lung 175
Trachea 175
Small intestine 175
Liver 175
Pancreas 175
Kidney 175
Bladder 175
Pituitary 176
Bone 176
Bone marrow 176
Blood 176
Spleen 176
Heart 176
Lung 176
Kidney 176
Lung 177
Fat 178
Lung 178
Kidney 178
Bladder 178
Bone marrow 179
Tongue 179
Bone 181
Bone marrow 181
Fat 182
Bone 182
Bone marrow 182
Blood 182
Spleen 182
Lung 182
Liver 182
Kidney 182
Bladder 182
Bone 183
Bone marrow 183
Blood 183
Lung 183
Blood H8
Spleen H8
Heart H8
Trachea H8
Small intestine H8
Bone H9
Bone marrow H9
Fat 184
Limb muscle 184
Diaphragm 184
Spleen 184
Thymus 184
Tongue 184
Small intestine 184
Pancreas 184
Kidney 184
Bladder 184
Brain cortex 186
Brainstem 186
Pituitary 186
Bone 187
Lung 188
Lung 189
Lung 191
Lung 192
Lung 193
Lung 194
Liver 195
Liver 196
Liver 197
Liver 198
Liver 199
Fat 200
Pancreas 200
Kidney 200
Mammary gland 201
Pancreas 201
Spleen 202
Kidney 202
Tongue 204
Small intestine 205
Kidney 206
Kidney 207
Uterus 209
Uterus 210
Bone 211
Spleen 211
Lung 211
Liver 211
Pancreas 211
Kidney 211
Bladder 211
Bone 212
Bone marrow 212
Spleen 212
Liver 212
Fat 213
Mammary gland 213
Spleen 213
Lung 213
Liver 213
Pancreas 213
Bladder 213
Bone marrow 214
Spleen 214
Lung 214
Small intestine 214
Liver 214
Pancreas 214
Skin 216
Liver 217
0
50
100
% splice isoform

**Fig. S3 - continued**

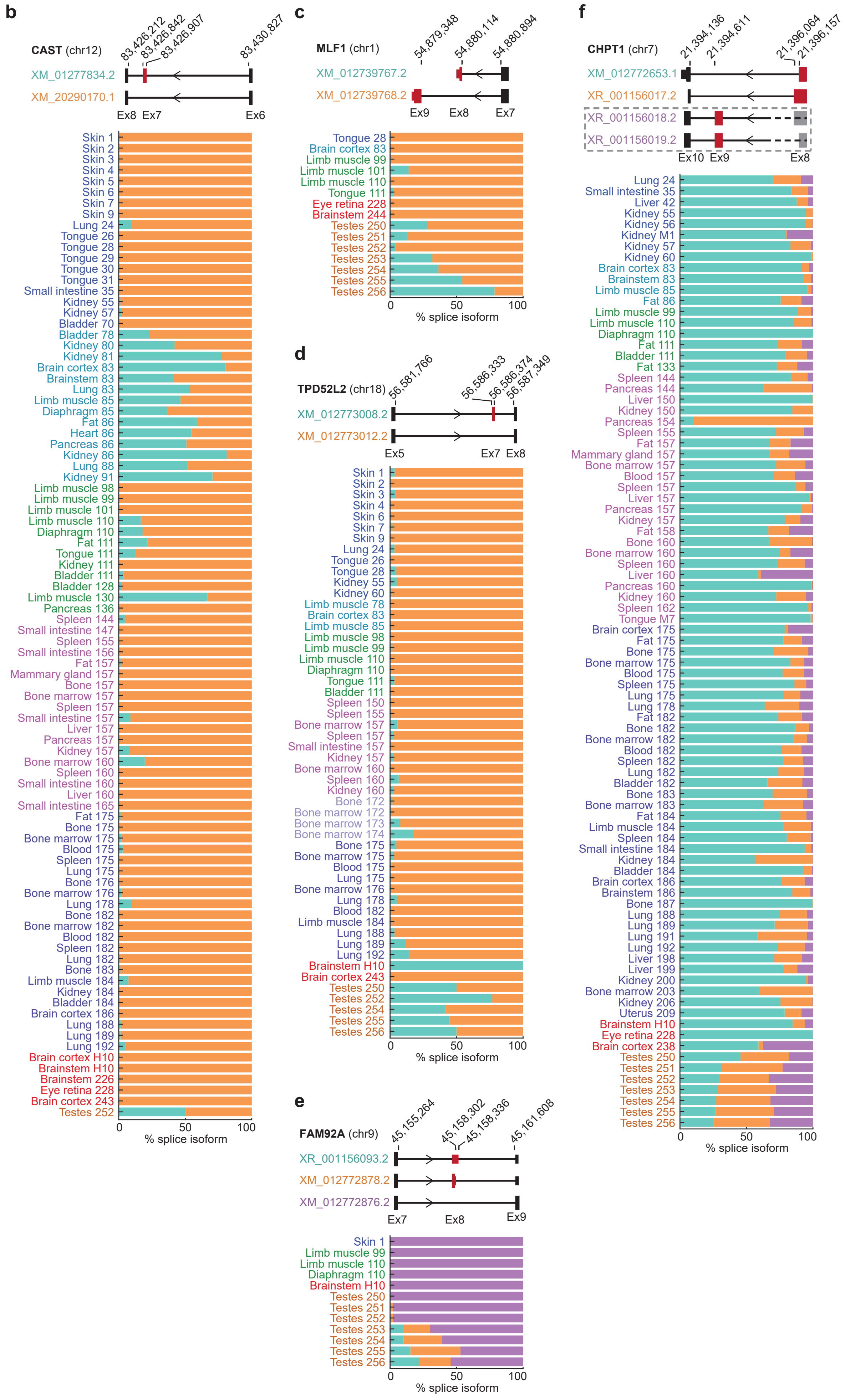

Fig. S4

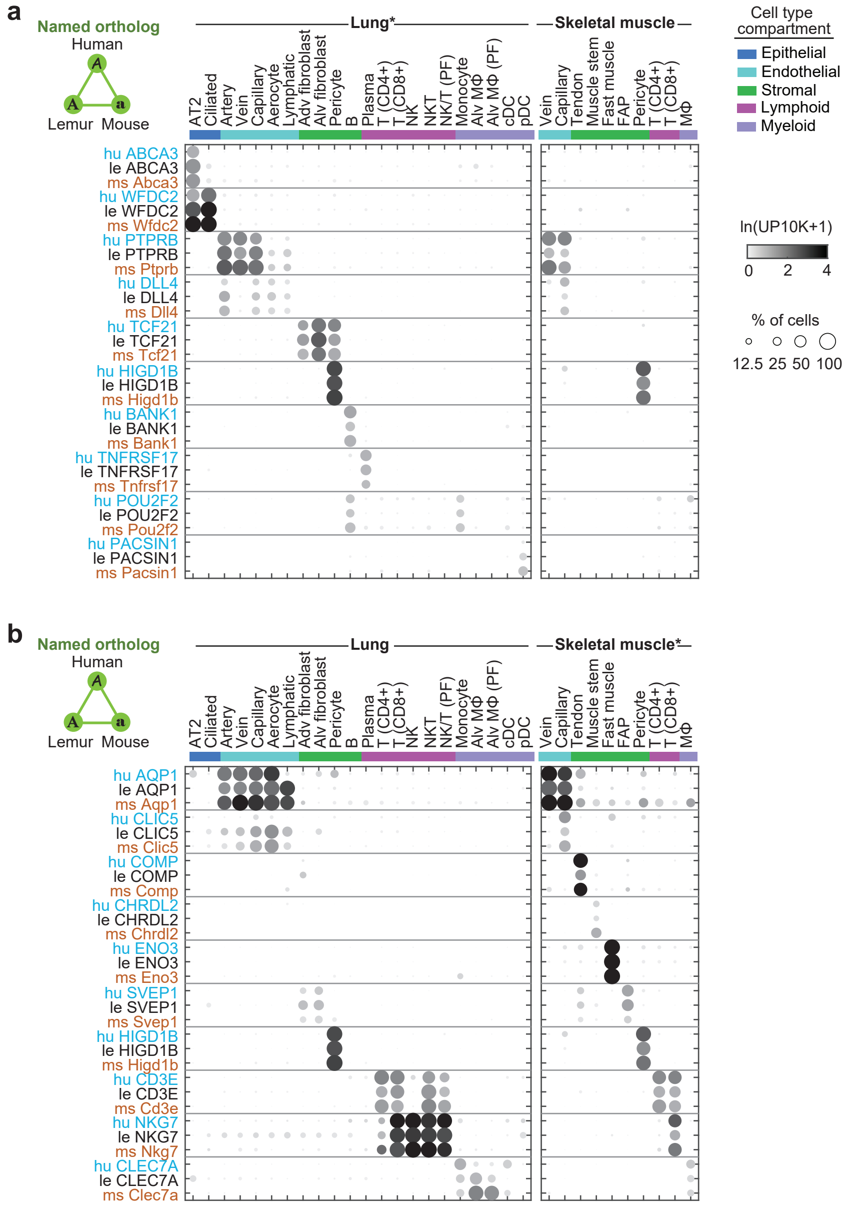

a
Named ortholog
Human
Lemur Mouse
Lung*
Skeletal muscle
AT2
Ciliated
Artery
Vein
Capillary
Aerocyte
Lymphatic
Adv fibroblast
Alv fibroblast
Pericyte
B
Plasma
T (CD4+)
T (CD8+)
NK
NKT
NK/T (PF)
Monocyte
Alv MΦ
Alv MΦ (PF)
cDC
pDC
Vein
Capillary
Tendon
Muscle stem
Fast muscle
FAP
Pericyte
T (CD4+)
T (CD8+)
MΦ
Cell type compartment
Epithelial
Endothelial
Stromal
Lymphoid
Myeloid
ln(UP10K+1)
0 2 4
% of cells
12.5 25 50 100
hu ABCA3
le ABCA3
ms Abca3
hu WFDC2
le WFDC2
ms Wfdc2
hu PTPRB
le PTPRB
ms Ptprb
hu DLL4
le DLL4
ms Dll4
hu TCF21
le TCF21
ms Tcf21
hu HIGD1B
le HIGD1B
ms Higd1b
hu BANK1
le BANK1
ms Bank1
hu TNFRSF17
le TNFRSF17
ms Tnfrsf17
hu POU2F2
le POU2F2
ms Pou2f2
hu PACSIN1
le PACSIN1
ms Pacsin1
b
Named ortholog
Human
Lemur Mouse
Lung
Skeletal muscle*
hu AQP1
le AQP1
ms Aqp1
hu CLIC5
le CLIC5
ms Clic5
hu COMP
le COMP
ms Comp
hu CHRDL2
le CHRDL2
ms Chrdl2
hu ENO3
le ENO3
ms Eno3
hu SVEP1
le SVEP1
ms Svep1
hu HIGD1B
le HIGD1B
ms Higd1b
hu CD3E
le CD3E
ms Cd3e
hu NKG7
le NKG7
ms Nkg7
hu CLEC7A
le CLEC7A
ms Clec7a

**Fig. S5**

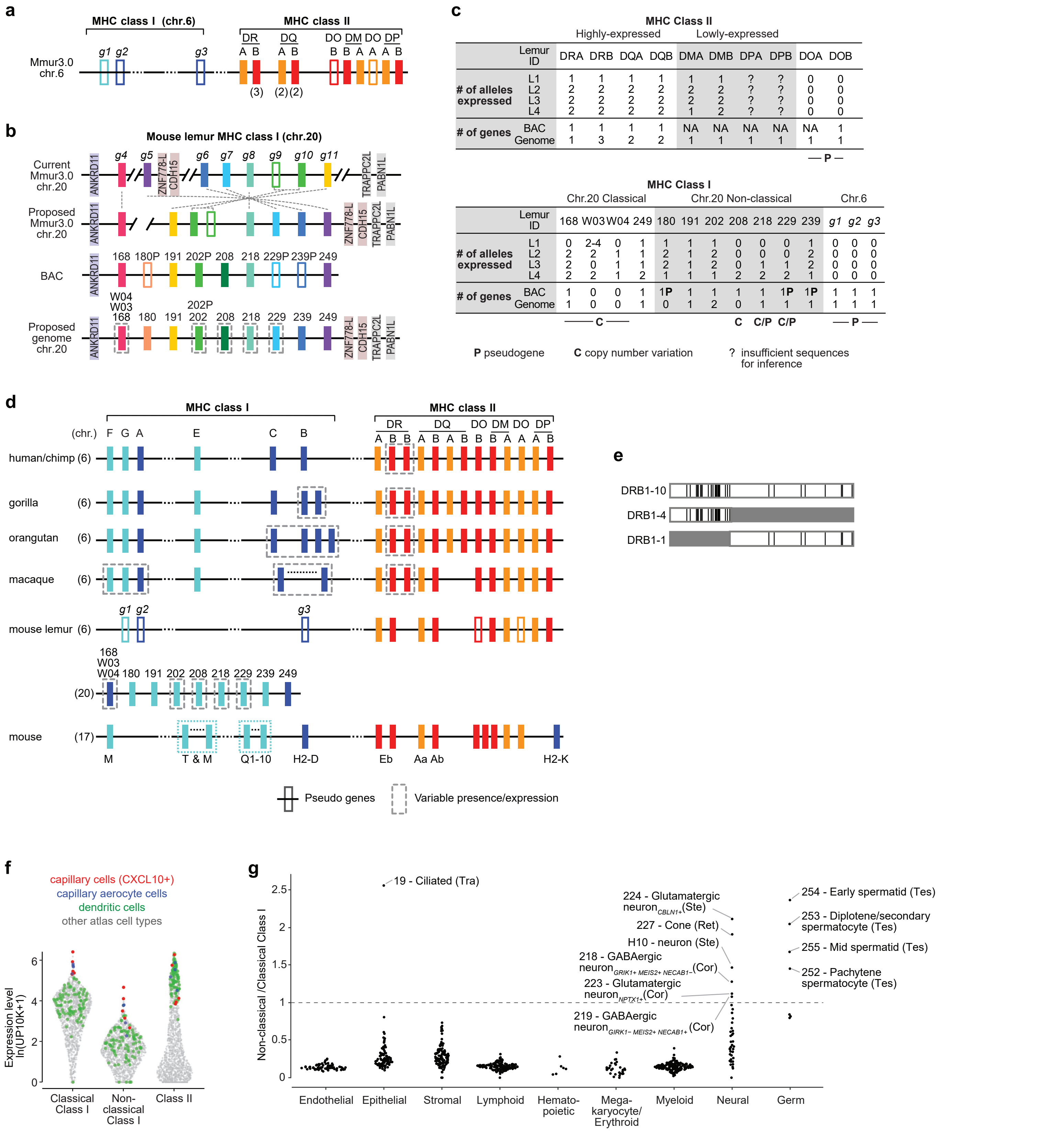
a
MHC class I (chr.6)
MHC class II
DR DQ DO DM DO DP
A B A B B B A A A B
Mmur3.0 chr.6
g1 g2 g3
(3) (2) (2)
b
Mouse lemur MHC class I (chr.20)
Current Mmur3.0 chr.20
g4 g5 g6 g7 g8 g9 g10 g11
ANKRD11 ZNF778-L CDH15 TRAPPC2L PABN1L
Proposed Mmur3.0 chr.20
BAC
168 180P 191 202P 208 218 229P 239P 249
Proposed genome chr.20
W04 W03 168 180 191 202P 202 208 218 229 239 249
c
MHC Class II
Highly-expressed
Lowly-expressed
Lemur ID DRA DRB DQA DQB DMA DMB DPA DPB DOA DOB
### of alleles expressed L1 1 1 1 1 1 1 ? ? 0 0
L2 2 2 2 2 2 2 ? ? 0 0
L3 2 2 2 2 2 2 ? ? 0 0
L4 2 2 2 2 1 2 ? ? 0 0
### of genes BAC 1 1 1 1 NA NA NA NA NA 1
Genome 1 3 2 2 1 1 1 1 1 1
P
MHC Class I
Chr.20 Classical
Chr.20 Non-classical
Chr.6
Lemur ID 168 W03 W04 249 180 191 202 208 218 229 239 g1 g2 g3
### of alleles expressed L1 0 2-4 0 1 1 1 1 0 0 0 1 0 0 0
L2 2 2 1 1 2 1 2 0 0 0 2 0 0 0
L3 2 0 1 1 2 1 2 0 1 1 2 0 0 0
L4 2 2 1 2 1 1 1 2 2 2 1 0 0 0
### of genes BAC 1 0 0 1 1P 1 1 1 1 1P 1P 1 1 1
Genome 1 0 0 1 0 1 2 0 1 1 1 1 1 1
C
C C/P C/P
P
P pseudogene
C copy number variation
? insufficient sequences for inference
d
MHC class I
MHC class II
DR DQ DO DM DO DP
(chr.) F G A E C B
A B B A B A B B B A A A B
human/chimp (6)
gorilla (6)
orangutan (6)
macaque (6)
mouse lemur (6)
g1 g2 g3
168 W03 W04 180 191 202 208 218 229 239 249
(20)
mouse (17)
M T & M Q1-10 H2-D Eb Aa Ab H2-K
Pseudo genes
Variable presence/expression
e
DRB1-10
DRB1-4
DRB1-1
f
capillary cells (CXCL10+)
capillary aerocyte cells
dendritic cells
other atlas cell types
Expression level ln(UP10K+1)
0 2 4 6
Classical Class I
Non-classical Class I
Class II
g
Non-classical /Classical Class I
0 0.5 1 1.5 2 2.5
19 - Ciliated (Tra)
224 - Glutamatergic neuron CBLN1+ (Ste)
227 - Cone (Ret)
H10 - neuron (Ste)
218 - GABAergic neuron GRIK1+ MEIS2+ NECAB1- (Cor)
223 - Glutamatergic neuron NPTX1+ (Cor)
219 - GABAergic neuron GRIK1- MEIS2+ NECAB1+ (Cor)
254 - Early spermatid (Tes)
253 - Diplotene/secondary spermatocyte (Tes)
255 - Mid spermatid (Tes)
252 - Pachytene spermatocyte (Tes)
Endothelial
Epithelial
Stromal
Lymphoid
Hemato-poietic
Mega-karyocyte/ Erythroid
Myeloid
Neural
Germ

#### Fig. S5 - continued

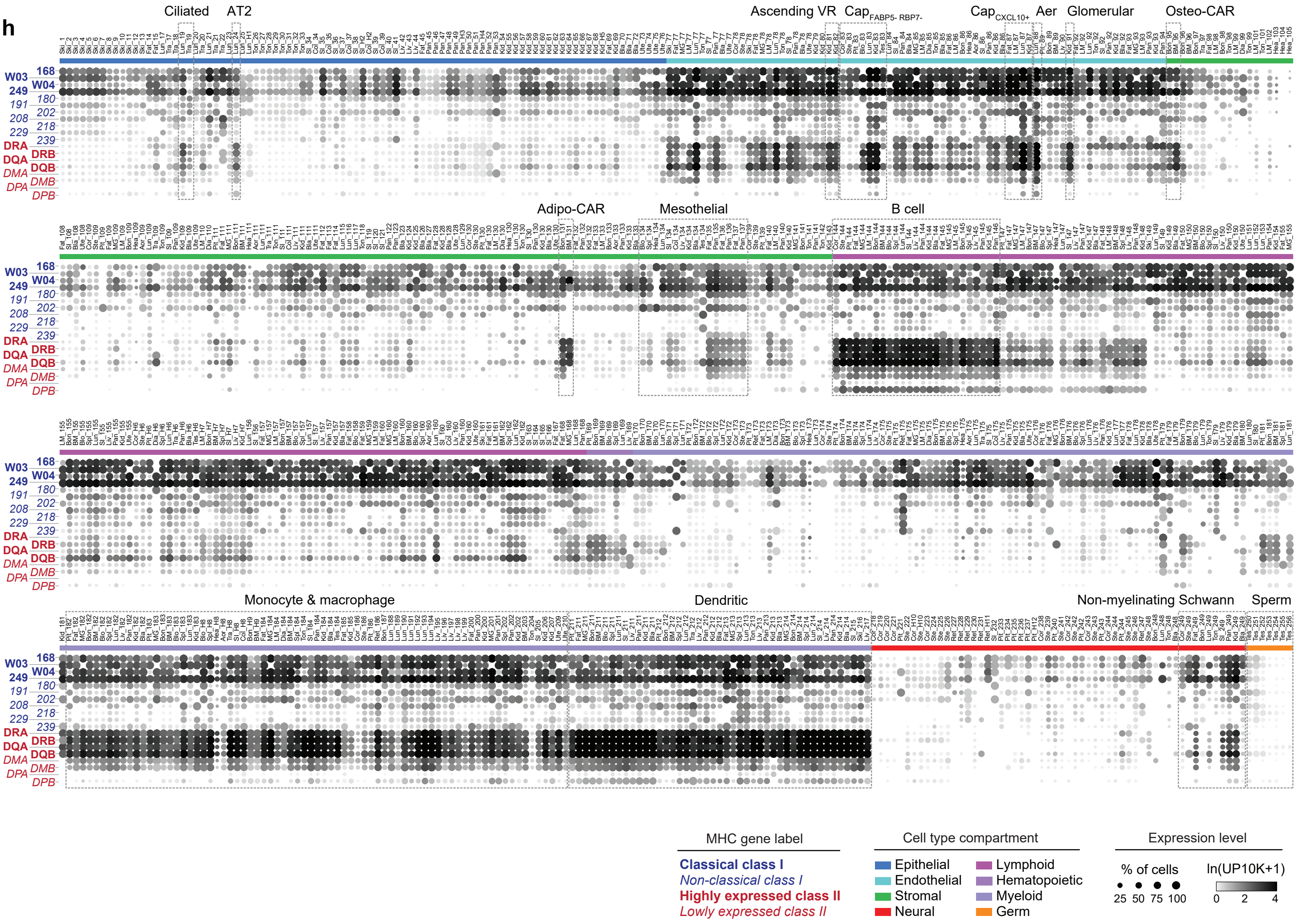

### Fig. S6

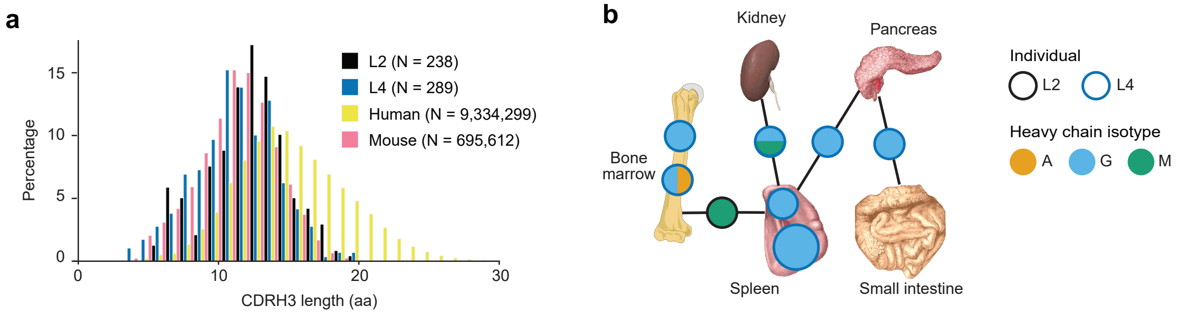
a
Percentage
CDRH3 length (aa)
0
5
10
15
20
30
L2 (N = 238)
L4 (N = 289)
Human (N = 9,334,299)
Mouse (N = 695,612)
b
Kidney
Pancreas
Bone marrow
Spleen
Small intestine
Individual
L2
L4
Heavy chain isotype
A
G
M

**Fig. S7**

**a**

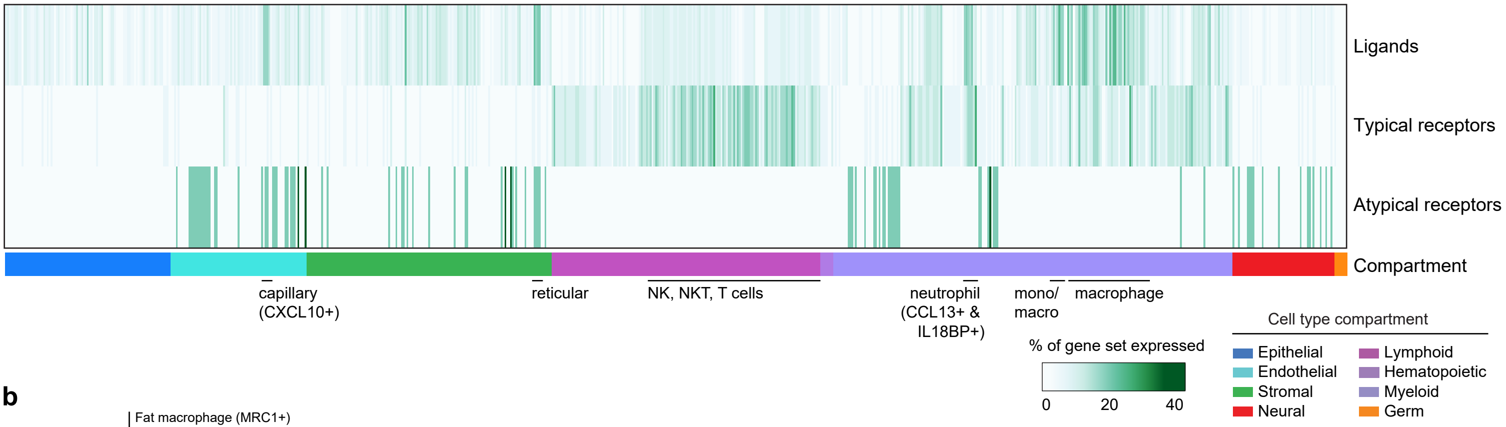
Ligands
Typical receptors
Atypical receptors
Compartment
capillary (CXCL10+)
reticular
NK, NKT, T cells
neutrophil (CCL13+ & IL18BP+)
mono/ macro
macrophage
% of gene set expressed
0
20
40
Cell type compartment
Epithelial
Endothelial
Stromal
Neural
Lymphoid
Hematopoietic
Myeloid
Germ

**b**

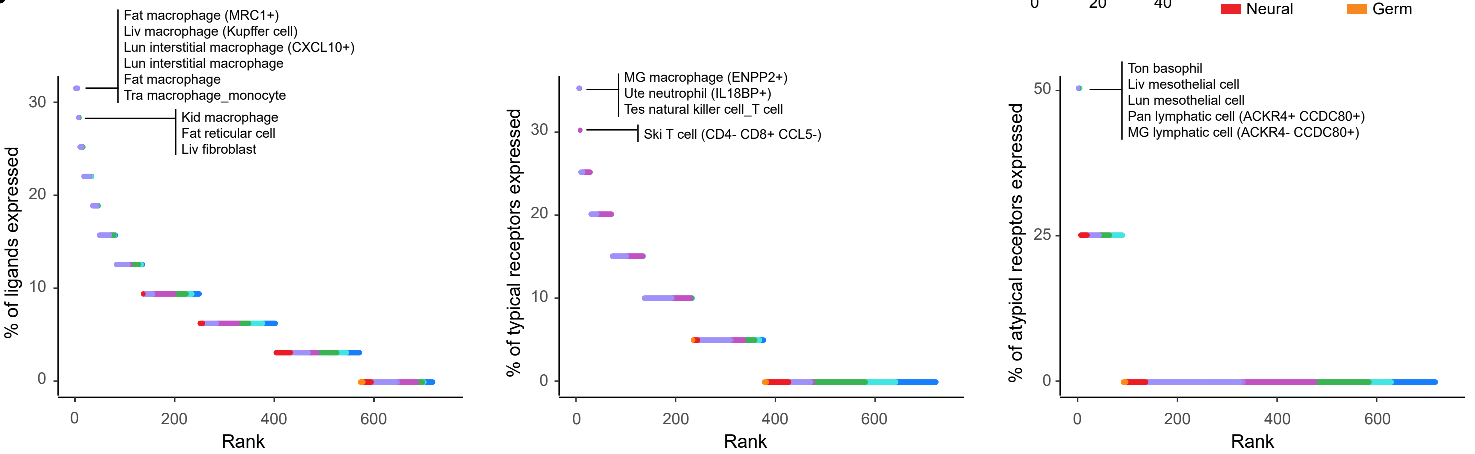
Fat macrophage (MRC1+)
Liv macrophage (Kupffer cell)
Lun interstitial macrophage (CXCL10+)
Lun interstitial macrophage
Fat macrophage
Tra macrophage_monocyte
Kid macrophage
Fat reticular cell
Liv fibroblast
% of ligands expressed
Rank
MG macrophage (ENPP2+)
Ute neutrophil (IL18BP+)
Tes natural killer cell_T cell
Ski T cell (CD4- CD8+ CCL5-)
% of typical receptors expressed
Rank
Ton basophil
Liv mesothelial cell
Lun mesothelial cell
Pan lymphatic cell (ACKR4+ CCDC80+)
MG lymphatic cell (ACKR4- CCDC80+)
% of atypical receptors expressed
Rank

**c**

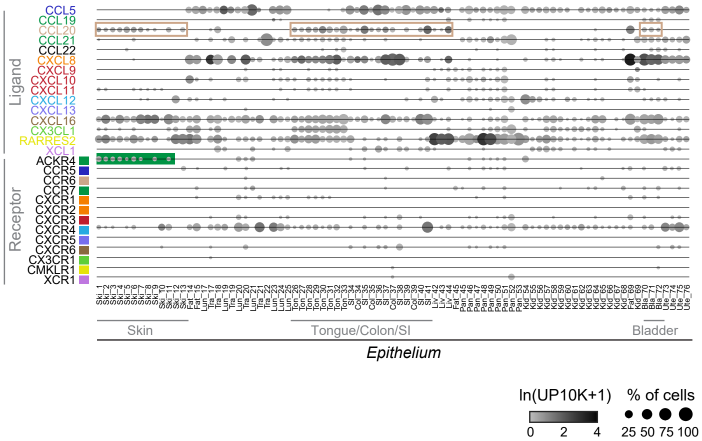
Ligand
CCL5
CCL19
CCL20
CCL21
CCL22
CXCL8
CXCL9
CXCL10
CXCL11
CXCL12
CXCL13
CXCL16
CX3CL1
RARRES2
XCL1
Receptor
ACKR4
CCR5
CCR6
CCR7
CXCR1
CXCR2
CXCR3
CXCR4
CXCR5
CXCR6
CX3CR1
CMKLR1
XCR1
Skin
Tongue/Colon/SI
Bladder
Epithelium
ln(UP10K+1)
0
2
4
% of cells
25 50 75 100

**Fig. S7 - continued**

**d**

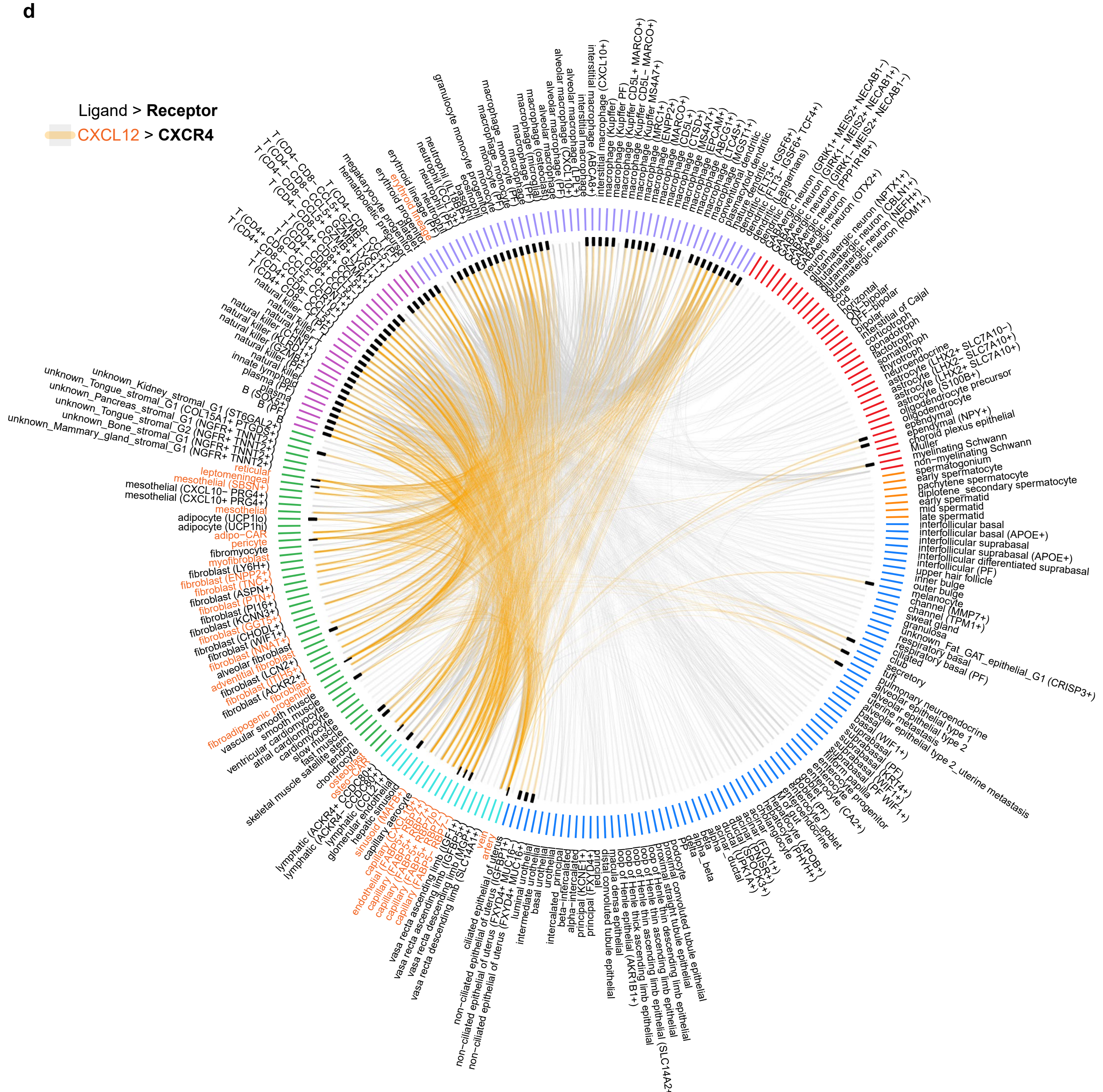

**Fig. S7 - continued**

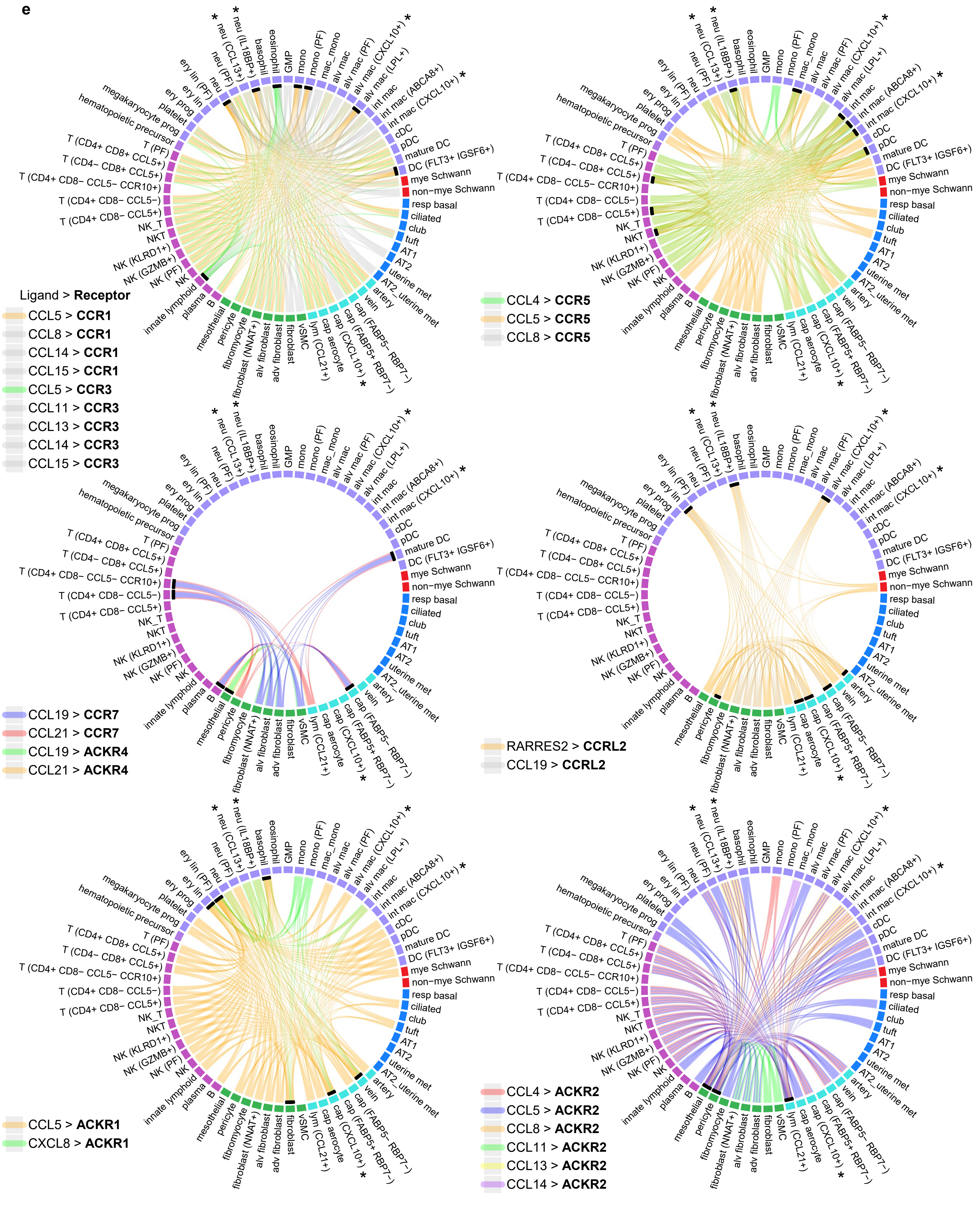

**Fig. S7 - continued**

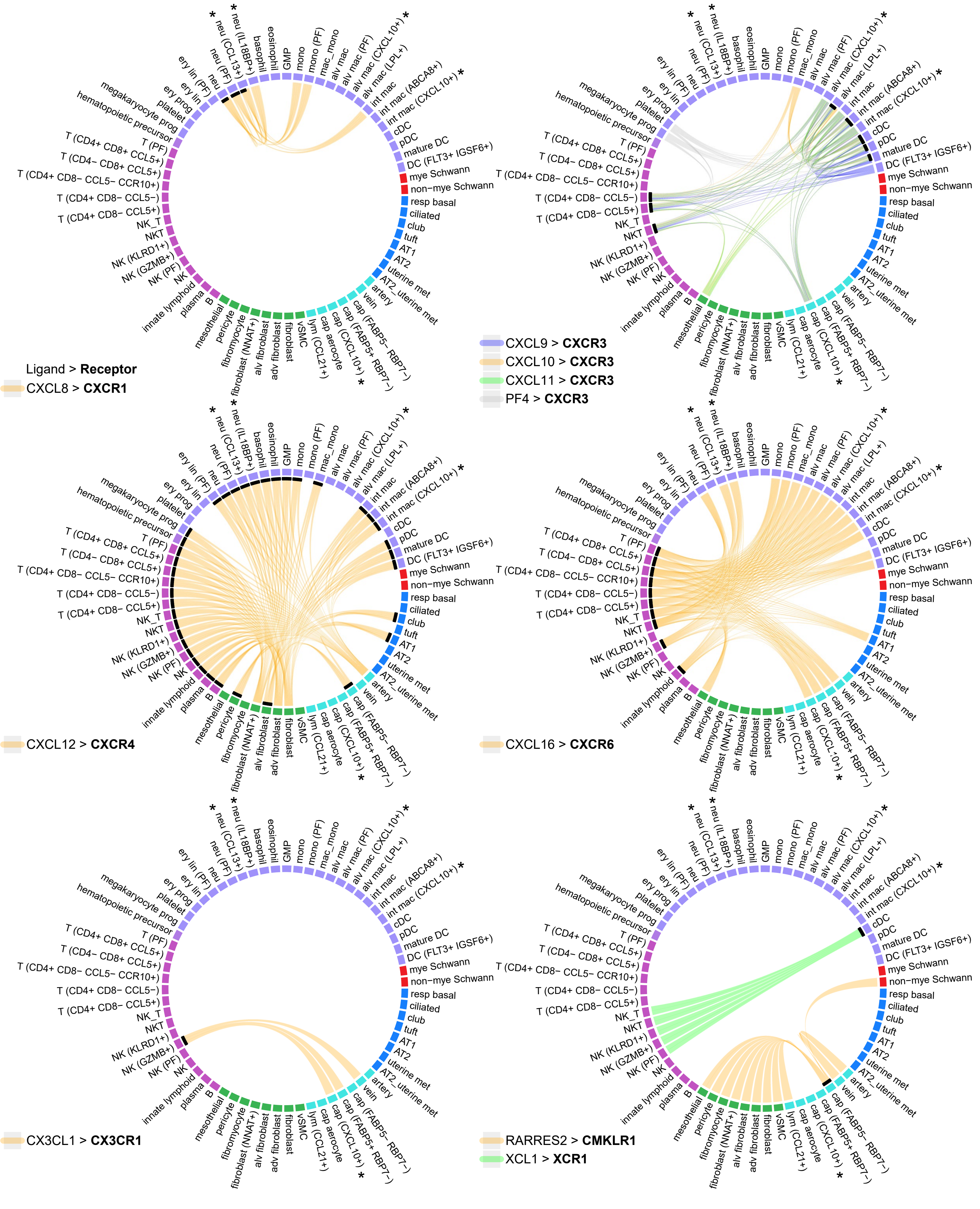

f

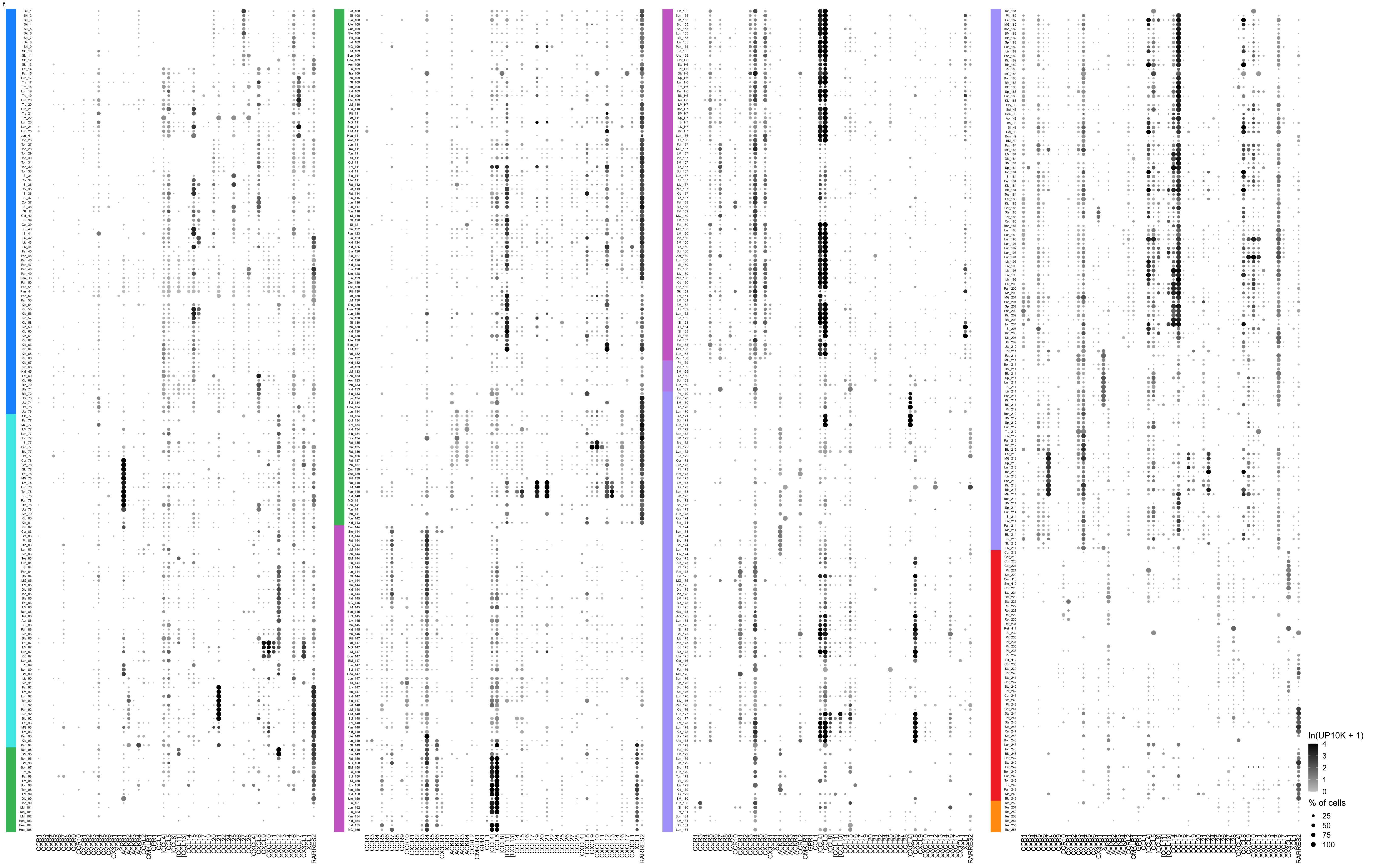

### Fig. S8

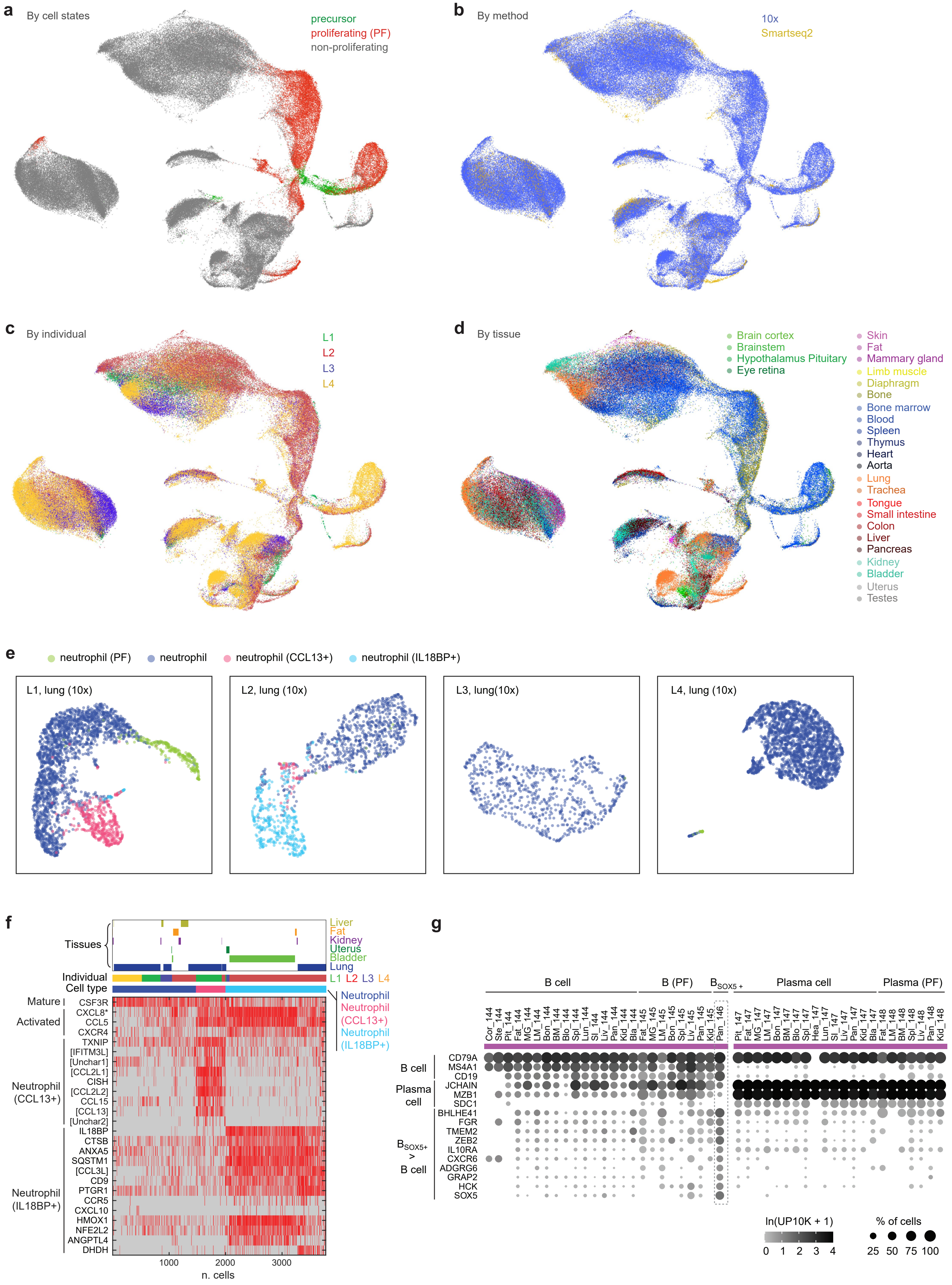

a
By cell states
precursor
proliferating (PF)
non-proliferating
b
By method
10x
Smartseq2
c
By individual
L1
L2
L3
L4
d
By tissue
Brain cortex
Brainstem
Hypothalamus Pituitary
Eye retina
Skin
Fat
Mammary gland
Limb muscle
Diaphragm
Bone
Bone marrow
Blood
Spleen
Thymus
Heart
Aorta
Lung
Trachea
Tongue
Small intestine
Colon
Liver
Pancreas
Kidney
Bladder
Uterus
Testes
e
neutrophil (PF)
neutrophil
neutrophil (CCL13+)
neutrophil (IL18BP+)
L1, lung (10x)
L2, lung (10x)
L3, lung(10x)
L4, lung (10x)
f
Tissues
Liver
Fat
Kidney
Uterus
Bladder
Lung
Individual
L1 L2 L3 L4
Cell type
Neutrophil
Neutrophil (CCL13+)
Neutrophil (IL18BP+)
Mature
CSF3R
Activated
CXCL8*
CCL5
CXCR4
TXNIP
[IFITM3L]
[Unchar1]
[CCL2L1]
Neutrophil (CCL13+)
CISH
[CCL2L2]
CCL15
[CCL13]
[Unchar2]
IL18BP
CTSB
ANXA5
SQSTM1
[CCL3L]
CD9
Neutrophil (IL18BP+)
PTGR1
CCR5
CXCL10
HMOX1
NFE2L2
ANGPTL4
DHDH
1000
2000
3000
n. cells
g
B cell
B (PF)
$B_{SOX5+}$
Plasma cell
Plasma (PF)
Cor_144
Ste_144
Pit_144
Fat_144
MG_144
LM_144
Bon_144
BM_144
Blo_144
Spl_144
Lun_144
SI_144
Liv_144
Pan_144
Kid_144
Bla_144
Fat_145
MG_145
LM_145
Bon_145
Spl_145
Liv_145
Pan_145
Kid_145
Pan_146
Pit_147
Fat_147
MG_147
LM_147
Bon_147
BM_147
Blo_147
Spl_147
Hea_147
Lun_147
SI_147
Liv_147
Pan_147
Kid_147
Bla_147
Fat_148
LM_148
BM_148
Spl_148
Liv_148
Pan_148
Kid_148
B cell
CD79A
MS4A1
CD19
Plasma cell
JCHAIN
MZB1
SDC1
$B_{SOX5+}$ > B cell
BHLHE41
FGR
TMEM2
ZEB2
IL10RA
CXCR6
ADGRG6
GRAP2
HCK
SOX5
ln(UP10K + 1)
0 1 2 3 4
% of cells
25 50 75 100

Fig. S9

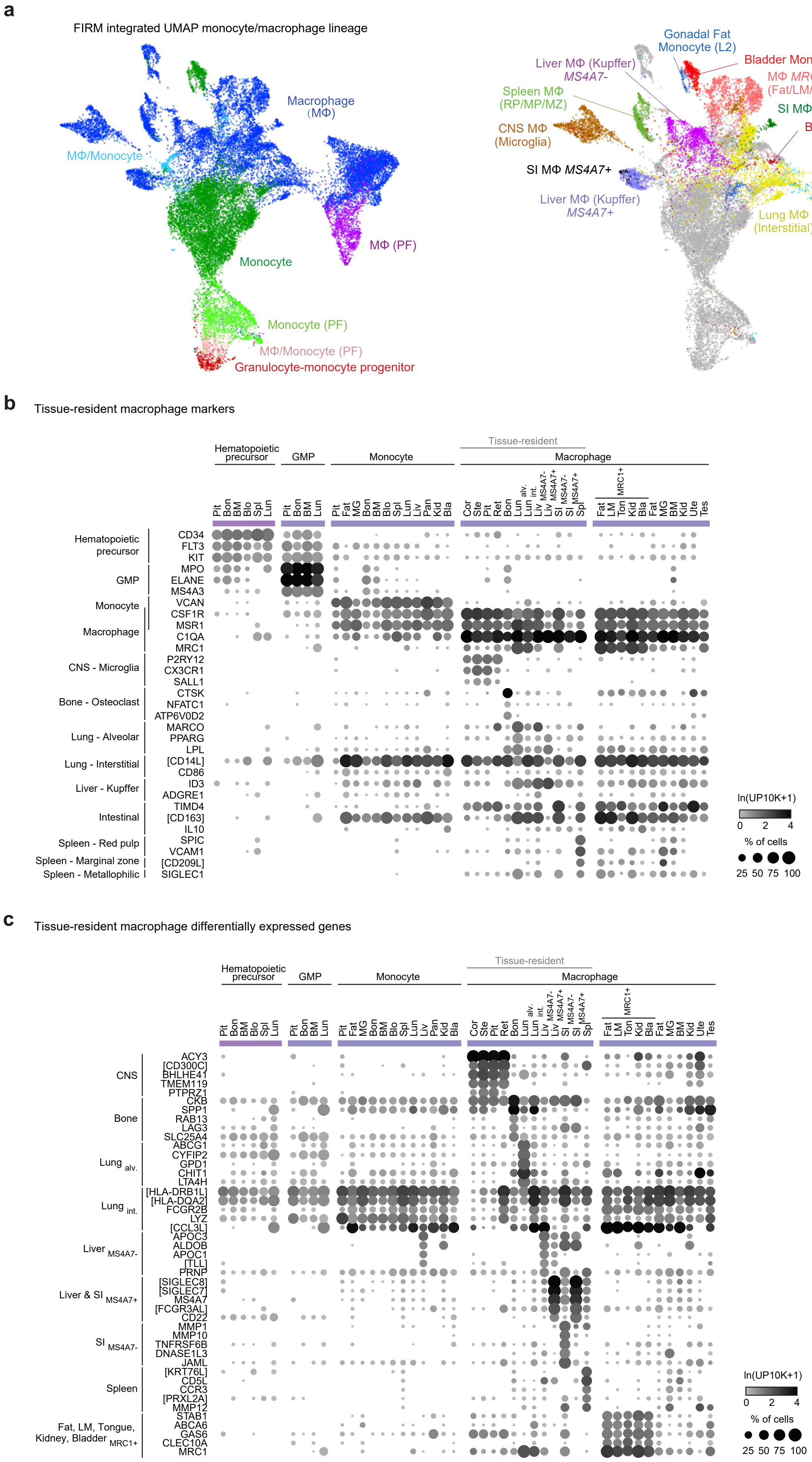
a
FIRM integrated UMAP monocyte/macrophage lineage
Macrophage (MΦ)
MΦ/Monocyte
MΦ (PF)
Monocyte
Monocyte (PF)
MΦ/Monocyte (PF)
Granulocyte-monocyte progenitor
Gonadal Fat Monocyte (L2)
Bladder Monocyte (L2)
Liver MΦ (Kupffer) MS4A7-
MΦ MRC1+ (Fat/LM/Tongue/Bladder/Kidney)
Spleen MΦ (RP/MP/MZ)
SI MΦ MS4A7-
CNS MΦ (Microglia)
Bone MΦ (Osteoclast)
SI MΦ MS4A7+
Liver MΦ (Kupffer) MS4A7+
Lung MΦ (Interstitial)
Lung MΦ (Alveolar)
b
Tissue-resident macrophage markers
Hematopoietic precursor
GMP
Monocyte
Tissue-resident
Macrophage
Pit Bon BM Blo Spl Lun
Pit Bon BM Lun
Pit Fat MG Bon BM Blo Spl Lun Liv Pan Kid Bla
Cor Ste Pit Ret Bon Lun alv. Lun int. Liv MS4A7- Liv MS4A7+ SI MS4A7- SI MS4A7+ Spl
MRC1+
Fat LM Ton Kid Bla Fat MG BM Kid Ute Tes
Hematopoietic precursor
CD34
FLT3
KIT
GMP
MPO
ELANE
MS4A3
Monocyte
VCAN
CSF1R
Macrophage
MSR1
C1QA
MRC1
CNS - Microglia
P2RY12
CX3CR1
SALL1
Bone - Osteoclast
CTSK
NFATC1
ATP6V0D2
Lung - Alveolar
MARCO
PPARG
LPL
Lung - Interstitial
[CD14L]
CD86
Liver - Kupffer
ID3
ADGRE1
Intestinal
TIMD4
[CD163]
IL10
Spleen - Red pulp
SPIC
VCAM1
Spleen - Marginal zone
[CD209L]
Spleen - Metallophilic
SIGLEC1
ln(UP10K+1)
0 2 4
% of cells
25 50 75 100
c
Tissue-resident macrophage differentially expressed genes
Hematopoietic precursor
GMP
Monocyte
Tissue-resident
Macrophage
Pit Bon BM Blo Spl Lun
Pit Bon BM Lun
Pit Fat MG Bon BM Blo Spl Lun Liv Pan Kid Bla
Cor Ste Pit Ret Bon Lun alv. Lun int. Liv MS4A7- Liv MS4A7+ SI MS4A7- SI MS4A7+ Spl
MRC1+
Fat LM Ton Kid Bla Fat MG BM Kid Ute Tes
CNS
ACY3
[CD300C]
BHLHE41
TMEM119
PTPRZ1
Bone
CKB
SPP1
RAB13
LAG3
SLC25A4
Lung alv.
ABCG1
CYFIP2
GPD1
CHIT1
LTA4H
Lung int.
[HLA-DRB1L]
[HLA-DQA2]
FCGR2B
LYZ
[CCL3L]
Liver MS4A7-
APOC3
ALDOB
APOC1
[TLL]
PRNP
Liver & SI MS4A7+
[SIGLEC8]
[SIGLEC7]
MS4A7
[FCGR3AL]
CD22
SI MS4A7-
MMP1
MMP10
TNFRSF6B
DNASE1L3
JAML
Spleen
[KRT76L]
CD5L
CCR3
[PRXL2A]
MMP12
Fat, LM, Tongue, Kidney, Bladder MRC1+
STAB1
ABCA6
GAS6
CLEC10A
MRC1
ln(UP10K+1)
0 2 4
% of cells
25 50 75 100

Fig. S9 - continued

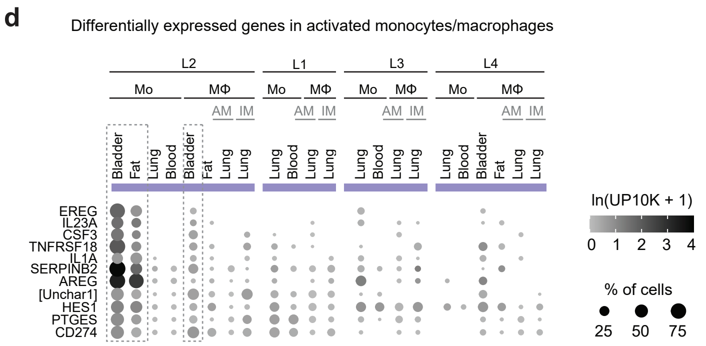


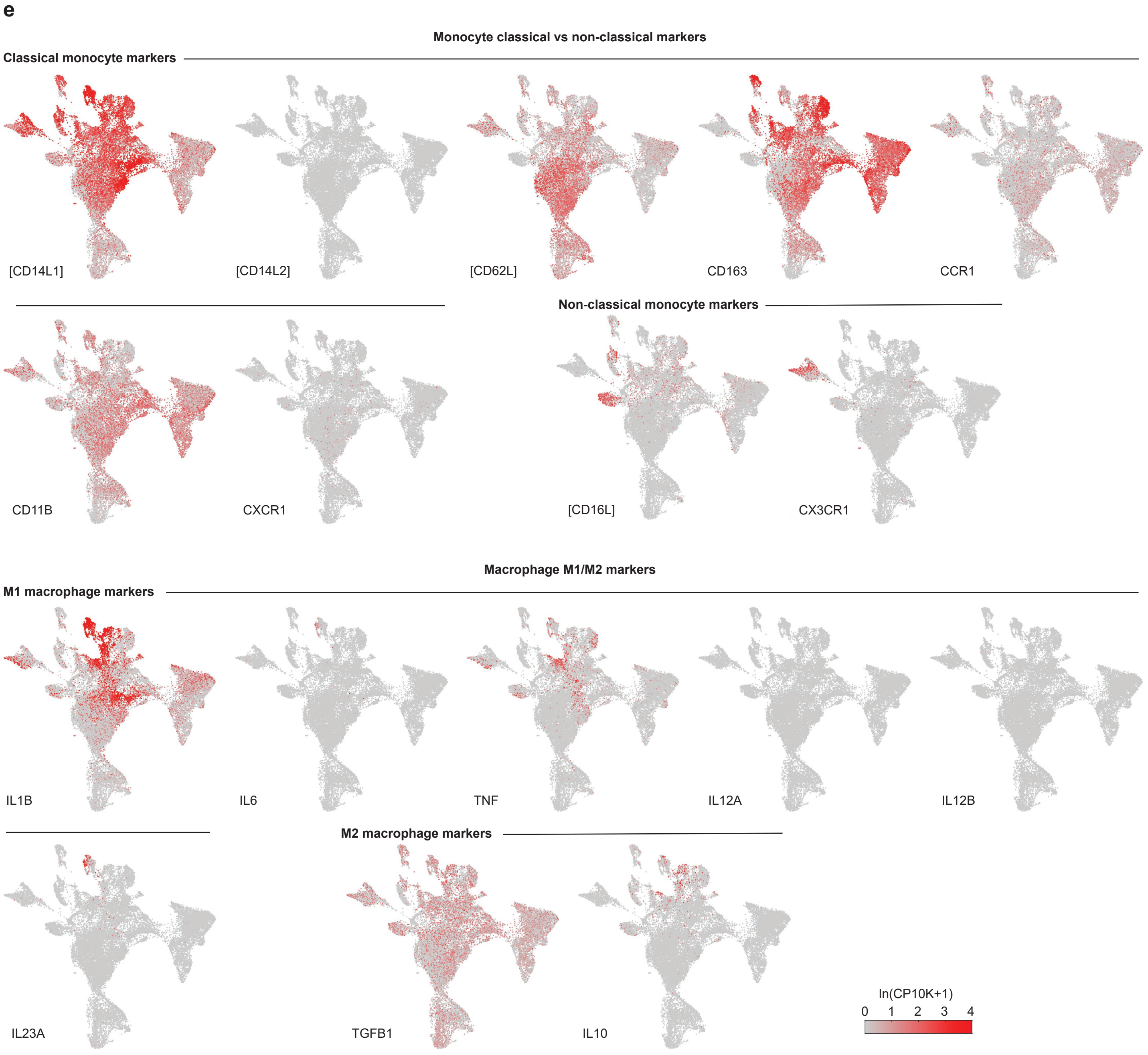

Fig. S9 - continued

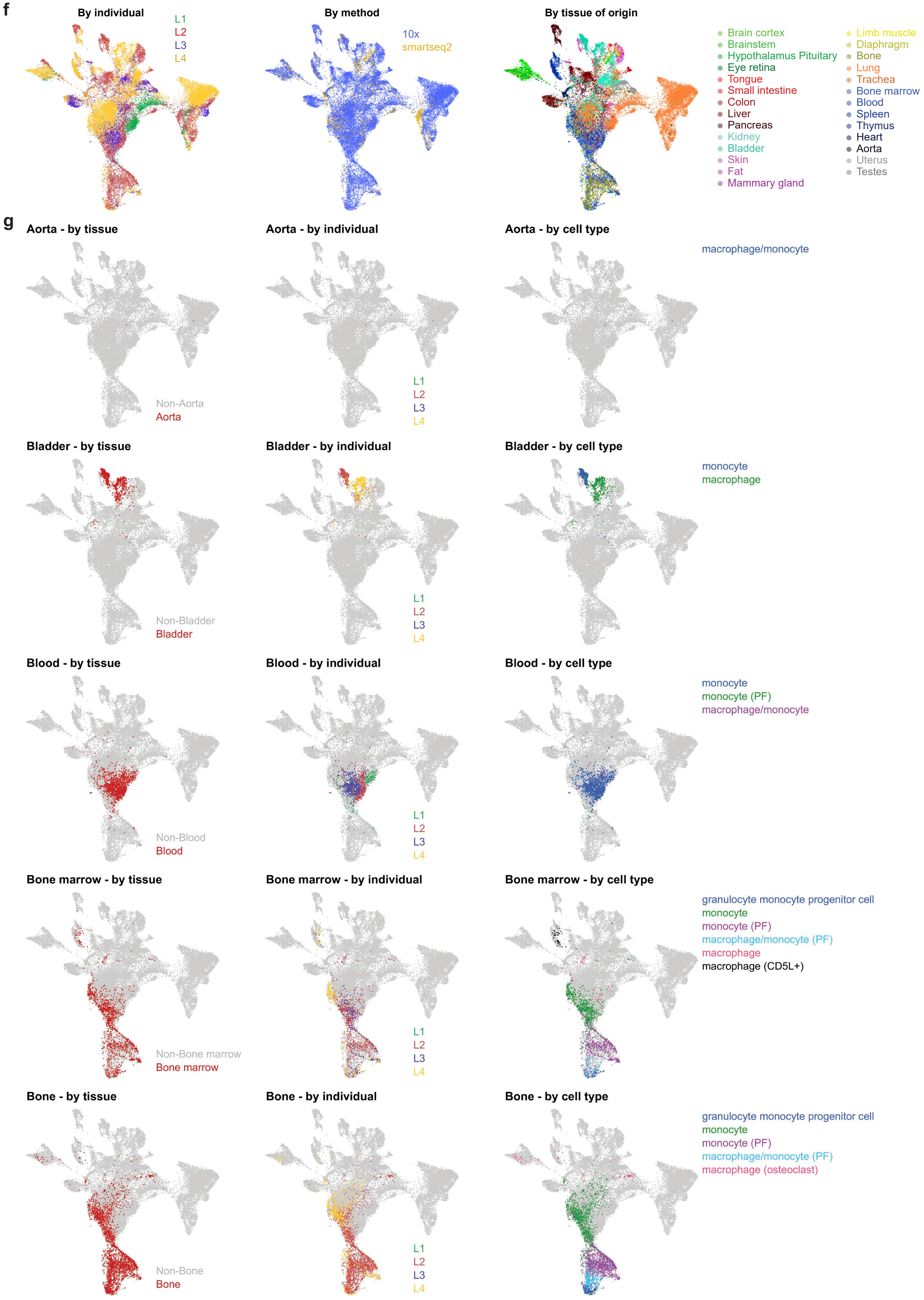

**Fig. S9 - continued**

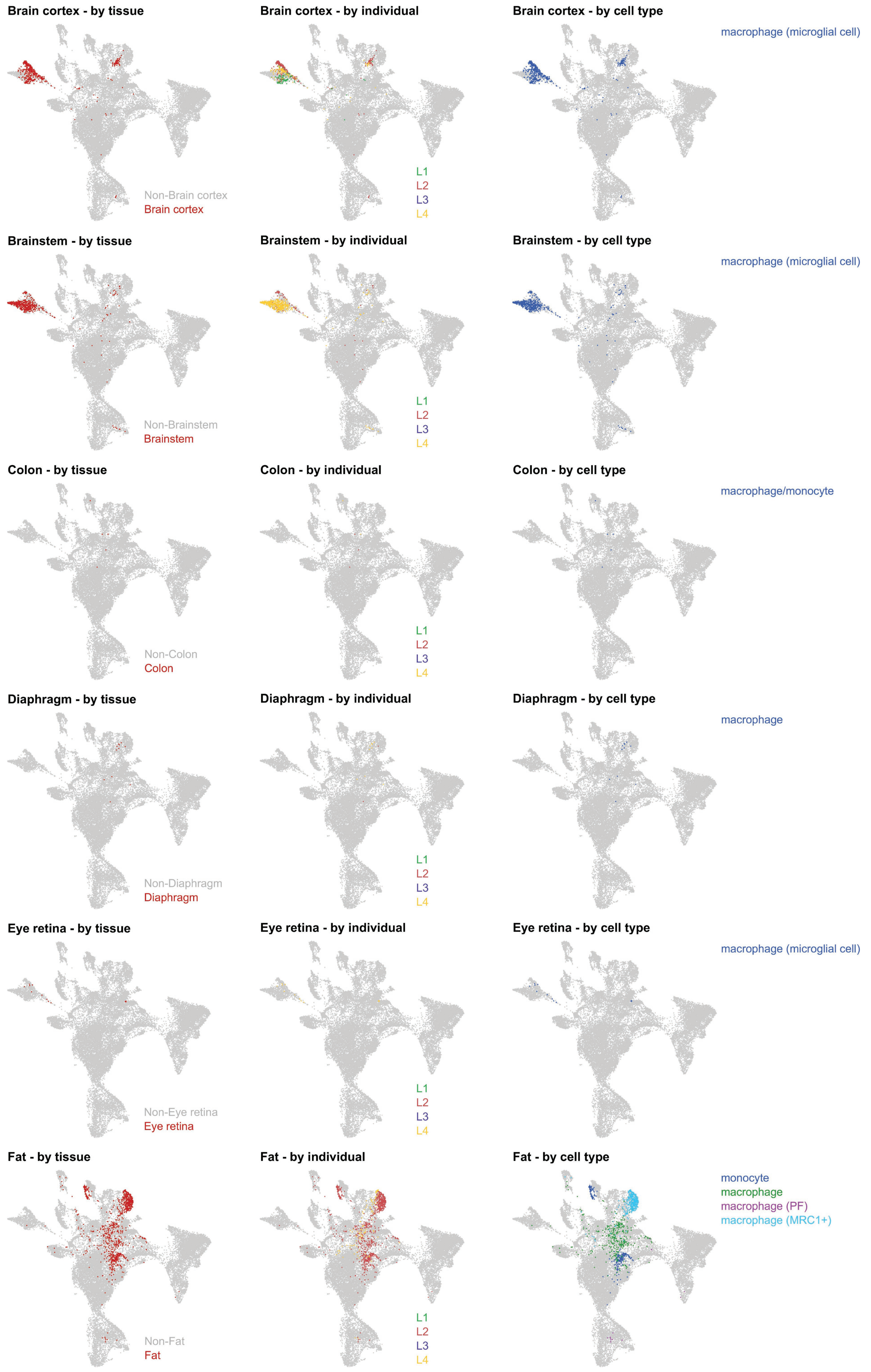

**Fig. S9 - continued**

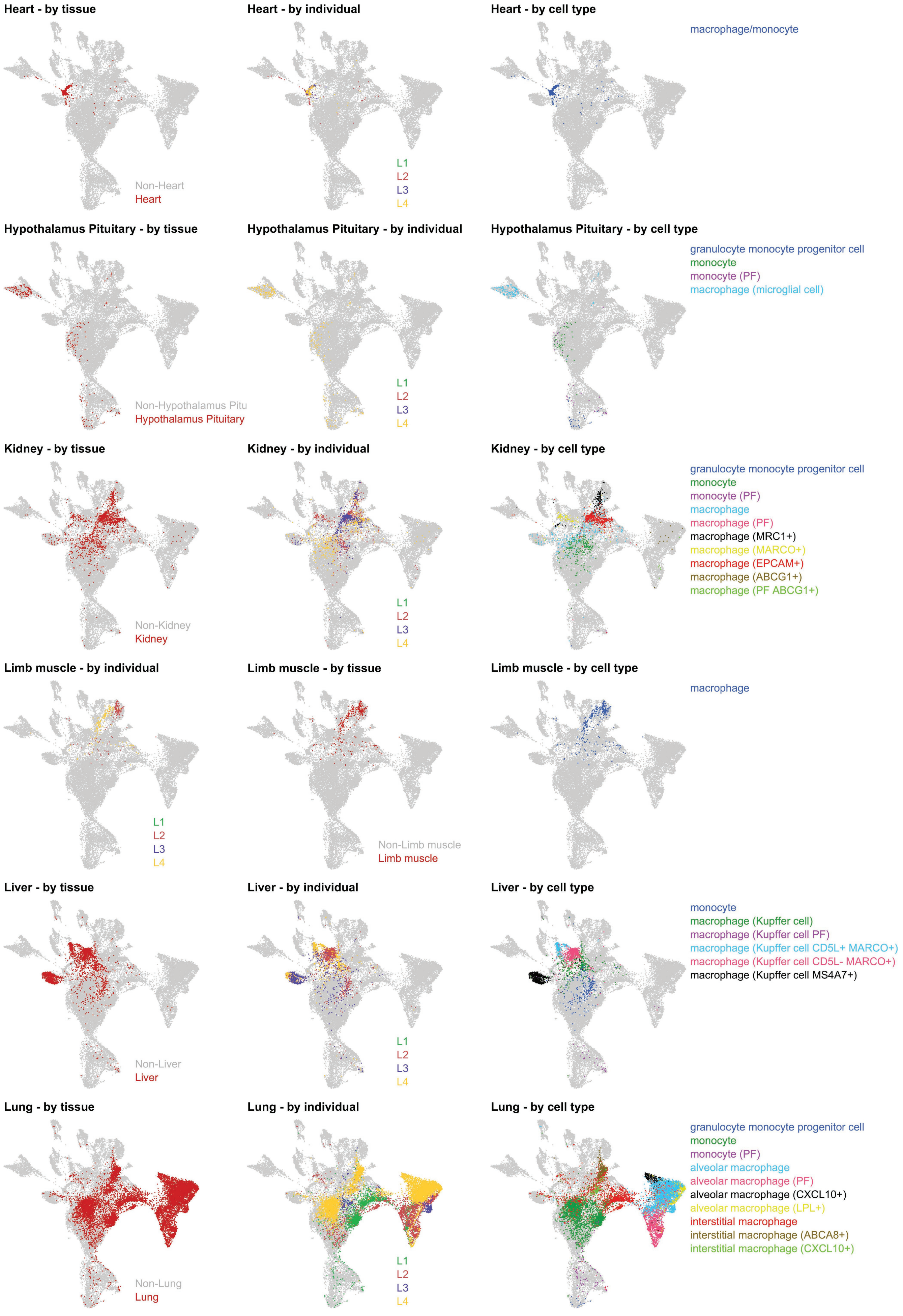

### Fig. S9 - continued

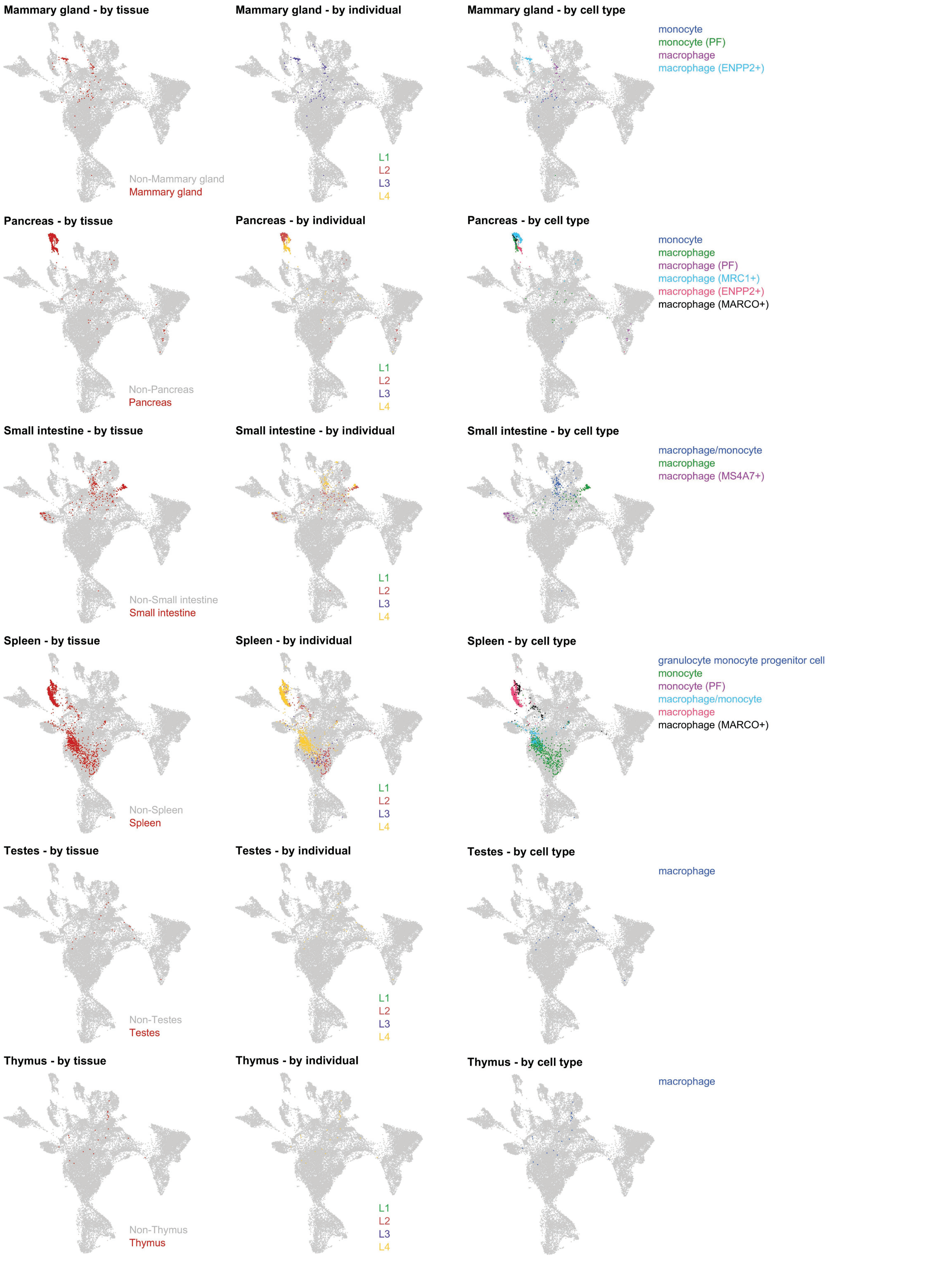

### Fig. S9 - continued

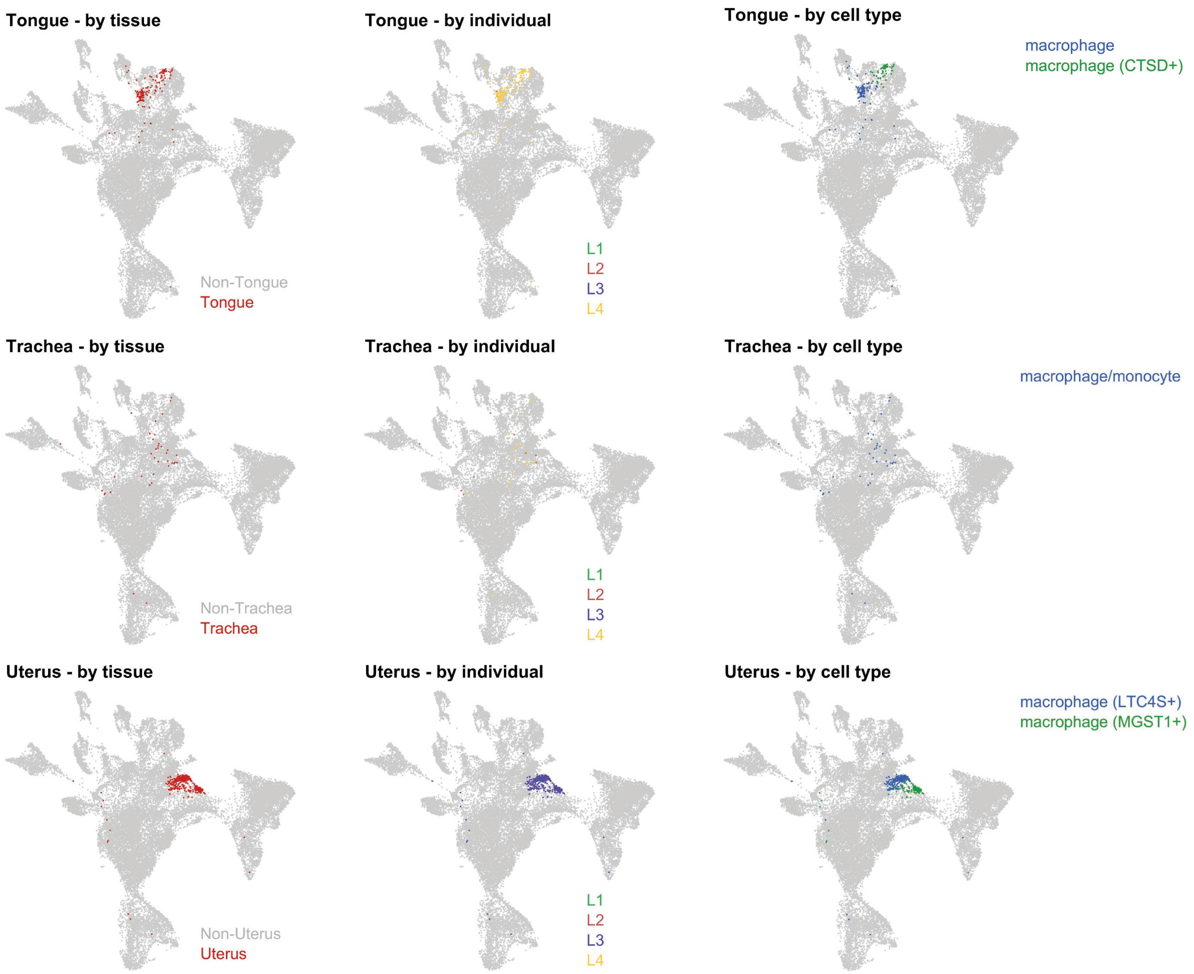

**Fig. S10**

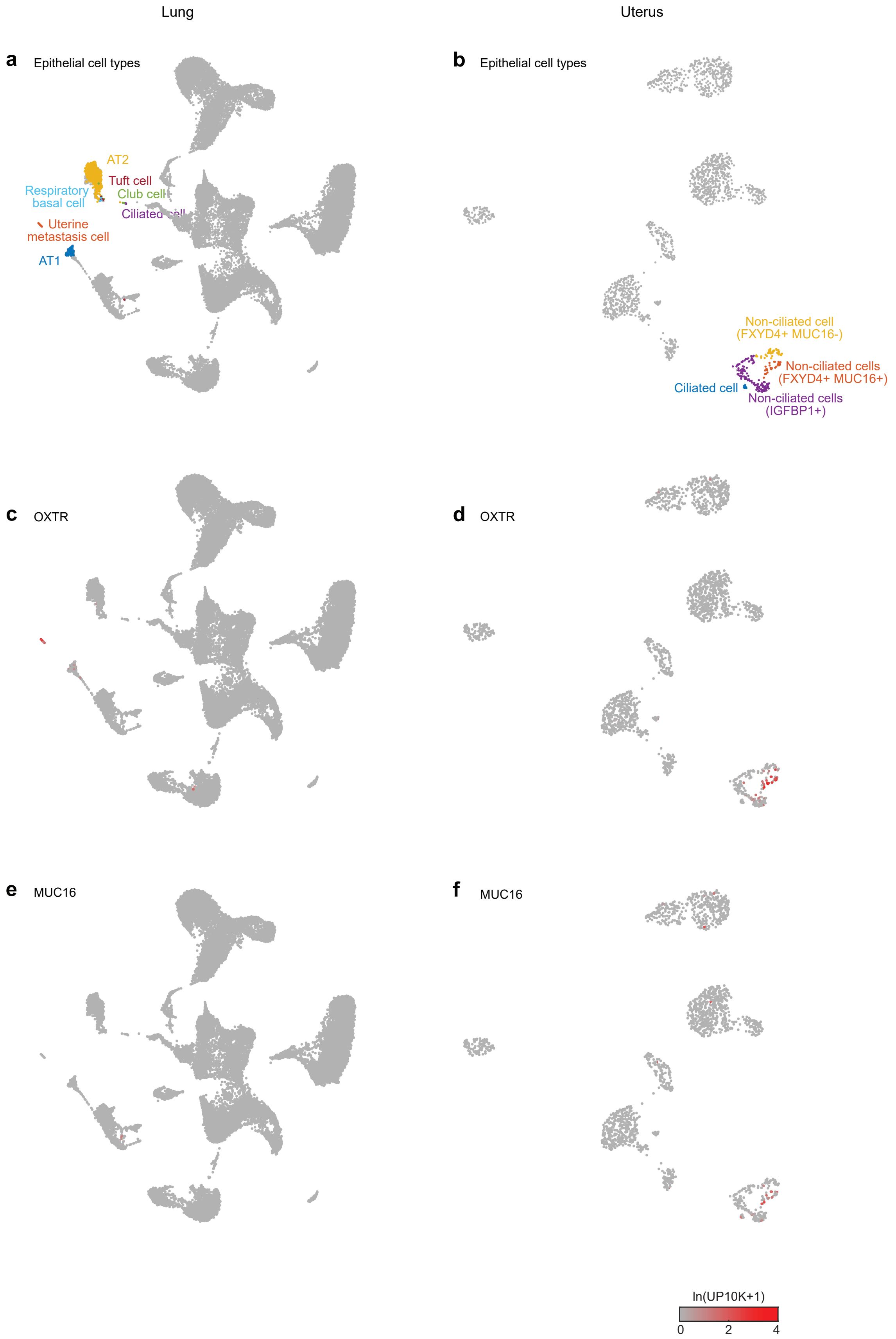

Lung
Uterus
a
Epithelial cell types
AT2
Tuft cell
Respiratory basal cell
Club cell
Uterine metastasis cell
AT1
b
Epithelial cell types
Non-ciliated cell (FXYD4+ MUC16-)
Non-ciliated cells (FXYD4+ MUC16+)
Ciliated cell
Non-ciliated cells (IGFBP1+)
c
OXTR
d
OXTR
e
MUC16
f
MUC16
ln(UP10K+1)
0
2
4

Fig. S11

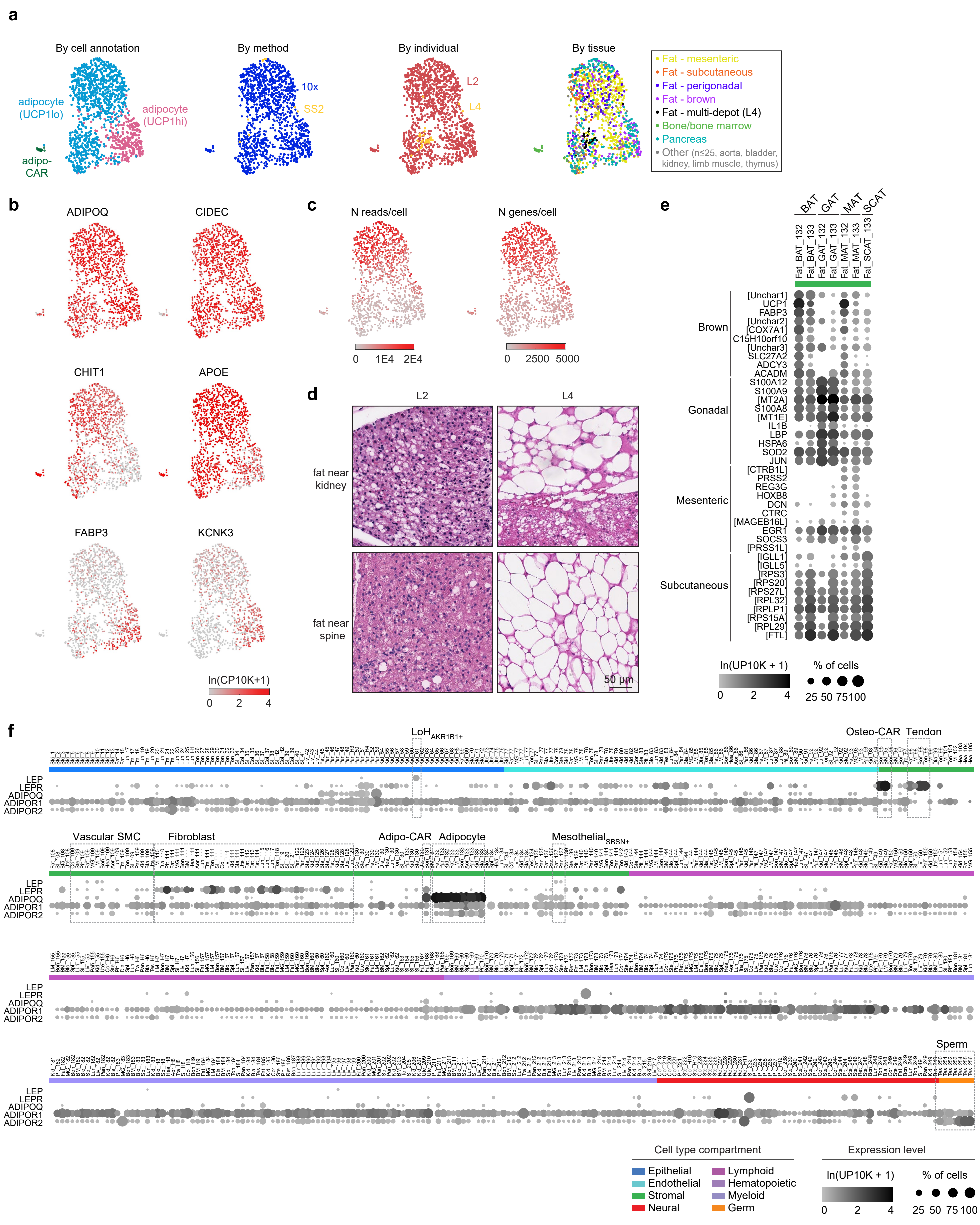
a
By cell annotation
adipocyte (UCP1lo)
adipocyte (UCP1hi)
adipo-CAR
By method
10x
SS2
By individual
L2
L4
By tissue
Fat - mesenteric
Fat - subcutaneous
Fat - perigonadal
Fat - brown
Fat - multi-depot (L4)
Bone/bone marrow
Pancreas
Other (n≤25, aorta, bladder, kidney, limb muscle, thymus)
b
ADIPOQ
CIDEC
CHIT1
APOE
FABP3
KCNK3
ln(CP10K+1)
0 2 4
c
N reads/cell
0 1E4 2E4
N genes/cell
0 2500 5000
d
L2
L4
fat near kidney
fat near spine
50 μm
e
BAT
GAT
MAT
SCAT
Fat_BAT_132
Fat_BAT_133
Fat_GAT_132
Fat_GAT_133
Fat_MAT_132
Fat_MAT_133
Fat_SCAT_133
Brown
[Unchar1]
UCP1
FABP3
[Unchar2]
[COX7A1]
C15H10orf10
[Unchar3]
SLC27A2
ADCY3
ACADM
Gonadal
S100A12
S100A9
[MT2A]
S100A8
[MT1E]
IL1B
LBP
HSPA6
SOD2
JUN
Mesenteric
[CTRB1L]
PRSS2
REG3G
HOXB8
DCN
CTRC
[MAGEB16L]
EGR1
SOCS3
[PRSS1L]
Subcutaneous
[IGLL1]
[IGLL5]
[RPS3]
[RPS20]
[RPS27L]
[RPL32]
[RPLP1]
[RPS15A]
[RPL29]
[FTL]
ln(UP10K + 1)
0 2 4
% of cells
25 50 75 100
f
LoH AKR1B1+
Osteo-CAR
Tendon
Vascular SMC
Fibroblast
Adipo-CAR
Adipocyte
Mesothelial SBSN+
Sperm
LEP
LEPR
ADIPOQ
ADIPOR1
ADIPOR2
Cell type compartment
Epithelial
Endothelial
Stromal
Neural
Lymphoid
Hematopoietic
Myeloid
Germ
Expression level
ln(UP10K + 1)
0 2 4
% of cells
25 50 75 100

**a.** Heatmap of PS gene expression - ordered by compartment, cell type, tissue

**Fig. S12**

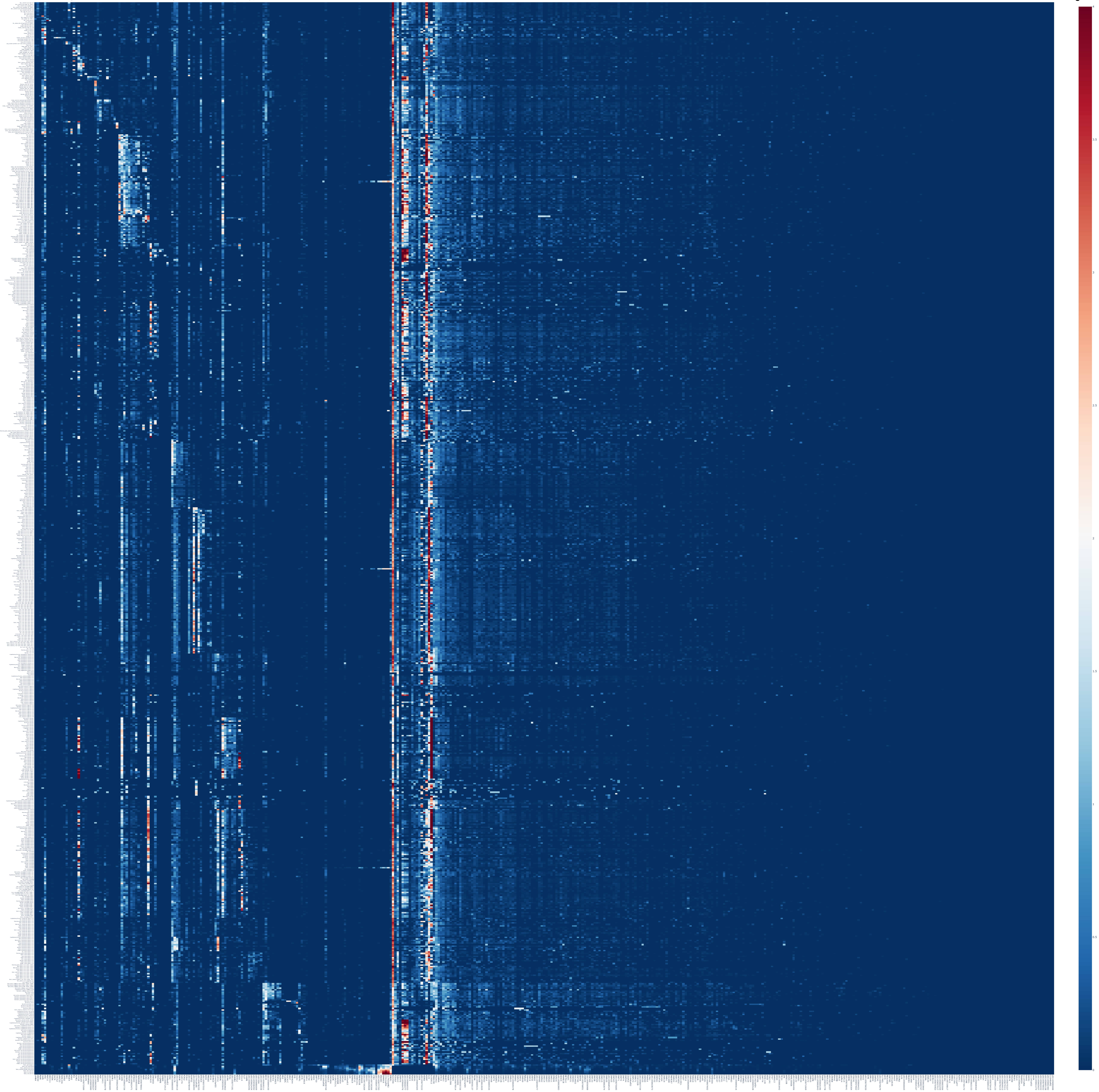

**b.** Heatmap of PS gene expression - ordered by tissue, compartment, cell type

**Fig. S12 - continued**

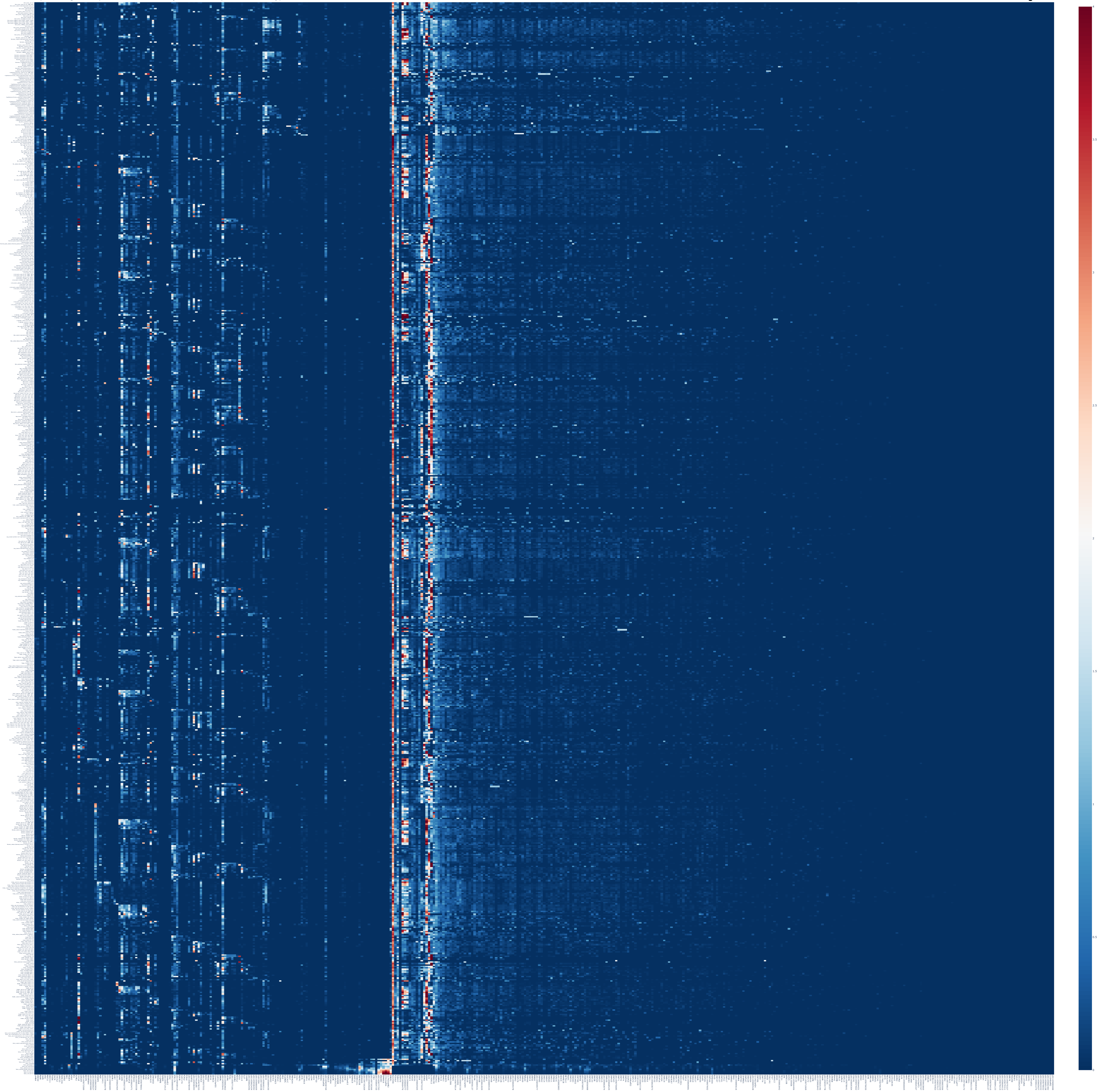

**c.** Dotplot of PS genes enriched in the epithelial compartment

**Fig. S12 - continued**

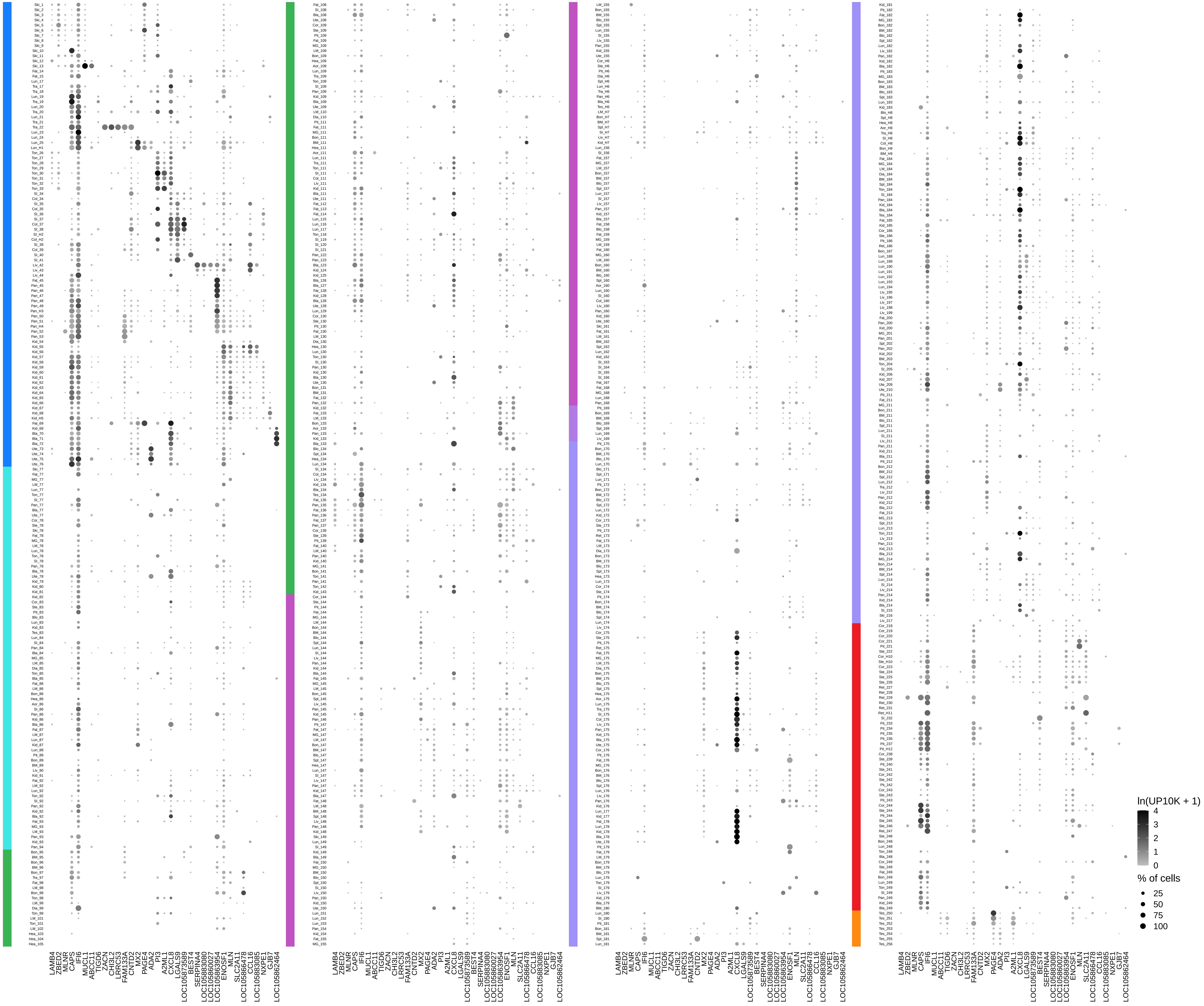

**d.** Dotplot of PS genes enriched in the endothelial compartment

**Fig. S12 - continued**

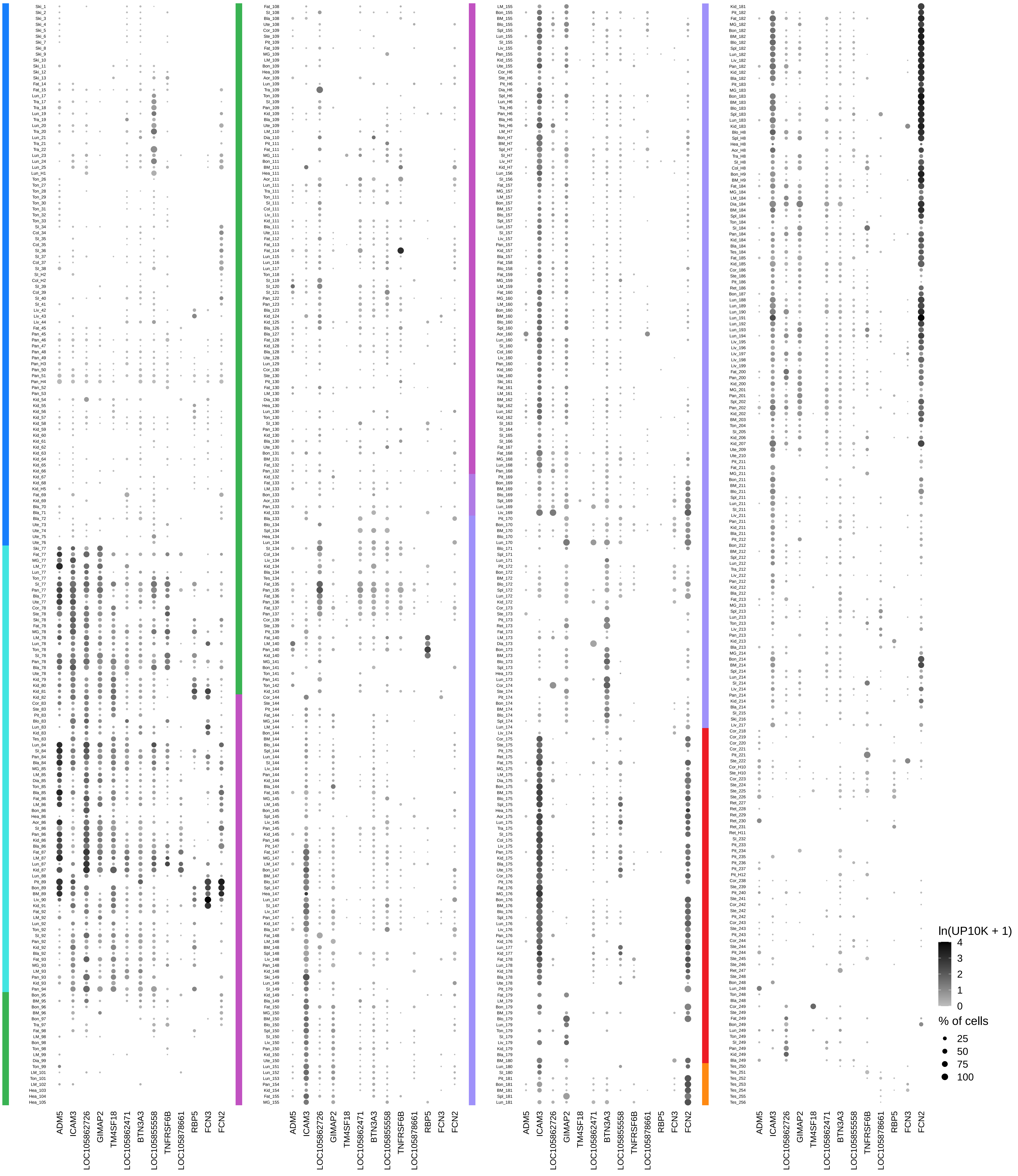

**e.** Dotplot of PS genes enriched in the stromal compartment

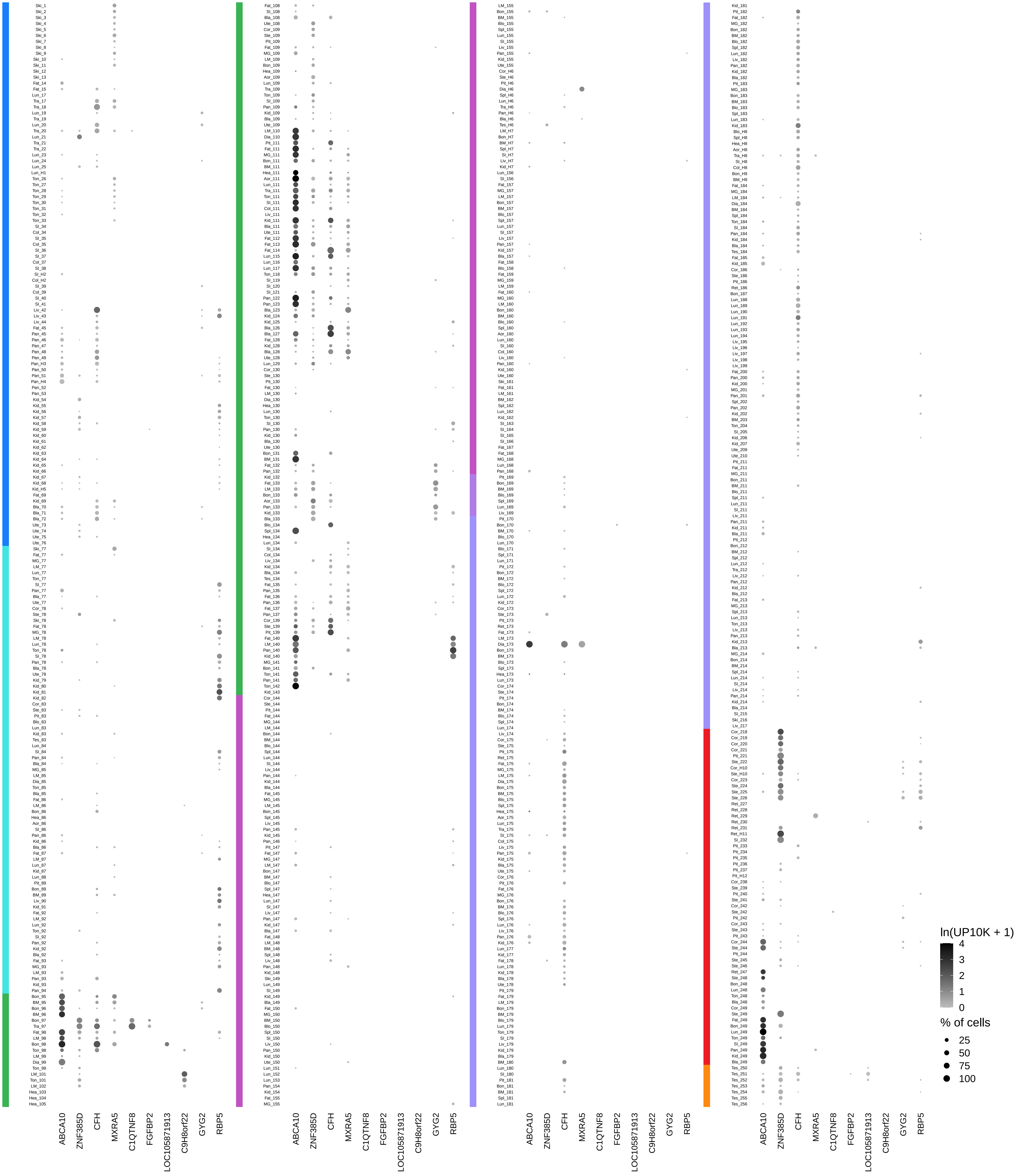

Fig. S12 - continued

f. Dotplot of PS genes enriched in the immune compartments

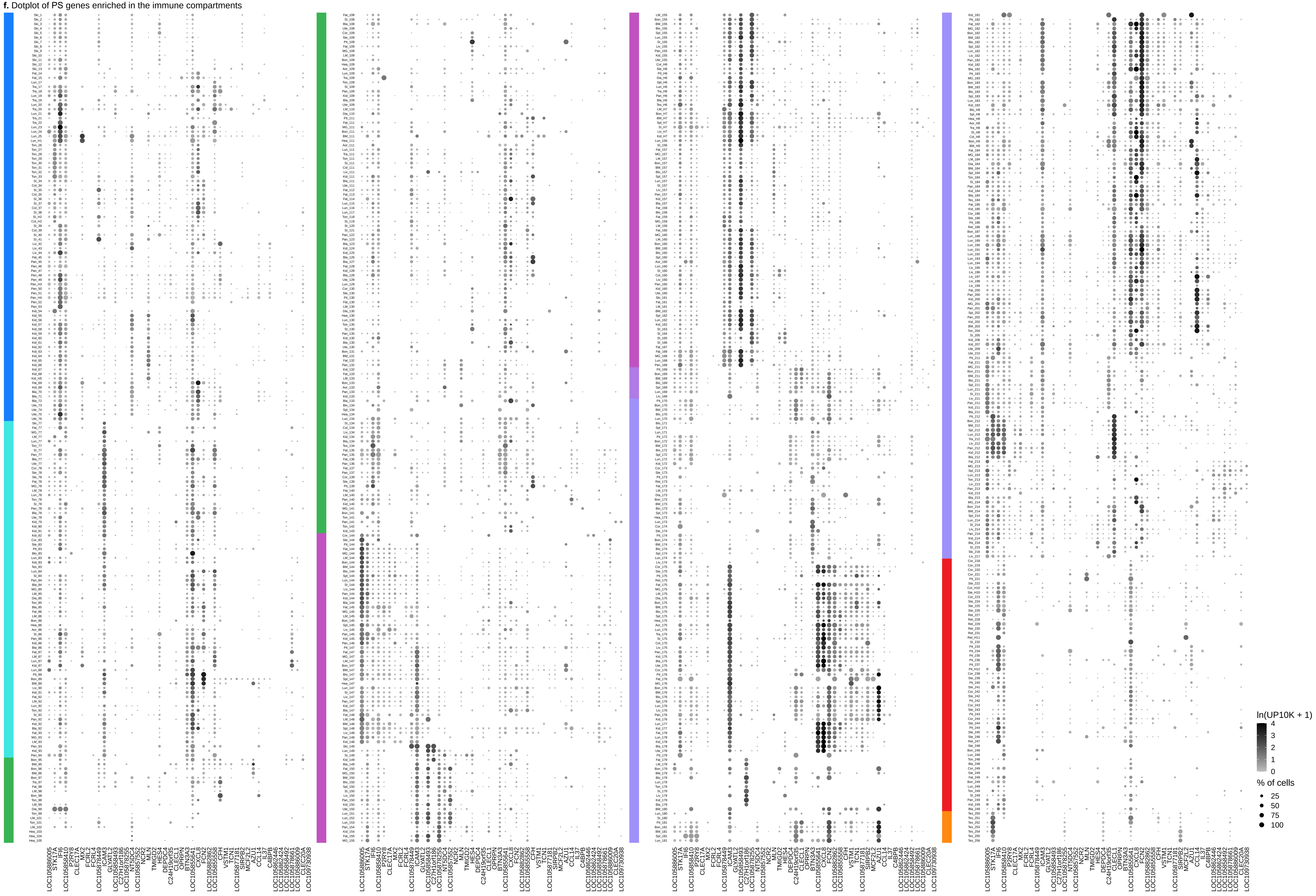

**g.** Dotplot of PS genes enriched in the neural compartment

**Fig. S12 - continued**

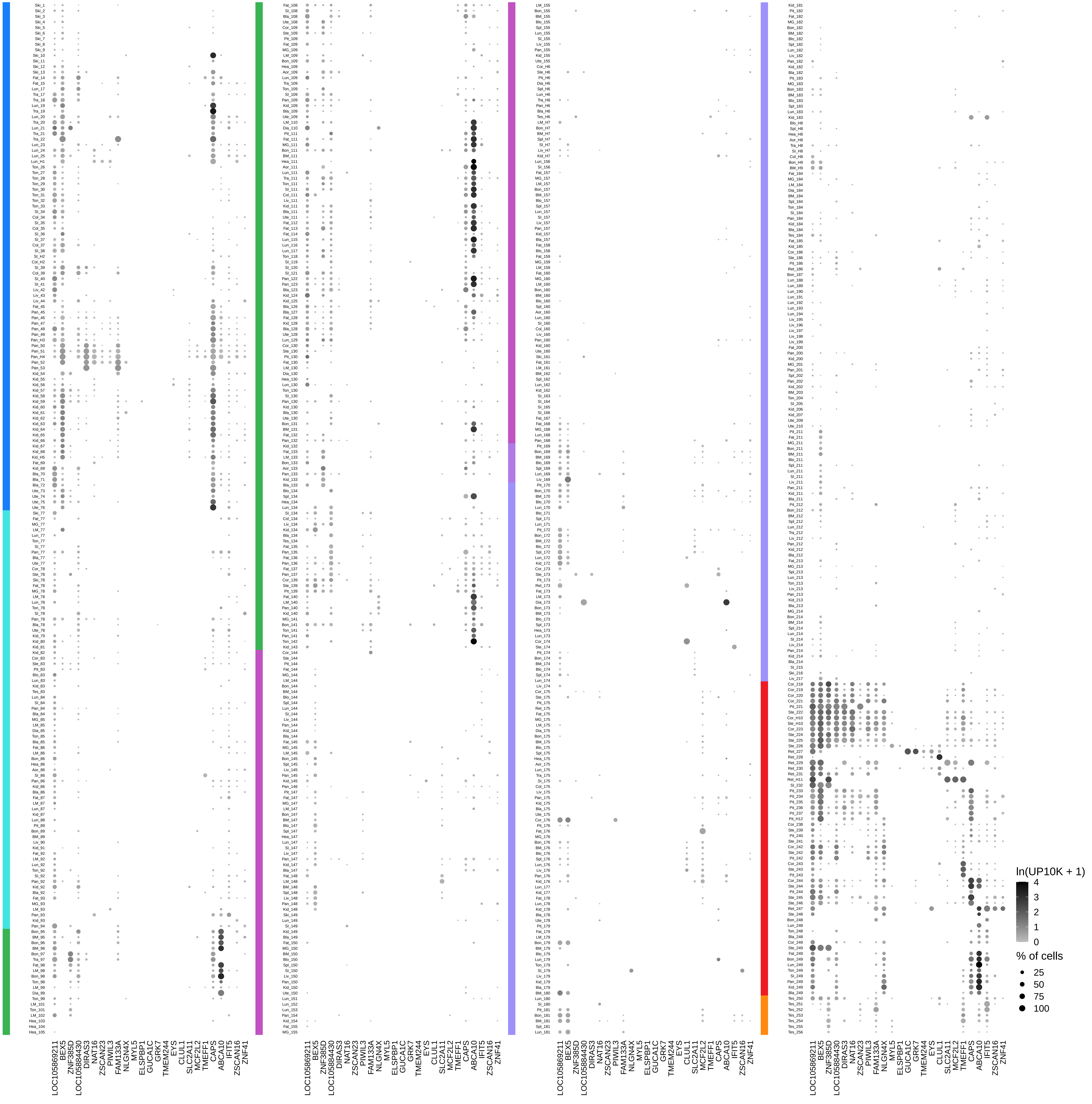

**h.** Dotplot of PS genes enriched/depleted in the germ compartment

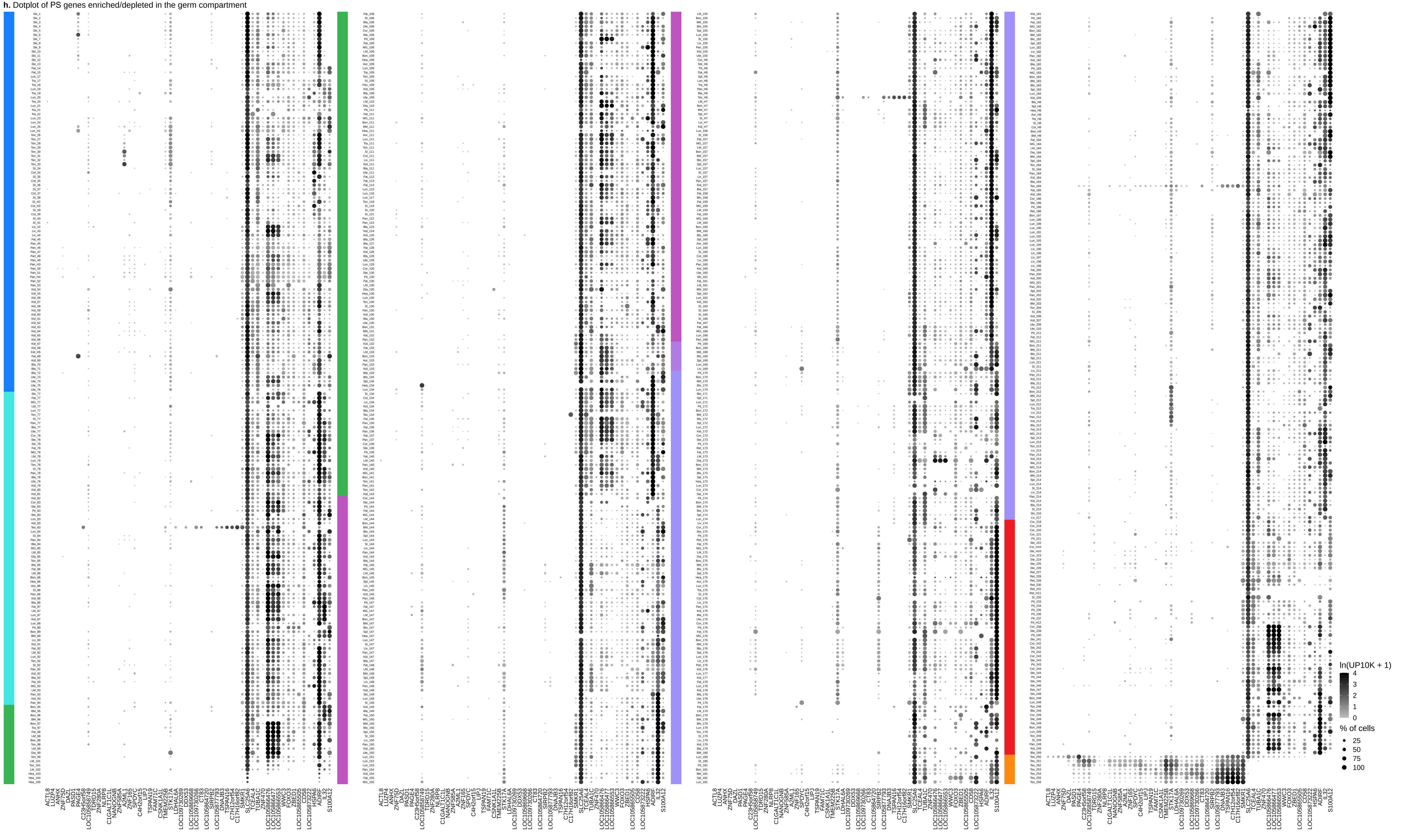

### Fig. S12 - continued

**i** Expression pattern in human vs. lemur

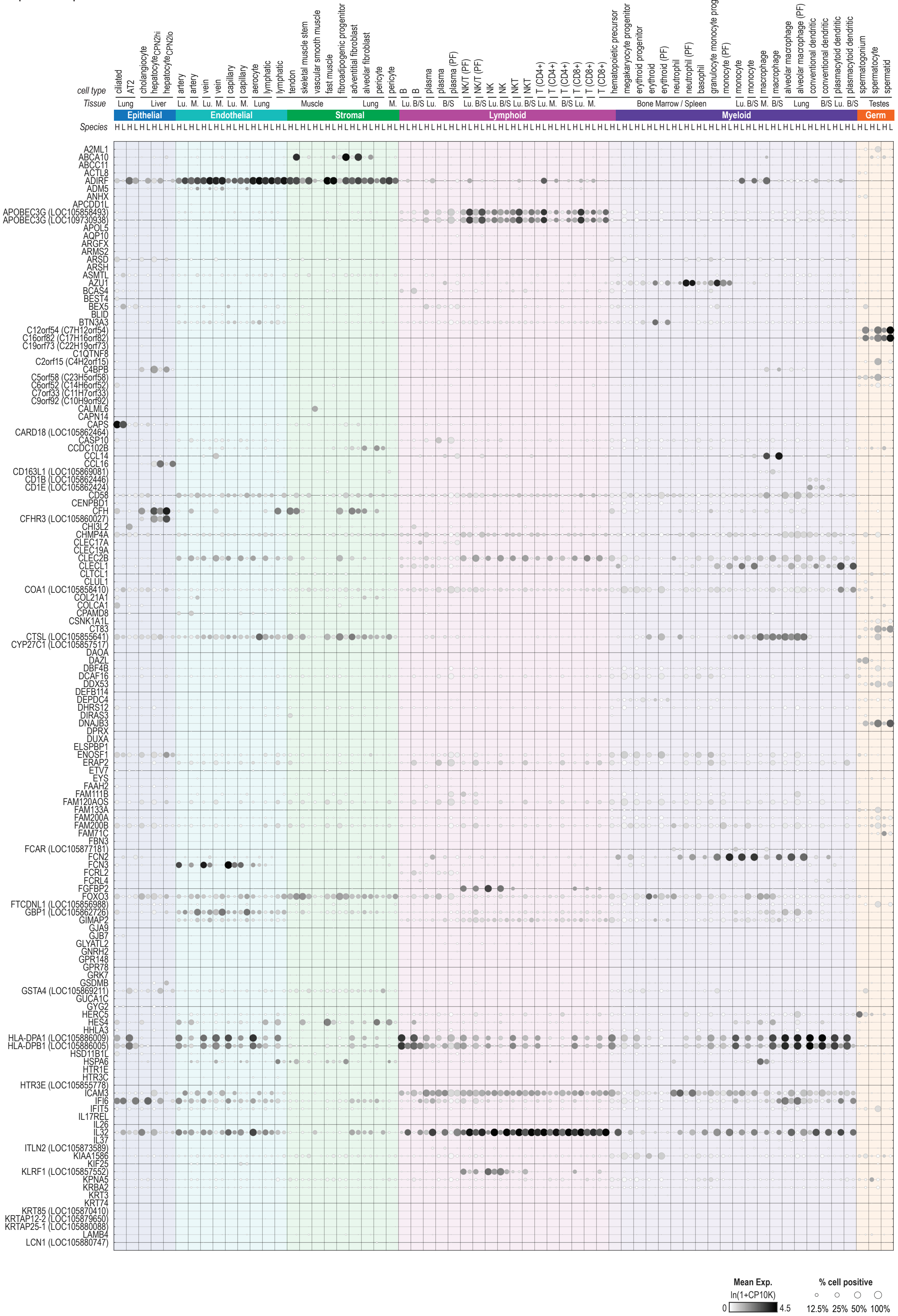

### Fig. S12 - continued

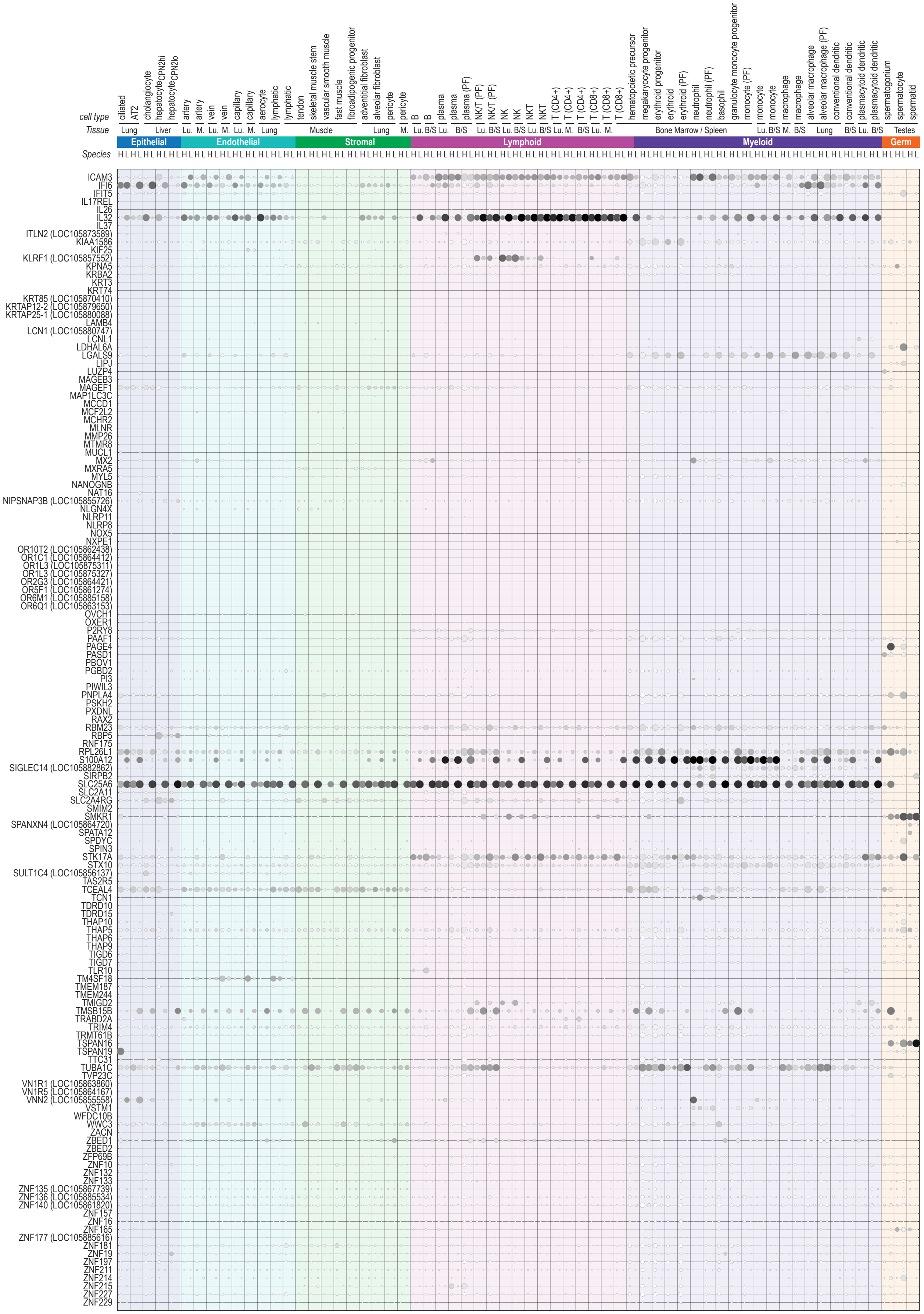

#### Fig. S12 - continued

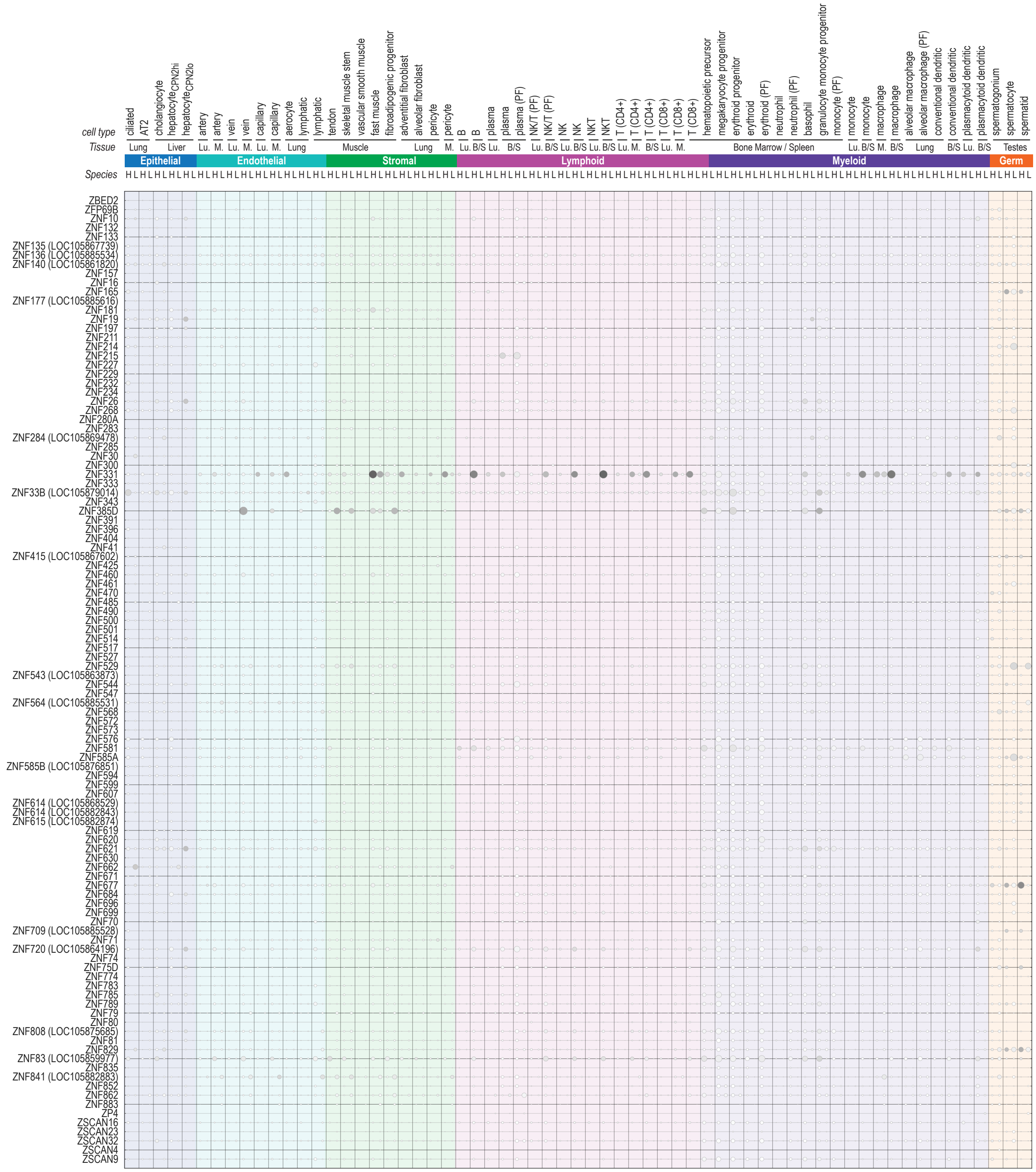

### Fig. S13

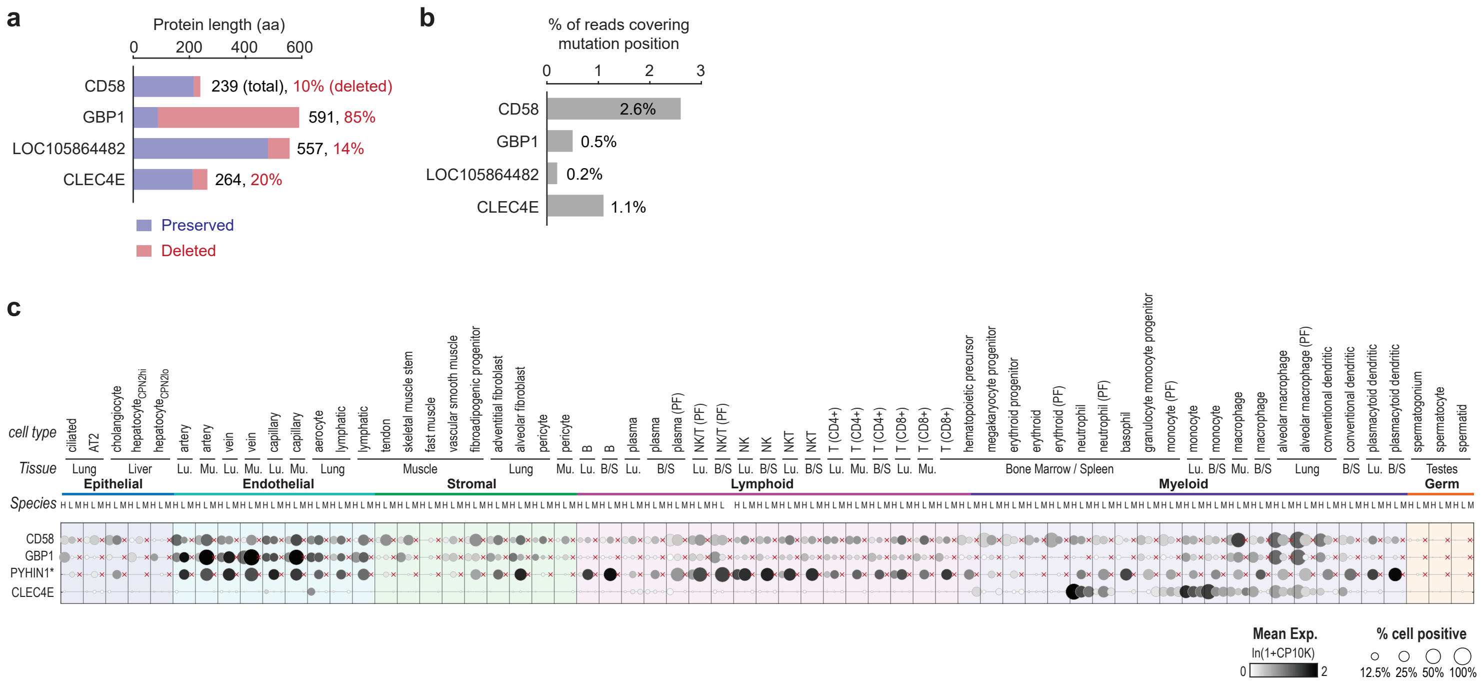
a
Protein length (aa)
0 200 400 600
CD58
239 (total), 10% (deleted)
GBP1
591, 85%
LOC105864482
557, 14%
CLEC4E
264, 20%
Preserved
Deleted
b
% of reads covering mutation position
0 1 2 3
CD58 2.6%
GBP1 0.5%
LOC105864482 0.2%
CLEC4E 1.1%
c
cell type
Tissue
Species
Epithelial
Endothelial
Stromal
Lymphoid
Myeloid
Germ
CD58
GBP1
PYHIN1*
CLEC4E
Mean Exp.
ln(1+CP10K)
0 2
% cell positive
12.5% 25% 50% 100%
