## Supplementary results for "Mouse lemur transcriptomic atlas informs primate genes, mutations, physiology, and disease"

### SUPPLEMENTARY RESULT

#### MHC annotation

We used our organism-wide atlas to enhance gene annotation of the major histocompatibility complex (MHC). MHC class II genes encode structurally-related alpha (A genes) and beta (B genes) subunits of transmembrane heterodimers expressed on dendritic and other professional antigen-presenting cells. They are loaded with exogenous peptides derived from phagocytosed pathogens and proteins, which serve as ligands for TCRs on helper T cells to initiate an adaptive immune response(Rock et al., 2016). Our allele-specific analysis of expression of MHC class II genes across the full atlas shows that each individual mouse lemur expresses one or two alleles of each MHC class II gene except *DOA* and *DOB*, which were not expressed in any of them, supporting their designation as pseudogenes based on sequence analysis(Averdam et al., 2011) (Fig. S5a-c). The other class II genes all appear to be single, non-duplicated genes, including two (*DQA*, *DQB*) that are currently annotated as duplicated and one (*DRB*) as triplicated, the latter from failed assembly of two genomic fragments of the second *DRB* allele(Guethlein et al., 2022) (Fig. S5e). *DQA*, *DQB*, *DRA*, and *DRB* are expressed at higher levels and in more cells than the other class II genes (*DMA*, *DMB*, *DPA*, *DPB*)(Guethlein et al., 2022) (Fig. S5h).

MHC class I genes encode the variable alpha chain that together with beta-2 microglobulin form widely-expressed transmembrane heterodimers that are loaded with peptides derived from cytosolic proteins, which serve as ligands for TCRs on cytotoxic T cells, or for KIRs (killer cell immunoglobulin-like receptors) or KLRs (killer cell lectin-like receptors) on NK cells, to destroy cells expressing viral, tumor or other "non-self" proteins(Djaoud & Parham, 2020; Rock et al., 2016). For MHC class I genes, no expression was detected in our atlas of genes in the canonical MHC locus at chromosome 6 (Fig. S5a), supporting their assignment as pseudogenes from sequence analysis(Averdam et al., 2009). All of the expressed mouse lemur class I genes mapped to a separate, duplicated cluster on chromosome 20q (Fig. S5b), an organization unique to mouse lemur(Guethlein et al., 2022) (Fig. S5d). Our allele-specific expression analysis identified 11 expressed genes in the cluster, four (*Mimu-168, -W03, -W04, -249*) we designate as "classical" based on their high and widespread expression, and the rest we designate "non-classical" (*Mimu-180ps, -191, -202, -208, -218, -229ps, -239ps*) including three previously thought to be pseudogenes (*Mimu-180ps, -229ps, -239ps*) based on sequence analysis (Fig. S5c, f-h).

Given the polymorphic nature of the MHC complexes(Norman et al., 2017), it remains challenging to distinguish some genes from allelic variation. Three of the classical genes (*Mimu-168, -W03, -W04*) are very similar in sequence and appear to be divergent allelic lineages, with some individuals (L1, L3) missing (or not expressing) one or more of the three genes. Three of the non-classical genes (*Mimu -208, -218, -229ps*) are also missing transcripts in two or three individuals (Fig. 4c). The current genome assembly (Mmur 3.022) contains gaps separating the complex into three sequence islands, rearrangement of which better matches the available BAC sequence(Averdam et al., 2009) (Fig. S5c), as detailed in Guethlein et al.(Guethlein et al., 2022); long range sequencing is needed to reveal the full number and order of genes for each individual.

#### Summary of mouse lemur clinical conditions

Details of the clinical conditions for the lemurs, including histopathological diagnosis, complete blood count (CBC) and complete metabolic panel (CMP) at time of euthanasia, necropsy report, and medical history were described in companion manuscripts(Casey et al., 2021; The Tabula Microcebus Consortium, 2021). Histology of all tissues in each individual can be viewed at the Tabula Microcebus web portal. Below we summarize clinical conditions relevant to this study:

L1 (male): widespread fibrous osteodystrophy with pathologic fractures; kidneys with bilateral acute tubular necrosis and pre-existing chronic renal disease; suppurative rhinitis, pharyngitis and tracheitis, as well as pulmonary emphysema.

L2 (female): mild fibrous osteodystrophy without pathologic fracture; kidneys with bilateral chronic renal disease (hydronephrosis, interstitial nephritis, and amyloidosis); uterus with diffuse uterine adenocarcinoma with suppurative metritis with tissue swab culture positive for *Klebsiella* species, *Enterococcus faecalis*, and aerobic gram-positive rods; lung with metastatic uterine adenocarcinoma with necrosis and hemorrhage, as well as multifocal bronchopneumonia with tissue swab culture positive for *Klebsiella* species; sinusitis and rhinitis; bladder with suppurative cystitis.

L3 (female): mild fibrous osteodystrophy without pathologic fracture; kidneys with bilateral mild diffuse glomerulopathy and interstitial nephritis/fibrosis; uterus with diffuse uterine adenocarcinoma with necrotic cysts; a 2-cm abdominal mass (suspected mesenteric lymph node) and a peri-adrenal lymph node with metastatic uterine adenocarcinoma.

L4 (male): mild fibrous osteodystrophy without pathologic fracture; kidneys with bilateral moderate diffuse glomerulopathy, interstitial nephritis/fibrosis and renal pelvis hemorrhage; severe pulmonary hemorrhage.

#### Analysis of chemokine receptors and ligands

We examined the expression of chemokine ligands and receptors across the atlas (Fig. 4a, Fig. 6a-f). While the ligands were broadly expressed across non-germ compartments, the receptors were mostly restricted to the immune populations (Fig. S7a). Macrophages expressed the highest number of chemokine ligands and receptors, supporting their critical role in directing immune cell trafficking (Fig. S7a-b). Activated neutrophils of the uterus (*IL18BP+*) also expressed a large number of chemokine receptors, suggesting their role in responding to inflammation. Linking chemokine-expressing cell types to the cell types expressing the corresponding receptor revealed a global cell-cell interaction network (Fig. S7d) with a moderate density (i.e., 6.8% of all possible interactions were present). Though as expected, the network was much more densely connected between immune cells (density = 33.5%) compared to between immune and non-immune cells (3.3%) and between non-immune cells (1.4%). There were also dramatic differences in the cell specificity of different chemokine signaling pairs, with some resulting in widespread connections across a tissue and others connecting only a few specific cell types (see the example of lung and blood in Fig. S7e).

Chemokines and their receptors showed a highly cell type specific expression pattern across the atlas (Fig. 4a, Fig. S7c, f). For example, each major type of dendritic cells (DC) in the atlas expresses a unique combination of chemokine ligands and receptors: conventional DCs (*XCR1+*), plasmacytoid DCs (*CXCL16-*), mature DC (*CCL19+*, *CCL22+*, *CCR7+*), and *IGSF6+* DCs (*CXCR3-*).

The analysis also suggested patterns of cell-cell chemoattraction (Fig. 4a). In addition to the examples highlighted in Section 4, we identified lemur B cells, *CD4+* T cells, and mature DCs selectively expressing the receptor *CCR7*, and thus could be directed by the corresponding ligand expressing lymphatic endothelial cells (*CCL21*) and reticular cells (*CCL21*, *CCL19*) to lymph nodes(Barone et al., 2016; Farnsworth et al., 2019), as illustrated in Fig. 4b. This may be further aided by *ACKR4*, a decoy receptor for CCL19/CCL21, expressed in other lymphatic endothelial cells, and thus could enhance local gradients of CCL19 and CCL21 in and around lymph nodes(Jalkanen & Salmi, 2020). *ACKR4* was also expressed in skin keratinocytes, mesothelial cells, and lung alveolar macrophages, likely to scavenge soluble CCL19/21 and create a gradient in the tissue to guide local immune cell trafficking, as previously suggested in mice(Bryce et al., 2016; Ulvmar et al., 2014). Reticular cells also expressed *CXCL13* and *CXCL12*, which could help attract mature B cells that express the CXCL13 receptor (*CXCR5*) and other immune cells that express the CXCL12 receptor (*CXCR4*)(Fletcher et al., 2015; Nie et al., 2004). The expression patterns of *RARRES2* and its receptor (*CMKLR1*) suggest they play a similar role for monocytes and macrophages in lymph node trafficking(Vermi et al., 2005).

In addition to the stereotypical expression of chemokines and/or receptors across lemur individuals and tissues under normal physiology, we also uncovered likely examples of disease-associated expression of the chemokine pathways, consistent with the clinical data and histopathology in specific individuals and tissues(Casey et al., 2021). These elucidate the lemur inflammatory response (as exemplified in Fig. 4b). For example, the three "interferon gamma-induced chemokines" (*CXCL9*, *CXCL10*, *CXCL11*) were co-expressed at high levels in several diseased or inflamed tissues: alveolar and interstitial macrophages and capillary cells of L2's and L4's lung (Fig. S7e, Fig. 4a) (beset with uterine metastases and pulmonary hemorrhage, respectively), as well as subpopulations of capillary and mesothelial cells in L2's fat (Fig. 4a-b) – presumably recruiting or retaining the *CXCR3*-expressing major dendritic cell types (conventional, plasmacytoid, mature) and other leukocytes into the inflamed tissues (and possibly suppressing angiogenesis)(Kochumon et al., 2020; Metzemaekers et al., 2017). Similarly, a subset of activated neutrophils, monocytes, and macrophages (see below) express higher levels of the potent neutrophil chemoattractant *CXCL8* with its receptors *CXCR1* and *CXCR2* broadly expressed across all neutrophils, allowing recruitment of additional neutrophils and amplifying inflammation (Fig. 4a-b).

#### Subpopulations of activated neutrophils

Activated neutrophils were found across multiple tissues in three individuals, and clustered into two subpopulations (Fig. 5d, Fig. S8e-f). One group, designated *CCL13+*, expresses chemoattractants for monocytes and eosinophils (*CCL15, CCL13 (LOC105859268*), *CCL2-like* (*LOC105859340/LOC105885684*))(Gschwandtner et al., 2019; Mendez-Enriquez & García-Zepeda, 2013; Shimizu & Dobashi, 2012). This population was found in L2's lung (site of metastatic adenocarcinoma and pneumonia) and sparsely in the kidney (possibly affected by neighboring bladder infection/cystitis and/or underlying chronic kidney disease) as well as in L1's lung (site of emphysema and suppurative rhinotracheitis).

The other neutrophil group, designated *IL18BP+*, shows enriched expression of genes including *IL18BP*, *ANXA5*, *SQSTM1*, *CD9*, *PTGR1*, and *CCR5*, many not previously implicated in neutrophil biology (Fig. S8f). This population was found in L2's bladder (site of suppurative cystitis) and lung (site of metastatic adenocarcinoma and pneumonia), and sparsely in the kidney (site of chronic kidney disease) and perigonadal fat (possibly affected by neighboring uterine cancer and infection), L1's lung (site of emphysema and suppurative rhinotracheitis), and L3's uterus (site of adenocarcinoma).

Both groups express high levels of chemoattractants such as *CXCL8* that recruits *CXCR1/2*-expressing maturing neutrophils and *CCL5* (*RANTES*) that recruits *CCR5*-expressing activated neutrophils and other immune cells ( Fig. 4a-b, Fig. S8f). These activated neutrophils also showed downregulation of L-selected (*SELL/CD62L*), facilitating their extravasation from the blood into sites of inflammation(Ivetic et al., 2019) ( Fig. 4b, Fig. 5c).

**Analysis of monocytes/macrophages**

Monocytes and macrophages are phagocytic leukocytes that work with neutrophils to engulf and destroy microbial pathogens and abnormal cells such as tumor cells. Like neutrophils, monocytes formed a continuous and homogeneous trajectory emanating from hematopoietic precursor and granulocyte-monocyte progenitor (Fig. 5a, Fig. S9a-b), with an only exception of likely activated monocytes as described below. Interestingly, we did not otherwise detect biologically meaningful monocyte subclusters in any tissue or identify subpopulations using canonical markers that distinguish human or mouse monocyte subtypes(The Tabula Microcebus Consortium, 2021) (Fig. S9e).

In contrast, lemur macrophages were highly diverse, showing many distinct, tissue-specific subpopulations (Fig. S9a-c, f-g). Several of the tissue-specific subpopulations clustered near monocytes, suggesting they may arise from circulating monocytes and undergo tissue-specific maturation. Other subpopulations segregated from the monocyte trajectory, suggesting these are likely stable populations of tissue-resident macrophages. Indeed, many expressed genes known to mark tissue-resident macrophages in human and mouse (Fig. S9a-b, and see Table 1 in accompanying manuscript(The Tabula Microcebus Consortium, 2021)), such as CNS microglia (e.g., *CX3CR1*)(Hickman et al., 2013; Jurga et al., 2020), liver Kupffer cells (*ID3*)(Bonnardel et al., 2019), and lung alveolar macrophages (*PPARG*)(Davies et al., 2013; Evren et al., 2021; Todd et al., 2016). Bone macrophages specifically expressed osteoclast markers (*CTSK, ATP6V0D2* and *NFATC1*)(Tsukasaki et al., 2020), but they did not form a separate cluster. However, lemur macrophages cannot be distinguish into canonical M1 ("classically activated") vs M2 ("alternatively activated") macrophages based on commonly used markers(Italiani & Boraschi, 2014; Orekhov et al., 2019) (Fig. S9e), though this distinction in other systems has blurred(Martinez & Gordon, 2014).

Nevertheless, we identified previously uncharacterized macrophage subtypes. For example, we identified a population of liver and small intestine *MS4A7+* macrophages that co-clustered and were separated from other *MS4A7-* populations in these tissues, including the likely canonical intestinal resident macrophages (*CD163+* (*LOC105869074*), *TIMD4+*) and the majority of liver Kupffer populations that express canonical mouse macrophage marker *ADGRE1+* (*F4/80*) (Fig. S9a-c, f-g). We also identified a cluster of *MRC1+* macrophages present in multiple tissues (fat, bladder, tongue, kidney, limb muscle) whose transcriptomic profile did not resemble that of any known resident macrophages (Fig. S9a-c, f-g). These results exemplify additional instances of our expanding understanding of molecular diversity within macrophages(Aizarani et al., 2019; MacParland et al., 2018; Wu et al., 2020).

A small fraction of monocytes and macrophages may be activated in response to inflammation, including *CXCL10+* alveolar and interstitial macrophages of L2 and L4's lungs (see above chemokine section). We also identified a population of likely inflammatory monocytes in L2's bladder (site of suppurative cystitis) and perigonadal fat (possibly affected by nearby uterine adenocarcinoma and infection) that clustered near macrophages from the same tissues and separately from all other monocytes (Fig. S9a). These monocytes expressed inflammation-associated genes, including *CD274/PD-L1*, *IL23A*, *AREG*, *CSF3*, *IL1A*, possibly teaming up with the activated neutrophils in these tissues to battle the local infections (cystitis and cancer) (Fig. S9d).

Hickman, S. E., Kingery, N. D., Ohsumi, T. K., Borowsky, M. L., Wang, L.-C., Means, T. K., &

El Khoury, J. (2013). The microglial sensome revealed by direct RNA sequencing. *Nature Neuroscience*, *16*(12), 1896–1905.
Italiani, P., & Boraschi, D. (2014). From Monocytes to M1/M2 Macrophages: Phenotypical vs. Functional Differentiation. *Frontiers in Immunology*, *5*, 514.
Ivetic, A., Hoskins Green, H. L., & Hart, S. J. (2019). L-selectin: A Major Regulator of Leukocyte Adhesion, Migration and Signaling. *Frontiers in Immunology*, *10*, 1068.
Jalkanen, S., & Salmi, M. (2020). Lymphatic endothelial cells of the lymph node. *Nature Reviews. Immunology*, *20*(9), 566–578.
Jurga, A. M., Paleczna, M., & Kuter, K. Z. (2020). Overview of General and Discriminating Markers of Differential Microglia Phenotypes. *Frontiers in Cellular Neuroscience*, *14*, 198.
Kochumon, S., Madhoun, A. A., Al-Rashed, F., Azim, R., Al-Ozairi, E., Al-Mulla, F., & Ahmad, R. (2020). Adipose tissue gene expression of CXCL10 and CXCL11 modulates inflammatory markers in obesity: implications for metabolic inflammation and insulin resistance. *Therapeutic Advances in Endocrinology and Metabolism*, *11*, 2042018820930902.
MacParland, S. A., Liu, J. C., Ma, X.-Z., Innes, B. T., Bartczak, A. M., Gage, B. K., Manuel, J., Khuu, N., Echeverri, J., Linares, I., Gupta, R., Cheng, M. L., Liu, L. Y., Camat, D., Chung, S. W., Seliga, R. K., Shao, Z., Lee, E., Ogawa, S., … McGilvray, I. D. (2018). Single cell RNA sequencing of human liver reveals distinct intrahepatic macrophage populations. *Nature Communications*, *9*(1), 4383.
Martinez, F. O., & Gordon, S. (2014). The M1 and M2 paradigm of macrophage activation: time for reassessment. *F1000prime Reports*, *6*, 13.
Mendez-Enriquez, E., & García-Zepeda, E. A. (2013). The multiple faces of CCL13 in immunity and inflammation. *Inflammopharmacology*, *21*(6), 397–406.
Metzemaekers, M., Vanheule, V., Janssens, R., Struyf, S., & Proost, P. (2017). Overview of the Mechanisms that May Contribute to the Non-Redundant Activities of Interferon-Inducible CXC Chemokine Receptor 3 Ligands. *Frontiers in Immunology*, *8*, 1970.
Nie, Y., Waite, J., Brewer, F., Sunshine, M.-J., Littman, D. R., & Zou, Y.-R. (2004). The role of CXCR4 in maintaining peripheral B cell compartments and humoral immunity. *The Journal of Experimental Medicine*, *200*(9), 1145–1156.
Norman, P. J., Norberg, S. J., Guethlein, L. A., Nemat-Gorgani, N., Royce, T., Wroblewski, E. E., Dunn, T., Mann, T., Alicata, C., Hollenbach, J. A., Chang, W., Shults Won, M., Gunderson, K. L., Abi-Rached, L., Ronaghi, M., & Parham, P. (2017). Sequences of 95 human MHC haplotypes reveal extreme coding variation in genes other than highly polymorphic HLA class I and II. *Genome Research*, *27*(5), 813–823.
Orekhov, A. N., Orekhova, V. A., Nikiforov, N. G., Myasoedova, V. A., Grechko, A. V., Romanenko, E. B., Zhang, D., & Chistiakov, D. A. (2019). Monocyte differentiation and macrophage polarization. *Vessel plus*, *3*, 10.
Rock, K. L., Reits, E., & Neefjes, J. (2016). Present Yourself! By MHC Class I and MHC Class II Molecules. *Trends in Immunology*, *37*(11), 724–737.
Shimizu, Y., & Dobashi, K. (2012). CC-chemokine CCL15 expression and possible implications for the pathogenesis of IgE-related severe asthma. *Mediators of Inflammation*, *2012*, 475253.
The Tabula Microcebus Consortium. (2021). Tabula Microcebus: A transcriptomic cell atlas of mouse lemur, an emerging primate model organism. In *bioRxiv* (p. 2021.12.12.469460). https://doi.org/10.1101/2021.12.12.469460

Todd, E. M., Zhou, J. Y., Szasz, T. P., Deady, L. E., D’Angelo, J. A., Cheung, M. D., Kim, A. H. J., & Morley, S. C. (2016). Alveolar macrophage development in mice requires L-plastin for cellular localization in alveoli. *Blood*, *128*(24), 2785–2796.

Tsukasaki, M., Huynh, N. C.-N., Okamoto, K., Muro, R., Terashima, A., Kurikawa, Y., Komatsu, N., Pluemsakunthai, W., Nitta, T., Abe, T., Kiyonari, H., Okamura, T., Sakai, M., Matsukawa, T., Matsumoto, M., Kobayashi, Y., Penninger, J. M., & Takayanagi, H. (2020). Stepwise cell fate decision pathways during osteoclastogenesis at single-cell resolution. *Nature Metabolism*, *2*(12), 1382–1390.

Ulvmar, M. H., Werth, K., Braun, A., Kelay, P., Hub, E., Eller, K., Chan, L., Lucas, B., Novitzky-Basso, I., Nakamura, K., Rülicke, T., Nibbs, R. J. B., Worbs, T., Förster, R., & Rot, A. (2014). The atypical chemokine receptor CCRL1 shapes functional CCL21 gradients in lymph nodes. *Nature Immunology*, *15*(7), 623–630.

Vermi, W., Riboldi, E., Wittamer, V., Gentili, F., Luini, W., Marrelli, S., Vecchi, A., Franssen, J.-D., Communi, D., Massardi, L., Sironi, M., Mantovani, A., Parmentier, M., Facchetti, F., & Sozzani, S. (2005). Role of ChemR23 in directing the migration of myeloid and plasmacytoid dendritic cells to lymphoid organs and inflamed skin. *The Journal of Experimental Medicine*, *201*(4), 509–515.

Wu, X., Hollingshead, N., Roberto, J., Knupp, A., Kenerson, H., Chen, A., Strickland, I., Horton, H., Yeung, R., Soysa, R., & Crispe, I. N. (2020). Human Liver Macrophage Subsets Defined by CD32. *Frontiers in Immunology*, *11*, 2108.
